# The genetic architecture of local adaptation is historically contingent

**DOI:** 10.64898/2026.02.01.703099

**Authors:** Tianlin Duan, Michael C. Whitlock, Tom R. Booker

## Abstract

Revealing the genetic basis of local adaptation is a common goal of evolutionary biology, but despite theoretical progress, general expectations for the genetic architecture of local adaptation are still unclear. Theoretical analyses usually model simplified ecologies or simplified genetic architectures of adaptive traits, so the interplay of these factors is missing from our understanding. In this study, we use simulations to explore how the interplay of ecological and genetic parameters influences the evolution and genetic architecture of local adaptation. With these simulations, we ask: i) What are the features of alleles that made the largest contribution to local adaptation, and how are they affected by polygenicity of adaptive traits, migration rates, demographic history, and the spatial pattern of the environment? And ii) does allele age moderate the confounding effect from population structure in genotype-environmental associations (GEA)? We find that the frequency, number, and phenotypic effect size of locally adaptive alleles are sensitive to trait polygenicity and demographic history, and that these factors shape the evolutionary dynamics of local adaptation. We find that population expansions can leave legacies in the genetic architecture of local adaptation, reducing the expected number of adaptive alleles relative to models with constant population size, and this effect is long-lasting. Compared to range expansion, other ecological variables known to affect the genetic basis of local adaptation had limited effects. Finally, allele age moderated the confounding effect of population structure and modified the causal effect of environmental variables on genotypes. Alleles that arose around the time of environmental changes often made large contributions to local adaptation, but young alleles often had the highest false positive rates and were the most common age category. We describe how incorporating allele age and its interactions with population structure and environmental variables may increase the sensitivity and specificity of GEA analysis. Overall, this work demonstrates the critical importance that a species’ demographic history can have on its genetic architecture of local adaptation.

## Introduction

Populations of a single species are often exposed to diverse environments, where heterogeneous selection may drive phenotypes toward different local optima. Such spatially heterogeneous selection can lead to local adaptation (Blanquart et al. 2013), a ubiquitous phenomenon in which local populations have higher fitness than non-local populations in their local environment (Hereford, 2009). Understanding the genetic basis of local adaptation is a common goal of evolutionary biology and is essential for applications in public health (Rees et al. 2020; Rochman et al. 2021), agriculture (Li et al. 2015; Takuno et al. 2015; Ndudzo et al. 2024), and forestry (Singh et al. 2024).

Revealing the genetic basis of local adaptation in nature requires some basic expectations on the genetic architecture of local adaptation. Assumptions about the number, effect size, frequency, and age of causal alleles of local adaptation are prerequisites for setting meaningful research goals and developing appropriate methods. Identifying locally adapted alleles often relies on genotype-environment association (GEA) studies, corroborated by outlier tests for genetic differentiation or other signals of population-specific selection (e.g., Evans et al., 2023; Lai et al., 2019; Leroy et al., 2020; Lovell et al., 2021; Razgour et al., 2019; Shi et al., 2023; Todesco et al., 2020; Whiting et al., 2024; Zou et al., 2024). These methods are naturally biased toward alleles with large effects and high minor-allele frequencies, especially when association studies have a limited sample size (Willing et al. 2012; Jakobsson et al. 2013; Lasky et al. 2023). When these methods are applied, researchers implicitly assume that a few large-effect/high-frequency alleles can explain a considerable amount of local adaptation and are a representative subset of locally adapted alleles; hence, recognizing and interpreting these alleles is of primary interest. Moreover, methods for detecting signatures of selection require assumptions on the age of locally adapted alleles, as distinct genetic signatures are expected for recent versus long-term balancing selection (Fijarczyk and Babik 2015; Jasper and Yeaman 2024), or selection on de novo mutations versus standing variation (Vitti et al. 2013). In summary, having an expected genetic architecture of local adaptation would help to determine whether assumptions about locally adaptive alleles are generally true, or under which conditions the assumptions hold.

In particular, knowing the expected age distribution of locally adapted alleles may also help to improve the performance of GEA. The age of alleles can be estimated with both mutation and recombination clocks using information from shared haplotype segments (Platt et al. 2019; Albers and McVean 2020). Once available, this information may be used in GEA for a more delicate control of confounding effects of population structure, which is still a major challenge in GEA (Lotterhos 2023). Due to geographically limited dispersal, younger alleles may form a cluster around the place of mutation origin. Their frequencies are more likely to be spatially autocorrelated and associated with environmental factors by chance. Therefore, we expect that allele age will be a moderator variable for the association between allele frequency and environmental factors for neutral alleles. Moreover, alleles which affect local adaptation experience balancing selection, which can cause them to persist longer than neutral alleles. Incorporating allele age into GEA methods would require a comprehensive understanding of the effects of allele age.

Most studies of genetic architecture have focused on global adaptation, in which a single population adapts to a single environmental optimum, rather than local adaptation. When assuming a population adapted to a new phenotypic optimum by stepwise fixations of new mutations as in Fisher’s geometric model (Fisher 1930), the phenotypic effect size of alleles that contribute to global adaptation is expected to have an exponential distribution (Orr 1998), with few large-effect adaptive alleles and many small-effect alleles. Notably, in this model, the relative effect size distribution of adaptive alleles is expected to be more or less independent of the raw distribution of phenotypic effects for new mutations before selection (Orr, 1998). Instead of the assumption of stepwise fixations, models of polygenic adaptation allow global adaptation to be achieved by simultaneous frequency changes at many loci. De Vladar and Barton (2014) analytically showed that during polygenic adaptation, alleles with an intermediate effect size that is close to a critical threshold determined by the mutation rate and the strength of selection are expected to drive the initial global adaptation. On the other hand, the long-term relative contribution from small- and large-effect alleles strongly depends on the size distribution of raw mutations before selection (De Vladar and Barton 2014; Jain and Stephan 2017). When small-effect alleles are abundant and contribute the bulk of genetic variance, large-effect alleles are unlikely to contribute to long-term global adaptation unless the shift in optimum is very large and there is an intermediate mutational input of large-effect mutations (Milligan et al., 2025).

Local adaptation can be approximated as an aggregation of multiple cases of global adaptation with heterogeneous environmental optima connected by migration, but the expected genetic architecture of local adaptation can dramatically differ from that of global adaptation. In contrast to Orr’s (1998) predictions for global adaptation, large-effect alleles are expected to be more resistant to the homogenizing effect of migration (Yeaman and Otto 2011), have a larger probability to be adaptive (Yeaman and Whitlock 2011), and eventually make a greater contribution to phenotypic differentiation between populations (Griswold 2006).

Despite the progress in theoretical studies, a general expectation for the genetic architecture of local adaptation is still not clear for at least three reasons. First, traits with a polygenic and redundant genetic architecture are likely common in nature (Shi et al. 2016; Walsh and Lynch 2018; Sella and Barton 2019; Láruson et al. 2020; Yengo et al. 2022), and may lead to distinct patterns in global adaptation (Jain and Stephan 2017) or local adaptation (Yeaman 2015; Yeaman 2022). Second, it is not clear how sensitive the genetic architecture of local adaptation is to genetic and ecological parameters. Simulations showed that the genetic architecture of complex traits under global adaptation is sensitive to demography and the raw effect size of mutations before selection (Lohmueller 2014; Stetter et al. 2018). This raises the question of whether these parameters can also affect the genetic architecture of local adaptation. Third, with a high genetic redundancy, small-effect alleles that contribute to local adaptation can be continuously replaced (Yeaman 2022), but the temporal change of genetic architecture itself has not been well summarized, e.g., whether the relative contributions from alleles with different effect sizes and frequencies change in different temporal phases of local adaptation. An analytical study of global adaptation showed that after a sudden change in optimal phenotype, the initial contribution of large-effect alleles can be almost completely replaced by the contribution of alleles with smaller effects in the long term (Hayward and Sella 2022). It is unknown whether this temporal change of genetic architecture also exists in local adaptation, in which large-effect alleles are expected to make larger contributions (Griswold 2006; Yeaman and Whitlock 2011).

To remain tractable, analytical studies on the genetic architecture of local adaptation often have to greatly simplify their models by assuming linkage equilibrium, a simple population structure (e.g., a two-patch model and panmictic populations), equilibrium demography, or a small number of mutations, which often have fixed fitness effects, a particular value of frequency, or equal phenotypic effect sizes. Another common assumption is that adaptive alleles all arise from *de novo* mutation or are sourced entirely from genetic variation present before the onset of heterogeneous selection. While these studies provided a sensible starting point and basic principles, they may also exaggerate cases that are less representative in nature, distort the magnitude of parameter effects, or overlook phenomena that arise from complex interactions. In the present study, we explore the genetic architecture of local adaptation using individual-based simulations modelling typical evolutionary scenarios with parameters informed by empirical estimates. Specifically, we first characterize the alleles that made the largest contribution to local adaptation, which could be the desired target of future empirical inquiries. To this end, we measure the total contribution to local adaptation from alleles with different phenotypic effect sizes, frequencies, and ages, and how these distributions are affected by genetic and ecological parameters, including levels of polygenicity, migration levels, demographic history, autocorrelation of environmental maps, and temporal phases of local adaptation. Second, as we are interested in the potential use of allele age in GEA, we test the moderating effect of allele age on the confounding effect of population structure in some typical demographic scenarios that are both common and notorious for confounding GEA analyses. We end by discussing the implications of the depicted genetic architecture of local adaptation on GEA studies.

## Methods

### Individual-based simulations

We conducted individual-based simulations to explore the genetic architecture of local adaptation. Simulations were implemented in SLiM (v4.3; Haller and Messer, 2023), and simulation scripts are available at https://github.com/euptelia/arg-for-gea. In brief, we simulated heterogeneous stabilizing selection with varying phenotypic optima on a single quantitative trait among individuals in continuous space, using a non-Wright-Fisher model. Values of important parameters of simulations are summarized in Table 1.

**Table 1.** Notations and descriptions of parameters of simulations.

| Symbol | Description | Values shared | Values specific to models with |  |  |  |
| --- | --- | --- | --- | --- | --- | --- |
|  |  |  | low migration | high migration | low polygenicity | high polygenicity |
| $C_{max}$ | Maximum strength of competition | 1.0 | - | - | - | - |
| $D_{comp}$ | Maximum distance of competition | 0.03 | - | - | - | - |
| $D_{mate}$ | Maximum distance of mate choice | - | 0.12 | 0.15 | - | - |
| $h$ | Dominance coefficient of functional mutations | 0.5 | - | - | - | - |
| $K$ | Carrying capacity density | 6000 (with selection*) or 5000 (no selection) | - | - | - | - |
| $R$ | Reproduction rate | 0.2 | - | - | - | - |
| $r$ | Recombination rate along a chromosome | 1e-7 | - | - | - | - |
| $\mu_n$ | Mutation rate of neutral mutations | 5e-8 | - | - | - | - |
| $\mu_f$ | Mutation rate of functional mutations | - | - | - | 1e-10 | 1e-8 |
| $\sigma_C$ | Standard deviation of the Gaussian competition function | 0.01 | - | - | - | - |
| $\sigma_D$ | Standard deviation of the distribution of dispersal distance | - | 0.03 | 0.06 | - | - |
| $\sigma_M$ | Standard deviation of the distribution of the phenotypic effect size of functional mutations | - | - | - | 0.1 | 0.01 |
| $\sigma_W$ | Standard deviation of the Gaussian fitness function | 0.4 | - | - | - | - |

\*Corresponding to 5357±127 (mean±SD) diploid individuals in the present simulations

Individuals occupied a continuous spatial range. We specified a single carrying capacity (6000) for the entire range that resulted in 5357±127 (mean±SD) hermaphroditic diploid individuals for models with selection at the main sampling stage. The simulated range was a two-dimensional square space, with boundaries of [0, 1] in both dimensions. In each simulated year, individuals were able to mate within a radius of *D*_*mate*_ (the maximum mating distance) of their location and self-fertilization was strictly prevented. The number of offspring generated by a given individual was drawn from a Poisson distribution with a mean of 0.2 (*R*), so the population rarely successfully reproduced. The population was allowed to have overlapping generations, although the probabilities of survival and reproduction were not age-dependent. After reproduction, parental individuals were not automatically removed but competed with their newborn offspring. The survival of all individuals was evaluated together under the same selection and density-dependent regulation, and applied before the start of the next simulated year. In all models, the median age of simulated individuals was 4 simulated years, and the 95th percentile was 22-26 simulated years (Table S1). Offspring dispersed away from maternal parents, assuming a 2-dimensional Gaussian dispersal kernel (mean of 0 and a radius of *σ*_*D*_).

Our simulations were genetically explicit. We modelled a 10-megabasepair (Mb) diploid genome divided into four equally sized chromosomes. The first three chromosomes contained 5-kbp-long “gene regions”(representing genes and flanking regions), separated by 10-kbp “neutral regions”. The fourth chromosome only contained neutral regions.

Individuals possessed a single quantitative trait, which was computed as the sum of its functional mutational effects. Functional mutations, those that could affect the quantitative trait, could occur at any position in gene regions with a mutation rate of *μ*_*f*_, so the “trait” had a broad mutational target size (2.5 Mb). The phenotypic effect size of functional mutations (*α*) was drawn from a Gaussian distribution, with a standard deviation of *σ*_*M*_ and a mean of 0. We modelled two levels of polygenicity: 1) high polygenicity, with many mutations and smaller phenotypic effect sizes (*σ*_*M*_ = 0.01, *μ*_*f*_ = 1 × 10^−8^ per base pair per generation); and 2) low polygenicity, with fewer mutations, but mutations with larger effects were more common (*σ*_*M*_ = 0.1, *μ*_*f*_ = 1 × 10^−10^). Note that the mutational variance contributed to the trait is identical in these two cases.

#### Selection

The single quantitative trait was under stabilizing selection on survivorship, with phenotypic optima varying across the simulated range. The relative survival of an individual with phenotype *z* was the probability density of a normal distribution with a standard deviation of 0.4 (*σ*_*W*_) and a mean of local phenotypic optima (*z*^∗^), rescaled by the maximum value of the function at *z*^∗^:

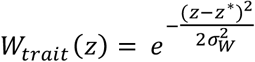

We modelled spatial heterogeneity in phenotypic optima using two different environmental maps: In the “cline map”, local phenotypic optima changed linearly in the *y*-dimension between 0 and 1, but was invariable along the *x*-dimension, forming an environmental cline along the *y*-dimension (Figure 1); We also generated a “patchy map” (Figure 1) by randomly drawing 100 values from a uniform distribution over [0, 1], which were used as the corners of 9×9 square patches. Interpolated values were then calculated for locations among the corners of these patches in SLiM (*patchyMap*.slim).

**Figure 1.**
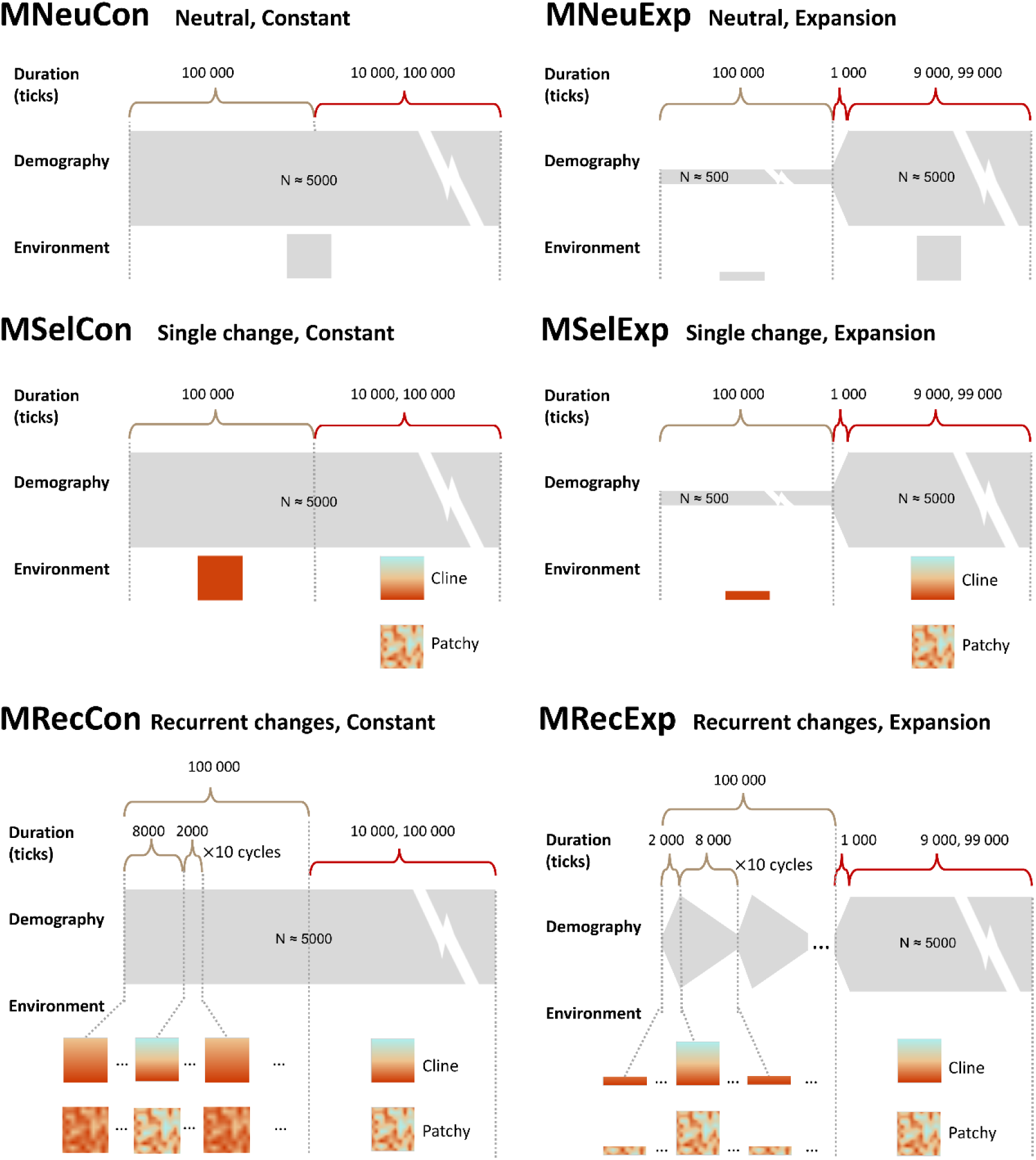
Models of demography and modes of selection in SLiM simulations. Observed census population sizes (*N*) were indicated by the numbers on the grey diagrams of demographic models. The chromatic colors on environmental maps represent the value of phenotypic optima (*z*^∗^) of Gaussian stabilizing selection on the trait, with red=0 and cyan=1, while grey means no selection on the trait. The size of the environmental map shows the area of available habitats. In models MRecCon and MRecExp, the change of phenotypic optima and area of habitats was continuous and linear in time. The duration of burn-in and sampling stages was indicated by brown and red curly brackets, respectively. For single-sample simulations, tree sequences were saved at simulated year (tick) 10,000 of the sampling stage. For time-series sampling, 16 samples were taken between 200 and 100,000 simulated years of the sampling stage.

In addition to selection on the quantitative trait, individuals were also subject to density-dependent regulation due to spatial competition in all cases. The total strength of competition on a focal individual (*I*(*x*, *y*)) from *q* neighbor individuals within the maximum interaction distance (*D*_*comp*_) was the product of the probability density of two normal distributions, in *x*-and *y*-dimensions:

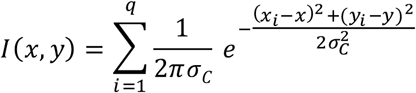

where (*x*, *y*) and (*x*_*i*_, *y*_*i*_) were the spatial coordinates of focal and neighbor individuals, respectively, and both normal distributions have a standard deviation of *σ*_*C*_. The total competition strength was calculated with the SLiM function totalOfNeighborStrengths(). Then the fitness effect of competition was defined as the carrying capacity density (*K*) divided by a rescaled competition strength:

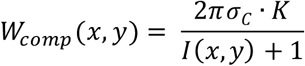

Finally, the overall fitness of an individual with phenotype *z* and location (*x*, *y*) was:

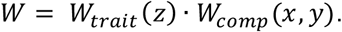

#### Migration

The level of migration was controlled by altering the maximum distance of mate choice (*D*_*mate*_) and the standard deviation of the dispersal kernel (*σ*_*D*_). We chose two migration levels that resulted in different levels of overall population differentiation. The overall level of population differentiation was measured by Weir and Cockerham’s *F*_*ST*_ (Weir and Cockerham 1984), as implemented in the Python package scikit-allel 1.3.13 (Miles et al. 2024). We analyzed the simulated area by dividing the continuous space into a 10×10 grid and calculated the overall *F*_*ST*_ among these 100 subpopulations using all individuals.

At the sampling stage, the average *F*_*ST*_ of models with high and low migration levels were 0.038±0.002 and 0.112±0.008 (mean±SD), respectively. The two average *F*_*ST*_ values are comparable with levels of differentiation in tree species with wide geographic ranges

(Candido-Ribeiro and Aitken 2024; Lazic et al. 2024).

### Demography and temporal modes of selection

We aimed to model the genetic architecture of local adaptation in some typical evolutionary scenarios, so we constructed a series of models with different demographic histories and modes of selection (Figure 1 and Table 2).

**Table 2.** Models of demographic history and modes of selection.

|  |  | Demography |  | 930 |
| --- | --- | --- | --- | --- |
|  |  | a. Constant | b. Expansion | 931 |
| No selection on the trait |  | MNeuCon: Constant population size and no selection on the quantitative trait | MNeuExp: Single population expansion and no selection on the quantitative trait |  |
| With heterogeneous stabilizing selection on the trait | Single change | MSelCon: Single environmental change | MSelExp: Single population expansion along with environmental change |  |
|  | Recurrent changes | MRecCon: Recurrent environmental changes | MRecExp: Recurrent population expansions and contractions along with environmental changes |  |

Local adaptation is often accompanied by population expansion, which may alter the genetic architecture of local adaptation as it does in global adaptation (Stetter et al. 2018), and is also a common confounder for GEA (Hoban et al. 2016; Nadeau et al. 2016; Capblancq et al. 2023). To reflect this common scenario, we modelled populations with a constant population size, a single population expansion, or ten cycles of recurrent population expansion and contraction (“glacial cycles”). In each glacial cycle, the population was allowed to expand into the entire landscape over 2,000 simulated years, after which time the landscape contracted for a period of 8,000 years. For both population expansion and contraction, we changed the area of available habitats linearly in time (Figure 1).

We modelled temporal variation in the pattern of selection. First, we modelled a scenario with a single environmental change from homogeneous selection to spatially heterogeneous selection (a single round of local adaptation), and a case with recurrent environmental changes (recurrent local adaptation to similar environmental variation). In the case of a single environmental change, phenotypic optima gradually changed from a single value (0) to one of the heterogeneous environmental maps. For recurrent environmental changes, the range of values on maps gradually changed from [0, 0.5] to [0, 1] in 2,000 ticks, and changed back to [0, 0.5] in 8,000 ticks.

In all models, simulations were run for a burn-in of 100,000 simulated years, and the most recent environmental change started at the end of the burn-in stages (time 0 of sampling stages). Tree sequences were recorded during the SLiM simulations (Haller et al. 2019) and were saved 10,000 years after the most recent environmental change at the sampling stage. We made 200 replicates of this single-sampling simulation for each parameter set. To explore changes in genetic architecture over time, we also ran 30 replicates of longer simulations with time-series sampling. During each simulation, we recorded the state of the population during 200-100,000 simulated years after the most recent environmental change, with denser sampling at earlier stages in which the genetic architecture changed more rapidly. For each time-series sampling simulation, 16 tree sequences were saved in total at 200, 400, 600, 800, 1,000, 2,000, 4,000, 6,000, 8,000, 10,000, 20,000, 30,000, 40,000, 60,000, 80,000, and 100,000 simulated years after the beginning of the sampling stage.

### Analyzing simulated data

After SLiM simulations, the recorded tree sequences were analyzed with Python packages tskit 0.6.0 (Ralph et al. 2020), msprime 1.3.3 (Baumdicker et al. 2022), SciPy 1.13.1 (Virtanen et al. 2020), NumPy 2.2.2 (Harris et al. 2020), and pandas 2.2.3 (McKinney 2010). We extracted the frequency, age, phenotypic effect size, and spatial location of each allele from tree sequences. Results were visualized using the Python package Matplotlib 3.9.2 (Hunter 2007).

#### Measuring local adaptation and allelic contribution

We measured the overall level of local adaptation of the whole population with a statistic 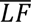 (Table 3), modified from the conventional *Mean[LF]*, the average mean fitness difference between local and foreign individuals that are transplanted to local patches (Blanquart et al. 2013). We defined “local patches” as circles centered at the locations of every existing individual at the sampling time, with a radius equal to the maximum distance of mate choice. For each local patch, we virtually “transplanted” all individuals that were outside of the patch (foreign individuals) to the center of the local patch, and re-calculated their fitness after this transplanting, using their original phenotypes and the local phenotypic optima. Then we calculated the difference between the mean fitness of local and foreign individuals and averaged the fitness differences among all patches.

**Table 3.** Notations and descriptions of variables and statistics.

| Symbol | Description |
| --- | --- |
| $K_{80,pos}$ | The minimum number of alleles needed to explain 80% of the total positive allelic contribution to local adaptation |
| $K_{80,net}$ | The minimum number of alleles needed to explain 80% of the total net allelic contribution to local adaptation |
| $\overline{LF}$ | The average mean fitness difference between local and foreign individuals of all subpopulations |
| $LF_{mut}$ | The contribution from a single allele to local adaptation, calculated as the difference between the original $\overline{LF}$ and recalculated $\overline{LF}$ after randomly shuffling the geographic location of all copies of the allele in the population |

To measure the contribution of each functional mutation to local adaptation, we defined another statistic *LF*_*mut*_. For each derived allele with a non-zero phenotypic effect size, we randomized its placement among all existing individuals in space, keeping the frequency of the focal allele and the placement of all other alleles the same as before. After shuffling the locations of this single allele, we calculated new phenotypes of all individuals and recalculated 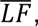 termed 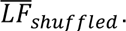 The contribution to local adaptation from the focal allele *i* is measured by: 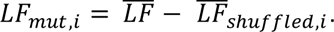 A relative *LF*_*mut*, *i*_ is *LF*_*mut*, *i*_ divided by the sum of *LF*_*mut*_ in that simulation replicate.

#### Measuring the change of genetic architecture

We calculated two statistics for monitoring the change of genetic architecture: *K*_80,*pos*_ and *K*_80,*net*_. They are the minimum number of alleles for explaining 80% of the total positive or net allelic contribution to local adaptation, respectively.

After the level of local adaptation reached a plateau, a further change of *K*_80,*net*_ will reflect a change of contribution from individual alleles, while the difference between the slopes of *K*_80,*pos*_ and *K*_80,*net*_ will reflect the contribution from compensatory evolution (the symmetrical increase of effects from both adaptive and maladaptive alleles) to the change of genetic architecture. We used a linear model with an interaction term to test whether the slope of *K*_80_on time is different between the two types of *K*_80_ during 1,000-10,000 simulated years after the last environmental change, in which *K*_80,*pos*_ largely increased in models with population expansions.

#### Genotype-environment association analysis on simulated data

To test the relationship between allele age and the performance of GEA, neutral mutations were simulated and superimposed on the recorded tree sequences at a rate of 5 × 10^−8^ (per base pair per generation) for the entire genome, using msprime.

We tested the ordinal association between allele frequency and environment with simulated data. The square map was divided into a 10×10 grid, and all individuals in each of the 100 small squares were used to calculate an allele frequency for that location. Then we performed Kendall’s rank correlation test between allele frequency of subpopulations and phenotypic optima.

We compared the false positive rates (FPR=False positives/(False positives+True negatives)) of neutral alleles in 40 allele age intervals. For neutral alleles, false positives were defined as alleles that showed a significant correlation with environmental factors (*p* < 1 × 10^−10^ in Kendall’s rank correlation test). Alleles with a minor allele frequency < 0.01 were excluded from the analyses, as is often the case for empirical studies.

## Results

### Local adaptation emerged under different parameter sets, with highly variable genetic architectures

Local adaptation emerged under different parameter sets with highly variable genetic architectures. For all parameters tested, local adaptation evolved in the 10,000 simulated years (ticks) following the most recent environmental change. In all cases, the average mean fitness difference between local and foreign individuals 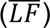 was significantly higher than zero in all models with heterogeneous selection on the quantitative trait (One-sample two-sided *t*-test, *p* < 0.001 in all 32 tests; Table S2). However, the level of local adaptation differed among models. The average 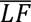 in models with a patchy environmental map was 0.032±0.002 (mean±SD), which was only one-sixth of that in a cline map (0.186±0.003; Welch’s *t*-test, *p* < 0.001), and a higher level of migration slightly reduced 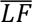 (Low migration: 0.111±0.077; High migration: 0.107±0.077; Welch’s *t*-test, *p*=0.026; Table S2). The genetic architectures underlying local adaptation in the simulations were highly variable, with phenotypic effect sizes, allele frequencies, and the temporal dynamics of alleles contributing to local adaptation varying depending on simulation parameters (Figures S1-S3). The most striking effect on the architecture of adaptation was found between models that differed in the polygenicity of the trait under selection. We contrasted a model with a high functional mutation rate with small phenotypic effects of new mutations to a model with a lower functional mutation rate with large effect sizes (low polygenicity). Note that the mutational variance for the quantitative trait was constant between the high and low polygenicity models. As expected, the low polygenicity model led to local adaptation by alleles with large phenotypic effects (Figures 2B and S2). In these cases, alleles with low minor allele frequencies (<0.1) had almost no contribution to local adaptation (Figures 2D and S2). The adaptive alleles mostly arose during the time around the first or the last environmental change and were then maintained in the population until the end of the simulation (Figures 2F and S3). Finally, in models with a patchy environment a very small fraction of these older alleles were replaced by young ones over time (Figure S3).

**Figure 2.**
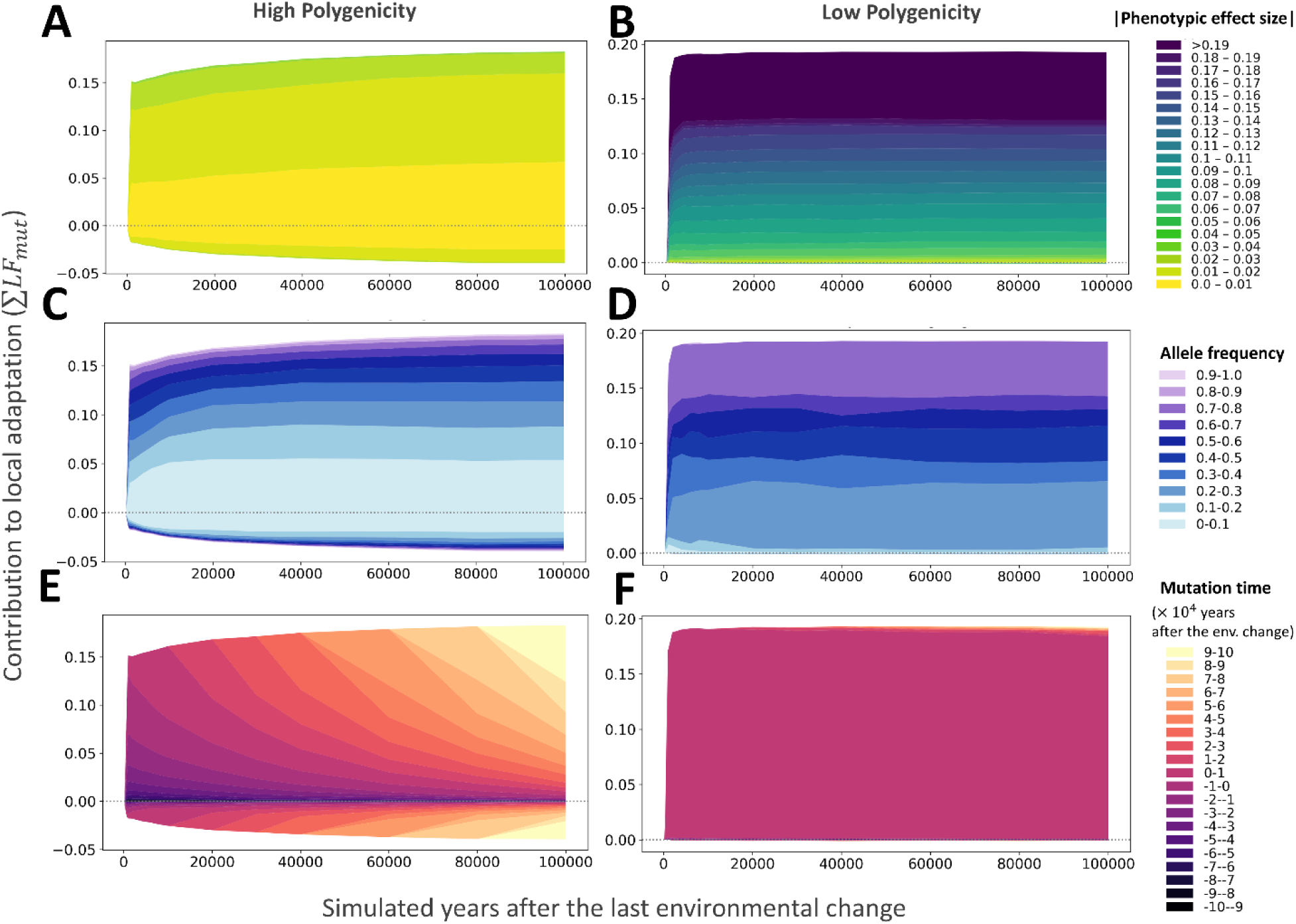
The contribution to local adaptation from alleles with different phenotypic effect sizes (A and B), allele frequencies (C and D), and mutation times (E and F) 0-100,000 simulated years (ticks) after the last environmental change in a model with recurrent population expansions and contractions (MRecExp), a cline environmental map, and a high migration level. The average values of 30 SLiM simulation replicates are displayed in each plot. High polygenicity: The standard deviation of the distribution of phenotypic effect size *σ*_*M*_ = 0.01, and functional mutation rate *μ*_*f*_ = 1 × 10^−8^ per base pair per simulated year; Low polygenicity: *σ*_*M*_ = 0.1, *μ*_*f*_ = 1 × 10^−10^.

In a model with high polygenicity, locally adaptive alleles had smaller mean phenotypic effects, as expected, but there was a greater fraction of local adaptation that came from alleles at low frequencies (<0.1) (Figures 2A, 2C, S1 and S2). In addition, there was a slow but continuous turnover of adaptive alleles, where most old alleles were gradually replaced by younger mutations (Figures 2E and S3).

### The genetic architecture of local adaptation was mainly shaped by the polygenicity of traits and demographic history

An outstanding empirical question is, how many causal alleles underlie patterns of local adaptation. Depending on the answer, methods focusing on detecting outliers or analyzing the variance of important traits may be preferred. We explored how the architecture of local adaptation was influenced by genetic and ecological factors using a statistic, *K*_80,*pos*_, which is the minimum number of alleles needed to explain 80% of the total positive allelic contribution to local adaptation. We first compared *K*_80,*pos*_ among samples taken 10,000 simulated years after the last environmental change.

Among our parameter sets, we found that *K*_80,*pos*_ was primarily determined by the polygenicity of the trait. In simulations with high and low polygenicity, the average *K*_80,*pos*_ was 190.2±46.1 and 2.9±1.2 (mean±SD), respectively. In simulations of high polygenicity, demographic history also had large effects on *K*_80,*pos*_. Compared to models with a constant population size, a demographic history consisting of a single environmental change with a single population expansion reduced *K*_80_ by approximately 42% (Welch’s *t*-test, *p* < 0.001), while recurrent cycles of environmental change with population expansion/contractions reduced it by 30% (Welch’s *t*-test, *p* < 0.001; Figure 3). The migration rate variation we implemented did not have a substantial effect on *K*_80_. The higher migration level reduced *K*_80,*pos*_ by about 6% in simulations of high polygenicity (Welch’s *t*-test, *p* < 0.001; Figure 3). Finally, the autocorrelation of environmental maps (cline/patchy) only had trivial effects. The average *K*_80,*pos*_ of simulations with cline maps was 4% lower than that with patchy maps (Welch’s *t*-test, *p* < 0.001; Figure 3).

**Figure 3.**
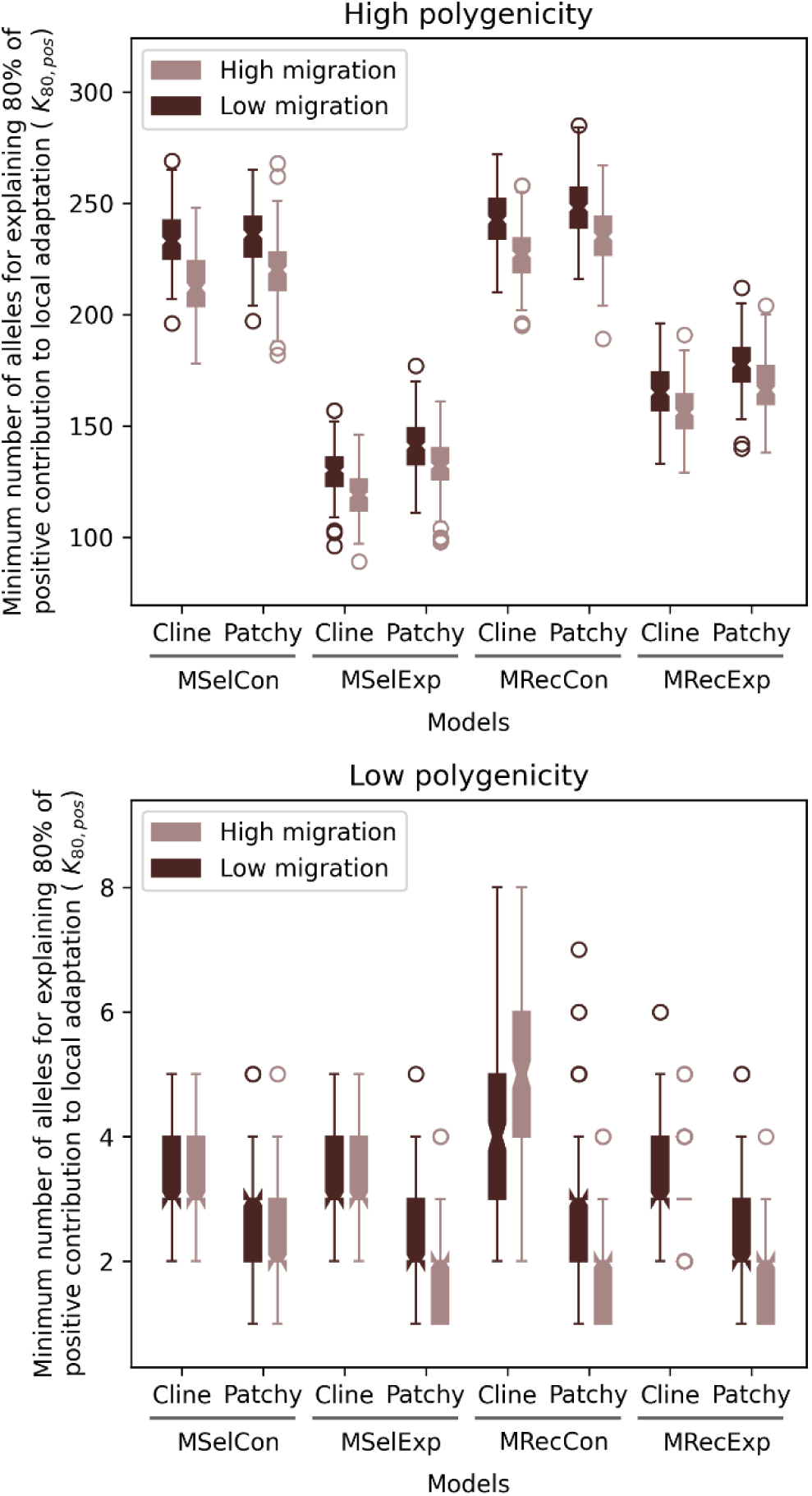
The minimum number of alleles needed to explain 80% of the total positive contribution to local adaptation (*K*_80_(*positive*)) 10,000 simulated years (ticks) after the last environmental change from 200 SLiM simulation replicates. The upper plot shows simulations with a high polygenicity (standard deviation of the distribution of phenotypic effect size *σ*_*M*_ = 0.01, and functional mutation rate *μ*_*f*_ = 1 × 10^−8^ per base pair per simulated year). The lower plot shows simulations with a low polygenicity (*σ*_*M*_ = 0.1, *μ*_*f*_ = 1 × 10^−10^). Migration levels were indicated by the color of boxes, with a darker color representing a lower migration level. The high and low migration levels resulted in average overall *F*_*ST*_ values of 0.038±0.002 and 0.112±0.008 (mean±SD), respectively. MSelCon: a single environmental change with a constant population size; MSelExp: a single environmental change with a population expansion; MRecCon: recurrent environmental changes with a constant population size; MRecExp: recurrent environmental changes with recurrent population expansions and contractions. “Cline" and "Patchy" are the types of environmental maps.

### The genetic architecture of local adaptation reached equilibrium slowly

The genetic architecture of local adaptation with population expansion took a long time to reach an equilibrium. In particular, we found that *K*_80,*pos*_ in models with population expansion and a high polygenicity kept increasing even after the level of local adaptation reached a plateau, and took a long time to reach a similar level as *K*_80,*pos*_ in the corresponding models with a constant range size (Figure 4B).

**Figure 4.**
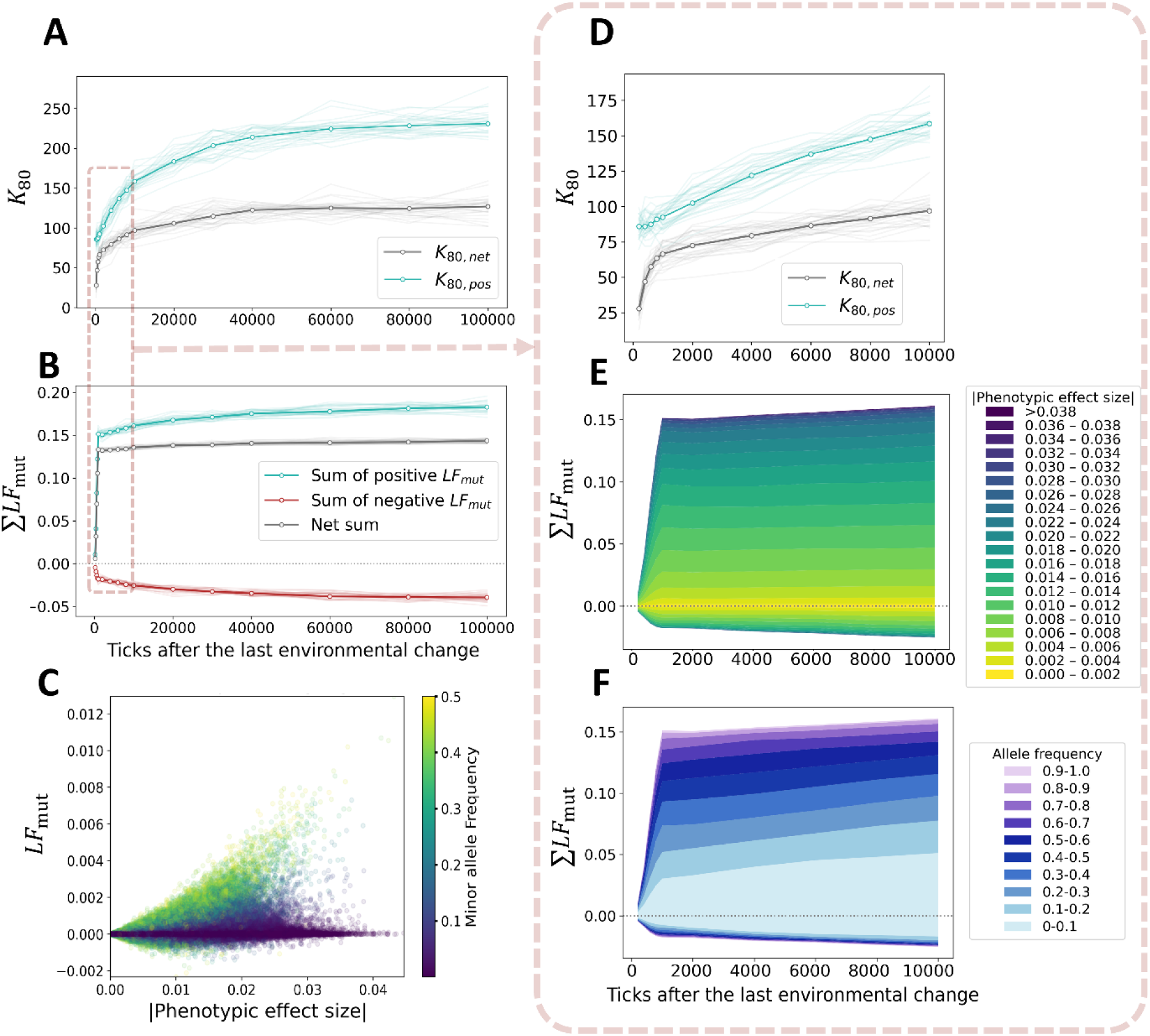
Temporal changes of the genetic architecture of local adaptation in a model with recurrent population expansions and contractions, a high polygenicity, a cline environmental map, and a high migration level. (A) The change of the minimum number of alleles for explaining 80% of the sum of positive allelic contributions to local adaptation (*K*_80,pos_) or net total contribution to local adaptation (*K*_80,net_) after the most recent environmental change. (B) The sum of total positive, negative, and net allelic contribution to local adaptation (∑*LF*_*mut*_). (C) The relationship among allelic frequency, absolute phenotypic effect size, and individual contribution to local adaptation (*LF*_*mut*_) 10,000 simulated years (ticks) after the environmental change. (D) *K*_80_ during 0-10,000 simulated years after the last environmental change, in which *K*_80_ changed rapidly. (E) The contribution to local adaptation from alleles with different phenotypic effect sizes. (F) The contribution to local adaptation from alleles with different frequencies. Results were aggregated from 30 simulation replicates, except for plot (C), which was from 200 replicates.

In all models with population expansion and high polygenicity, local adaptation was always established quickly, within the first 1000 simulated years following environmental change, and the level of local adaptation only had negligible changes afterward (Figures 4B and S4). However, the minimum number of alleles for explaining 80% of the total positive contribution to local adaptation did not stop increasing until 40,000 simulated years after the environmental change (Figures 4A and S5). The change of *K*_80,*pos*_ was most striking in models with a single population expansion, in which the increase did not stop until the end of simulations (100,000 simulated years after the most recent environmental change, Figure S5). By the end of the simulation, *K*_80,*pos*_ was roughly four times as high as that at the time when local adaptation was first established, and close to the *K*_80,*pos*_ of the corresponding model without population expansion (Figure S5).

In contrast, when a population had a constant population size, the number of needed alleles for explaining 80% of the total positive contribution to local adaptation only mildly fluctuated after the most recent environmental changes (Figure S5).

### Both compensatory evolution and allele-frequency changes contributed to the temporal change of genetic architecture

In models of high polygenicity, although the overall level of local adaptation quickly reached a plateau, the total positive contribution to local adaptation (sum of positive *LF*_*mut*_) could keep increasing, and was cancelled out by a symmetric increase of total negative contribution from maladaptive alleles (sum of negative *LF*_*mut*_, Figure 4B). The steady increase in *K*_80,*pos*_ overtime in models of local adaptation and population expansion may simply reflect the increase in the sum of positive *LF*_*mut*_, or alternatively, due to a decrease in the absolute contribution from each allele. To distinguish these two effects, we calculated another statistic *K*_80,*net*_, the minimum number of alleles for explaining 80% of the total *net* allelic contribution to local adaptation, which is the net sum of both positive and negative effects. The net sum of *LF*_*mut*_ plateaued 1000 simulated years after the last environmental change (Figures 4B and S4), so the change of *K*_80,*net*_ after 1000 simulated years should reflect the change of individual allelic contributions. The difference between the slopes of *K*_80,*net*_ and *K*_80,*pos*_ would reflect an extra increase of *K*_80,*pos*_ due to compensatory evolution, the symmetric increase of adaptive and maladaptive alleles.

Both the increase of total positive *LF*_*mut*_ and the decrease of individual allelic contribution led to the increase of *K*_80,*pos*_ after population expansion. In all models of population expansion, *K*_80,*net*_ significantly increased between 1,000 and 10,000 simulated years (Table S3). For example, in the model with recurrent population expansions and contractions, high migration rates, and the cline map, the median *K*_80,*net*_ increased from 67 to 97 between 1,000 and 10,000 simulated years, while the total net allelic contribution only increased by 1% during this time (Figure 4). This shows that the contributions of each locally adaptive allele decreased over time, which partly explained the increase of *K*_80,*pos*_. On the other hand, we also found a significant interaction between time and the type of *K*_80_ during 1,000 and 10,000 simulated years in all models of expansion (Table S3). The slope of *K*_80,*pos*_ was two to four times the slope of *K*_80,*net*_ (Table S3), suggesting that compensatory evolution, the increase of new adaptive alleles that compensated for new maladaptive alleles, also largely contributed to the increase of *K*_80,*pos*_.

The decrease in contribution from individual alleles may be further attributed to the shift of phenotypic effect size, allele frequency, or the geographic distribution of locally adapted alleles, as the allelic contribution to local adaptation depended on both phenotypic effect size and current frequency (Figure 4C). Overall, during the whole sampling period (200-100,000 simulated years after the last environmental change), the relative contributions from alleles of different sizes remained stable, though the contribution from large-effect alleles slowly decreased in models of a single population expansion (Figure S1).

However, during the period in which *K*_80,*net*_ of all expansion models experienced a fast increase (1000-10 000 simulated years after the last environmental change), the contribution from alleles of different sizes did not show an obvious change (Figures 4E and S1). In contrast, during this time, the contribution from low-frequency alleles more obviously increased (Figures 4F and S2). Therefore, the shift of allele frequency was more likely responsible for the decrease in the contribution from individual alleles.

### Alleles arising around the time of environmental change may make a large contribution to local adaptation

Is new mutation or preexisting standing variation more important for local adaptation? A continuous counterpart to this binary question would be: What is the age distribution of the alleles that make the largest contributions to local adaptation? We explored this question with our diverse models, using samples that were taken 10,000 simulated years after the most recent environmental change.

To jointly examine allele counts and individual contributions of locally adapted alleles, we compared the total allelic contribution to local adaptation of alleles across different ages. We found that the relationship between the relative contribution to local adaptation and allele age varied between two extremes (Figure S6). One extreme occurred when a model had a combination of high polygenicity for the trait, a constant population size, a low migration level, and recurrent local adaptation to the same environmental factor. With this parameter combination, the contribution to local adaptation across allele ages had a smooth, approximately geometric distribution (Figure 5). Any violation of these conditions led to a greater contribution from alleles that arose around the time of environmental change, though to different degrees. A single population expansion or recurrent expansions/contractions could drastically increase the contribution around the time of environmental change (Figure 5A), while a higher migration level mildly increased the peak (Figure 5A and 5C). It is worth noting that the alleles that made up these peaks tended to have a larger phenotypic effect size (Figure 5).

**Figure 5.**
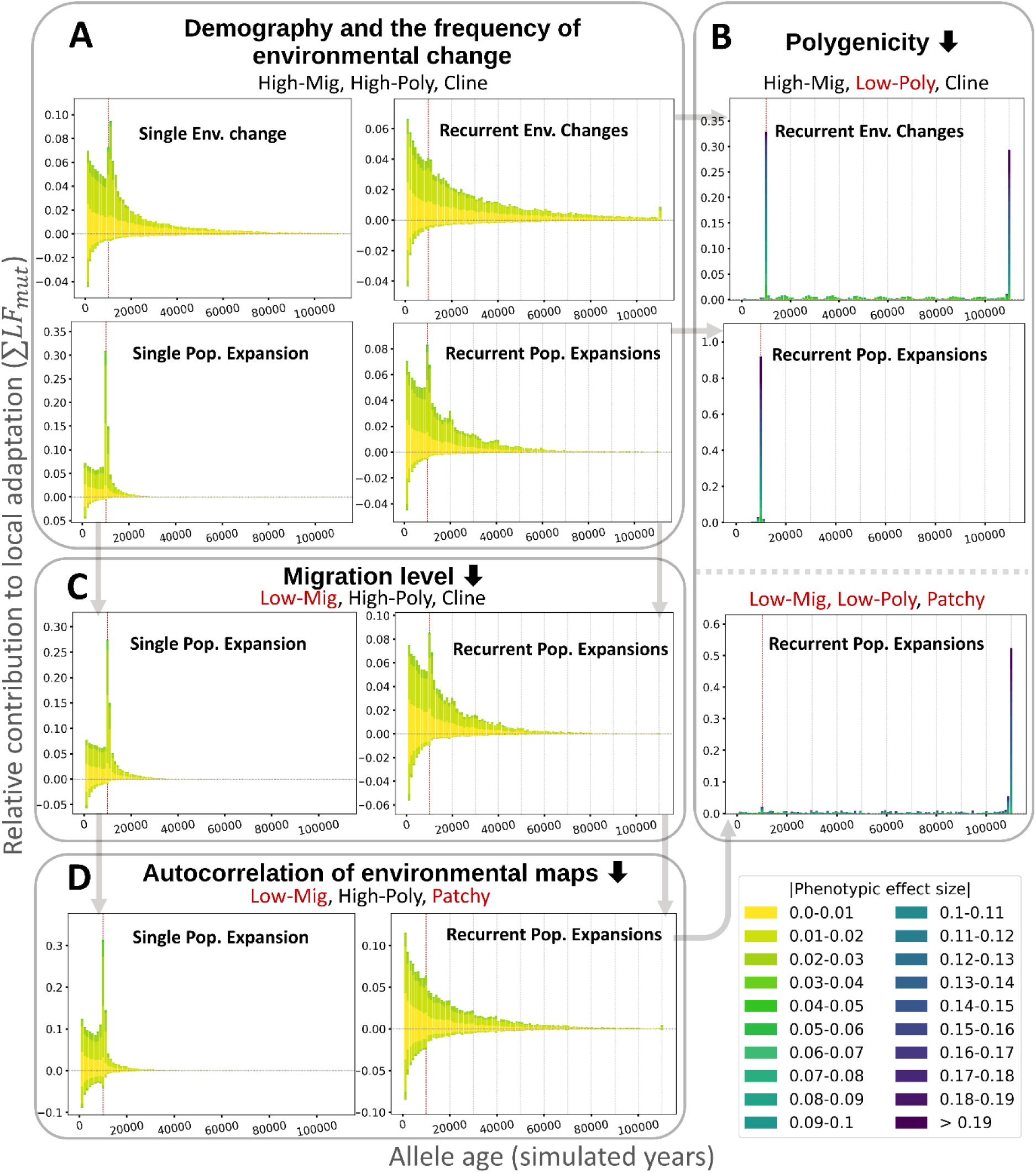
The relative contribution from alleles arising at different times to local adaptation was affected by multiple genetic and ecological parameters. (A) Effects of demographic history and the frequency of environmental change. (B) Effects of a lower level of polygenicity. (C) Effects of a lower migration level. (D) Effects of the autocorrelation of environmental maps (Cline/Patchy). Plots connected by grey arrows only differ by one parameter. In all panels, samples were taken at 10,000 simulated years after the most recent environmental change, with 200 SLiM simulation replicates. The red dashed vertical lines indicate the time of the (last) environmental change. For models with recurrent environmental changes or recurrent population expansions, grey dashed vertical lines indicate the starting times of the past ten rounds of environmental change. Contributions from alleles of different phenotypic effect size categories are distinguished by color. Env.: environmental; Pop.: population; Low-Mig./High-Mig.: a lower/higher migration level (corresponding to average overall *F*_*ST*_ values of 0.112±0.008 and 0.038±0.002, respectively); Low-Poly/High-Poly.: a lower or a higher level of polygenicity (The standard deviation of the distribution of phenotypic effect size *σ*_*M*_ = 0.1 and 0.01, functional mutation rate *μ*_*f*_ = 1 × 10^−10^ and 1 × 10^−8^ per base pair per simulated year, respectively).

Models with a low polygenicity represent the other extreme. In all cases, most local adaptation was due to alleles that arose around the time of environmental changes (Figure 5B). When there was only a single environmental change, almost all contributions were from alleles that arose around the time of that environmental change. When a population went through recurrent environmental changes or recurrent population expansions, the age of locally adaptive alleles depended on factors that affect the capacity of retaining adaptive alleles. The contribution to local adaptation could be 1) almost only from alleles arising around the time of first environmental change (recurrent expansions with a patchy map), 2) almost only from the time of the last environmental change (recurrent environmental changes with a patchy map or recurrent expansions with a cline map), or 3) from alleles arising around the times of both the first and the last environmental change (recurrent environmental changes with a cline map). In the third case, there were also subtle contributions around the time of the other nine environmental changes in the middle (Figure 5B).

The autocorrelation of environmental maps had different effects in different models. In general, replacing a cline map with a patchy map largely increased both adaptive and maladaptive effects from the most recent alleles, presumably due to migration from nearby patches with contrasting environments. For models with a single environmental change, replacing a cline map with a patchy map only slightly reduced the contribution of alleles from the time of environmental change, and slightly increased the total contribution from new mutations arising after the environmental change (Figure 5C and 5D). However, for models with recurrent population expansion and contractions, replacing a cline map with a patchy map almost removed the peak around the time of the last environmental change (Figure 5C and 5D). This may reflect that the implemented patchy environmental map has a higher capacity for maintaining standing genetic variation in “refuges” during population contractions, compared to a cline map.

### Young neutral alleles were more likely to correlate with environmental factors

For species with restricted dispersal, the spatial distribution of alleles is likely related to allele age. Specifically, we predicted that younger alleles would tend to be more associated with environmental factors by chance and thus have a higher false positive rate (FPR) in genotype-environmental association (GEA) analyses. To test this prediction, we compared the false positive rates of GEA among neutral alleles of different ages. For this analysis, we also included models with no selection (Figure 1 MNeuCon and MNeuExp) to distinguish the effects of isolation by distance and isolation by adaptation.

In models with constant population size and a cline map, we observed that the youngest neutral alleles had the highest FPR. As allele age increased, FPR quickly decreased and remained at a low value (Figures 6A and S7). This pattern held with or without selection, suggesting that the decrease of FPR with allele age was mainly mediated by limited dispersal, rather than selection. The presence of spatially heterogeneous selection increased the overall FPR in a cline map, presumably due to neutral alleles hitchhiking with locally adaptive ones.

**Figure 6.**
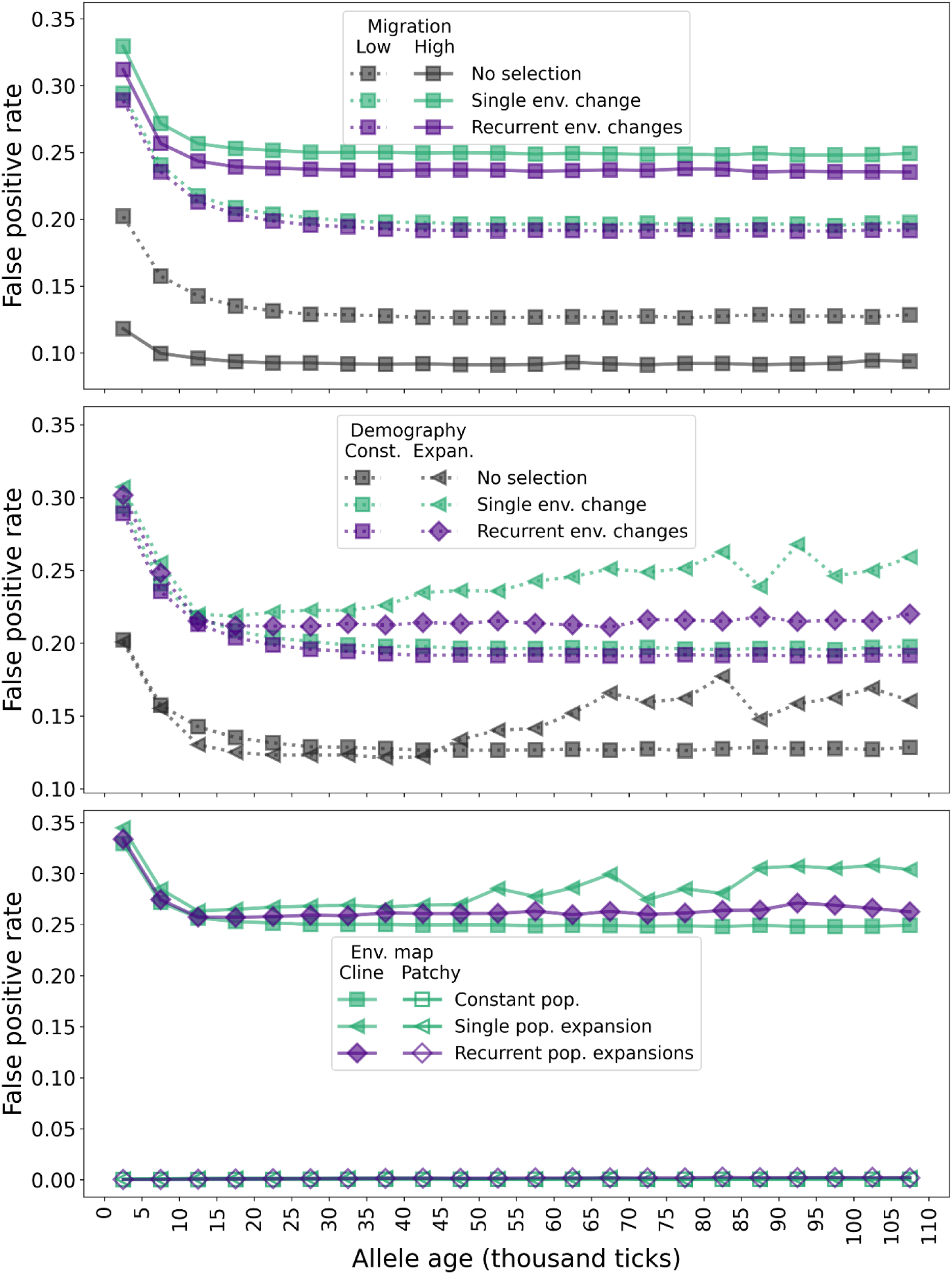
The relationship between allele age and false positive rate (FPR=false positives/(false positives + true negatives)) of neutral alleles in genotype-environment association analysis among simulations with a high polygenicity (the standard deviation of the distribution of phenotypic effect size *σ*_*M*_ = 0.01, and functional mutation rate *μ*_*f*_ = 1 × 10^−8^ per base pair per simulated year). (A) The effect of migration rate in simulations with a constant population size and a cline environmental map; The high and low migration levels correspond to average overall *F*_*ST*_ values of 0.112±0.008 and 0.038±0.002, respectively. (B) The effect of demographic history in simulations with a low level of migration and a cline environmental map; Const.: a constant population size; Expan.: a single population expansion or recurrent population expansions/contractions. (C) The effect of environmental maps in simulations with a high level of migration. In all plots, colors indicate the mode of selection, with grey for models with no selection, teal green for a single environmental change, and purple for recurrent environmental changes. The FPR of neutral alleles in each age category was calculated as the number of neutral alleles that had a p-value<1 × 10^−10^ in Kendall’s rank correlation test divided by the total number of neutral alleles within each age category, using results of 200 simulation replicates. Alleles with minor allele frequency < 0.05 were excluded from the analysis. Allele ages are shown in thousand SLiM ticks. One tick is equivalent to a simulated year, in which all individuals reproduced once.

Migration rates had different effects on FPR in models with and without selection (Figures 6A and S7). When there was no selection, the higher migration rate largely reduced the inflation of FPR in young alleles. As allele age increased, FPR more quickly decreased to a steady low level, and the overall FPR was lower than that in models with the lower migration rate. This effect of migration rate also suggests that the pattern is related to the spatial distribution of alleles. In the presence of selection, the higher migration rate still led to a faster decrease of FPR as allele age increased, but also a higher overall FPR.

When there was a single population expansion, FPR usually increased again in older alleles, though there was wide variation around these FPRs (Figures 6B and S7). The larger variation in older alleles may reflect a smaller sample size in these age categories and the stochastic effect of allele surfing. However, when there were recurrent population expansions and contractions, the FPR did not increase again in older alleles, but was only mildly higher than that in a constant population (Figures 6B and S7).

Compared to models with a cline map, a patchy environmental map greatly decreased the overall FPR (Figures 6C and S7), showing that the confounding effect of population structure is less severe in a patchy environment map than in a cline map. A patchy map also distorted the relationship between FPR and allele age, resulting in no obvious relationship between FPR and allele age (Figures 6C and S7).

## Discussion

### Predicting the genetic architectures of local adaptation requires information on mutation and demographic history

Studying the genetic basis of local adaptation relies on assumptions about adaptive alleles, as different methods target alleles with different numbers, effect sizes, frequencies, and ages (Berg and Coop 2014; Field et al. 2016; Lasky et al. 2023). With growing evidence from experimental evolution (Barghi et al. 2019; Besnard et al. 2020) and large-scale association studies (Shi et al. 2016; De La Torre et al. 2021; Yengo et al. 2022), an emerging consensus is that many traits of interest are highly polygenic (Walsh and Lynch 2018), in the sense that many new mutations can affect a trait, and many functional alleles are maintained in a population as standing genetic variation. What is less clear is to what extent heterogeneous selection with migration reshapes the attributes of a set of alleles that directly constitute local adaptation. Small-effect mutations may be abundant in number, but not necessarily have a large total contribution to local adaptation, not only due to their trivial individual effects, but also because they are less resistant to the homogenizing effect of migration during local adaptation (Griswold 2006; Yeaman and Otto 2011; Yeaman and Whitlock 2011).

We modeled local adaptation in some typical demographic and environmental scenarios. The resulting genetic architecture of local adaptation was highly variable and strongly depended on the rate and effect size of raw mutations and demographic history (Figures 2, 3, and 5). The polygenicity of the trait under selection led to contrasting genetic architectures of local adaptation. With a similar level of local adaptation (Table S2), on average, 3 alleles could explain 80% of local adaptation in our models of low polygenicity, whereas about 190 alleles were needed in models of high polygenicity (Figure 3). Compared to the overwhelming effect of polygenicity, the effects of migration levels and the autocorrelation of environmental maps seemed limited (Figure 3), despite the values of these two parameters also representing a typical range or pattern for tree species (Methods).

Models of low polygenicity reproduced some classic predictions from population genetics, such as locally adaptive alleles being maintained in a population for a long time by spatially heterogeneous selection (Figure S3), and small-effect or low-frequency alleles having almost no contribution to local adaptation (Figures S1 and S2). However, in the models of high polygenicity, low-frequency alleles (frequency<0.1) made a considerable contribution to local adaptation (∼30%, Figure S2), although many of these alleles may not be identified by association studies with a limited sample size (Croucha and Bodmer 2020; Lasky et al. 2023) or tests of selection (Voight et al. 2006; Hejase et al. 2022) and genetic differentiation (Jakobsson et al. 2013). We also observed the turnover of locally adapted alleles, as in previous studies (Yeaman 2015; Yeaman 2022). In models of high polygenicity, locally adaptive alleles were continuously replaced by younger adaptive alleles, but the level of local adaptation almost remained the same (Figures 2E, S3 and S4). It is worth noting that compared to the time scale of environmental changes, this turnover can be a slow process. For example, 20,000 simulated years after a major environmental change (similar to the time since the last glacial maximum, Hughes, 2022), for a population of 5000 individuals with overlapping generations and annual reproduction, more than half of all local adaptation could be explained by alleles that arose when local adaptation was first established (Figure S3).

Demography is also a strong factor affecting the genetic architecture of local adaptation. Previous studies have shown that population growth increases the relative contribution from low-frequency deleterious alleles to the genetic variance of complex traits, making deleterious causal alleles harder to detect in GWAS (Lohmueller 2014), and demographic changes can explain up to 55% of variation of allele frequency-effect size matrices in a simulation of global adaptation after a sudden change of environmental optima (Stetter et al. 2018). Here, we showed that population expansion also affected the number, effect size, frequency, and age of alleles that constitute local adaptation (Figures 3, 5, S1 and S2). A single population expansion could reduce the minimum number of alleles needed to explain 80% of local adaptation (*K*_80,*pos*_) by about 40% (Figure 3, High polygenicity), and increase the relative contribution from alleles with large effects, high frequencies or ages around the time of environmental change (Figures S1-S3). Similar to the observations of Lohmueller (2014), we also found that low-frequency deleterious alleles (alleles with negative *LF*_*mut*_) had larger relative contribution to the total maladaptive effects after a population expansion (Figure S2). This effect was most obvious when there was only a single round of local adaptation with population expansion.

The effect of population expansion on the number of adaptive alleles is long-lasting. It took at least 60,000 simulated years for *K*_80,*pos*_ to recover from the effect of a single population expansion and reach the level observed in the corresponding model without a population expansion (Figures 4A and S5). Although the number of locally adapted alleles kept increasing over time after population expansion, the total net contribution to local adaptation almost stayed constant. The absolute contribution from large-effect or high-frequency alleles mildly decreased over time after a population expansion, but this effect alone is not enough to explain the increase in *K*_80,*pos*_ (Figures S1 and S2, Table S3). It was mostly the absolute contribution from low-frequency and small-effect young alleles that kept increasing and compensating for the growing maladaptive effects of new deleterious alleles (Figures 2E, 4, and S1-S3). From a practical perspective, local adaptation soon after a population growth is expected to be achieved by a smaller number of alleles with larger effect sizes and higher allele frequencies. However, while recent population growth should increase the chance of identifying a few underlying alleles making the largest contribution to local adaptation, such alleles may be difficult to identify with GEA, as population expansion is known to be a major source of false positives (Lotterhos and Whitlock 2015).

The variability of the genetic architecture of local adaptation stresses the importance of investigating the distribution of mutational effect size for diverse organisms and traits. Information on the polygenicity of traits and population demographic history can help to set a meaningful goal and choose methods accordingly when studying the genetic basis of local adaptation. Although the contrasting parameter sets in our models represent a plausible and wide parameter space, those values should not be regarded as lower and upper limits. Considering that many complex traits can be affected by thousands of alleles that are scattered across a genome (Shi et al. 2016; Yengo et al. 2022), it is likely that we have not captured the upper boundary of polygenicity, and our simulations only displayed a partial spectrum of the genetic architecture of local adaptation.

### Allele age interacts with multiple variables in GEA

The confounding effect mediated by population structure has long been a serious problem (Meirmans 2012; Lotterhos 2023). When controlling population structure in GEA, it is often assumed that population structure affects all alleles in a similar way (Lasky et al. 2012; Günther and Coop 2013; Gautier 2015). However, the effect of population structure may vary across alleles of different ages, so adding population structure alone to a statistical model may not be able to fully account for its confounding effects.

To understand how population structure interacts with allele age, we modelled population range expansion for a species with spatially-restricted mating and dispersal in a continuous space. Both processes are common and known for generating population structures that confound GEA (Eckert et al. 2010; Meirmans 2012). Our results suggest that allele age is indeed a moderator variable for the confounding effects from population structure (Figure S8, path 6). Young neutral alleles were more likely to show a significant association between allele frequency and environmental factor than older alleles, resulting in an L-shape curve of FPR across allele age (Figure 6). This decrease of FPR in older alleles occurred in all models with a cline environmental map, both with and without selection, but can be reduced by a higher migration level in the population without selection, suggesting that this effect is mediated by a population structure generated by isolation by distance. When there was a range expansion along the axis of the environmental gradient, FPR gradually increased again in older alleles after the initial drop, leading to a “U” shape curve (Figure 6B). This increase of FPR in older alleles may be explained by the effect of “allele surfing”. Allele surfing is the phenomenon that population range expansion stochastically increases the frequency of low-frequency alleles and forms allele frequency clines, especially along the expansion axis (Klopfstein et al. 2006). The chance of “surfing” is expected to be proportional to the initial allele frequency before range expansion (Excoffier and Ray 2008). As older alleles are more likely to be present at higher frequencies than younger alleles (Figure S9), they are more likely to form a frequency gradient after a range expansion, and therefore have a higher FPR in GEA.

Apart from moderating confounding effects from population structure, allele ages may also strongly modify the causal effect of environmental variables on genotypes (Figure S8, path 7). In other words, the probability that an allele has participated in local adaptation may not be constant but greatly varies across allele age. Treating the allele age of locally adaptive alleles as a variable revealed a great variation in the contribution to local adaptation along allele age (Figure 5). In particular, our results showed that alleles arising around the time of environmental changes may have disproportionately large contributions to local adaptation, especially in cases with a lower polygenicity, a population expansion, or a new type of environmental change (Figure 5). The level of polygenicity again had the strongest effects. When mutations were rare and had larger effect sizes, new mutations arising around the time of environmental changes were almost the only source of local adaptation (Figure 5).

For GEA analyses, it may be worth adding both allele age and its interaction with population structure and environmental variables to statistical models. When a species/population is known to have limited dispersal abilities, a recent population growth, or affected by clinal environmental variables, adding the interaction between allele age and population structure should more appropriately control the confounding effects from population structure and reduce FPR in GEA.

Adding the interaction between allele age and environmental variables to control the modifying effect of allele age should increase sensitivity (TPR) and also reduce FPR. However, the exact relationship between allele age and the extent of genotype-environment causal association may not be linear but a complex function, and highly variable across genetic and ecological scenarios (Figure 5). This variability may make this second form of interaction practically hard to model.

## Data availability

Scripts are available at GitHub (https://github.com/euptelia/arg-for-gea).

## Acknowledgements

Thanks to Peter Ralph and Martin Lascoux for discussions. We would like to thank Kung-Ping Lin and Rafael Candido-Ribeiro for their constructive comments on the presentations of this project, which have led to modifications of the analyses. TRB and MCW both gratefully acknowledge support from NSERC Discovery Grants.

## Notes

### Competing Interest Statement

The authors have declared no competing interest.

