## Supplementary Figure S1 for "The genetic architecture of local adaptation is historically contingent"

**Supplementary figure S1.** The contribution to local adaptation from alleles with different phenotypic effect sizes. Results were aggregated from 30 SLiM simulation replicates. High polygenicity: The standard deviation of the distribution of phenotypic effect size $\sigma_M = 0.01$, and functional mutation rate $\mu_f = 1 \times 10^{-8}$per base pair per simulated year; Low polygenicity : $\sigma_M = 0.1$, $\mu_f = 1 \times 10^{-10}$; Low-Mig./High-Mig.: a lower/higher migration level (corresponding to average overall $F_{ST}$ values of 0.112±0.008 and 0.038±0.002, respectively).

### High polygenicity

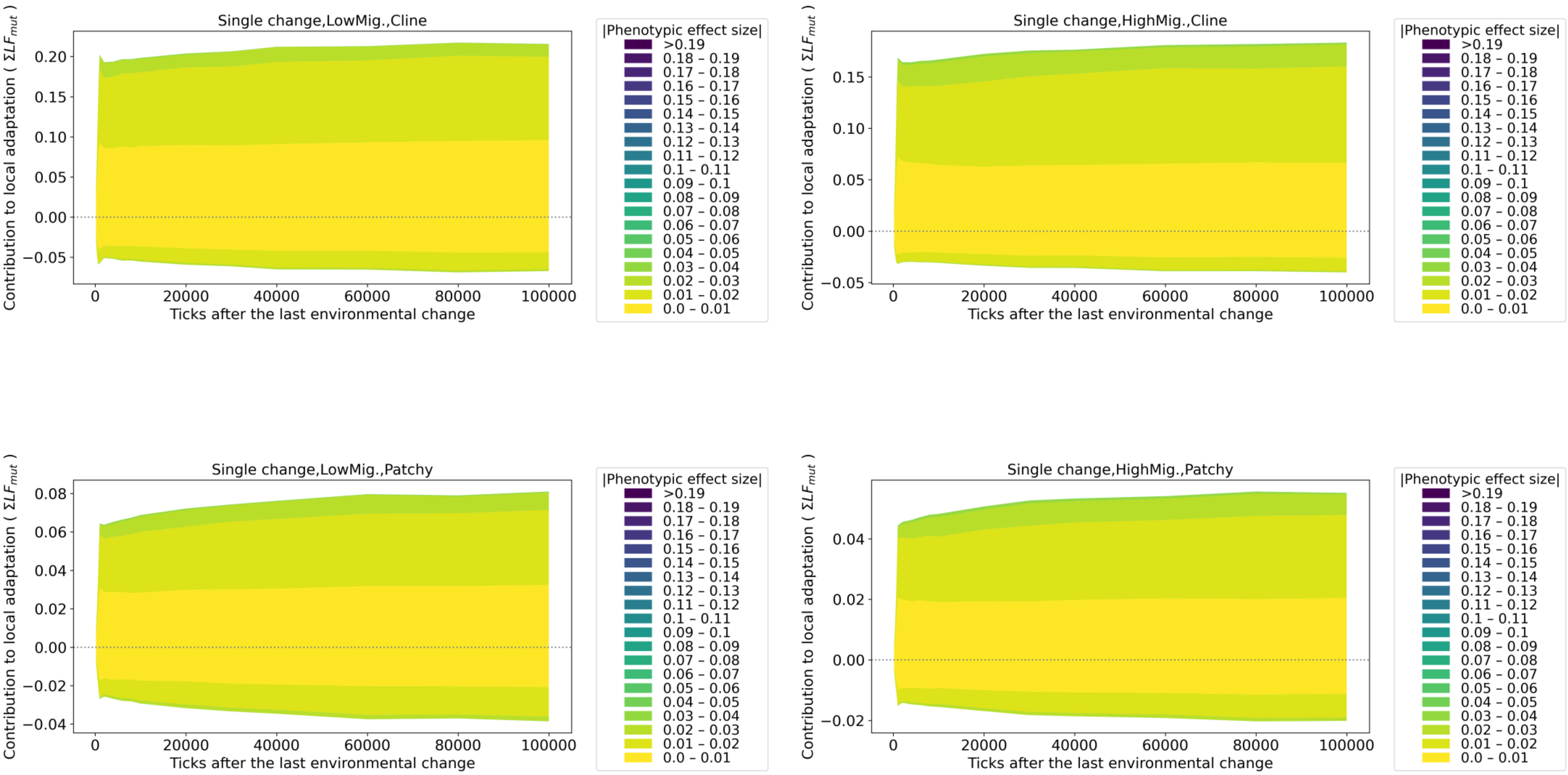

### High polygenicity

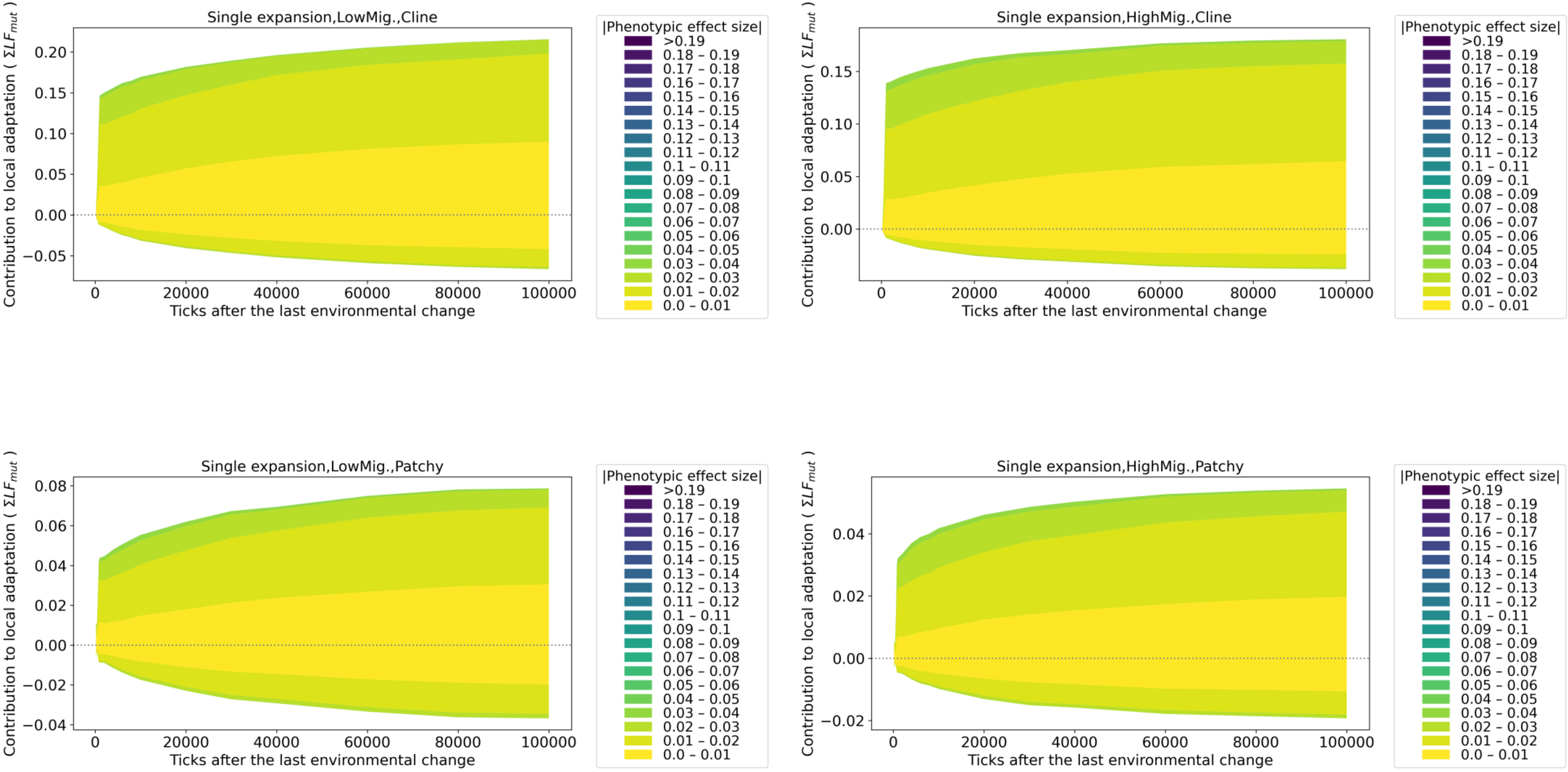

### High polygenicity

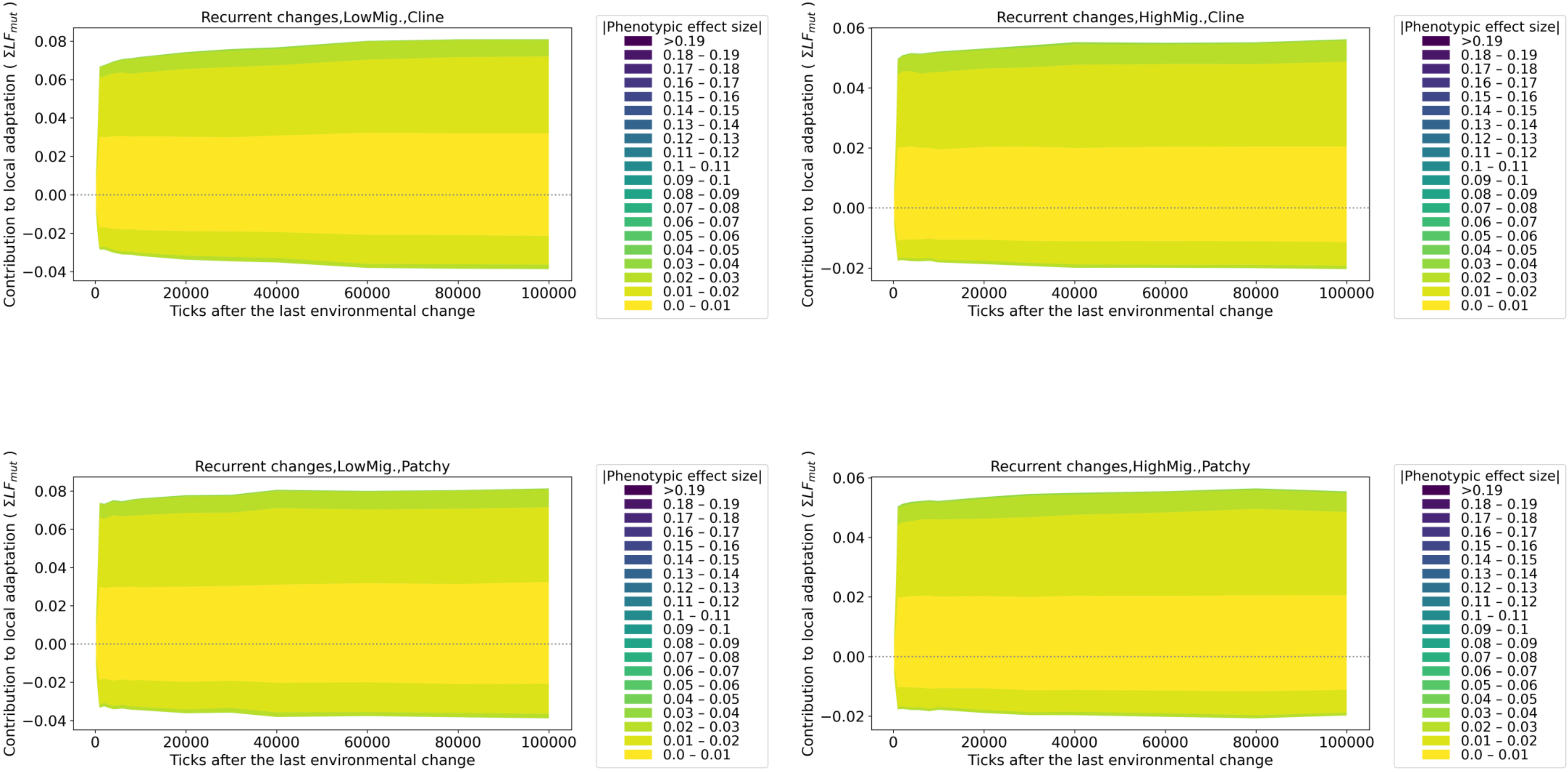

### High polygenicity

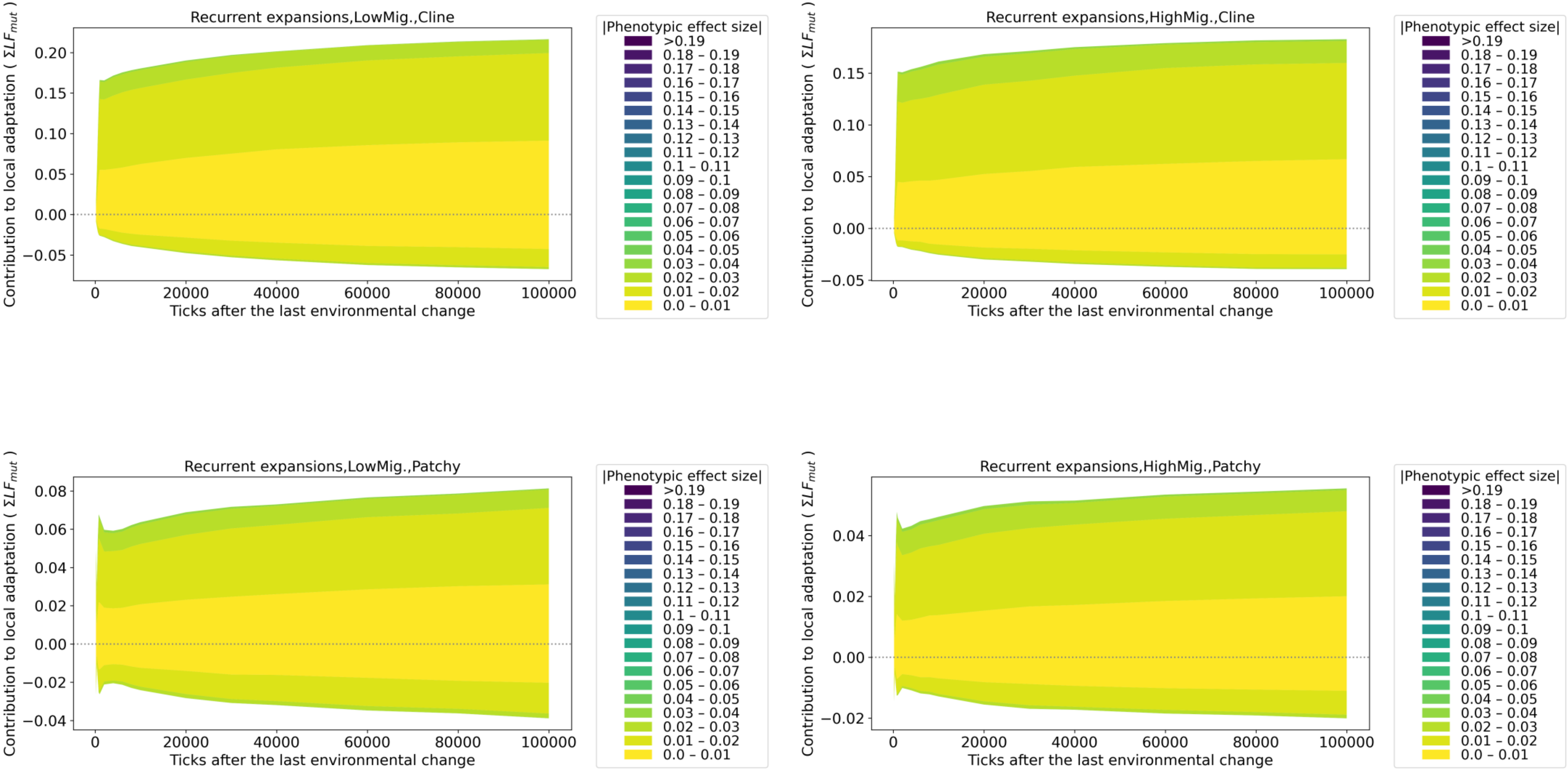

### Low polygenicity

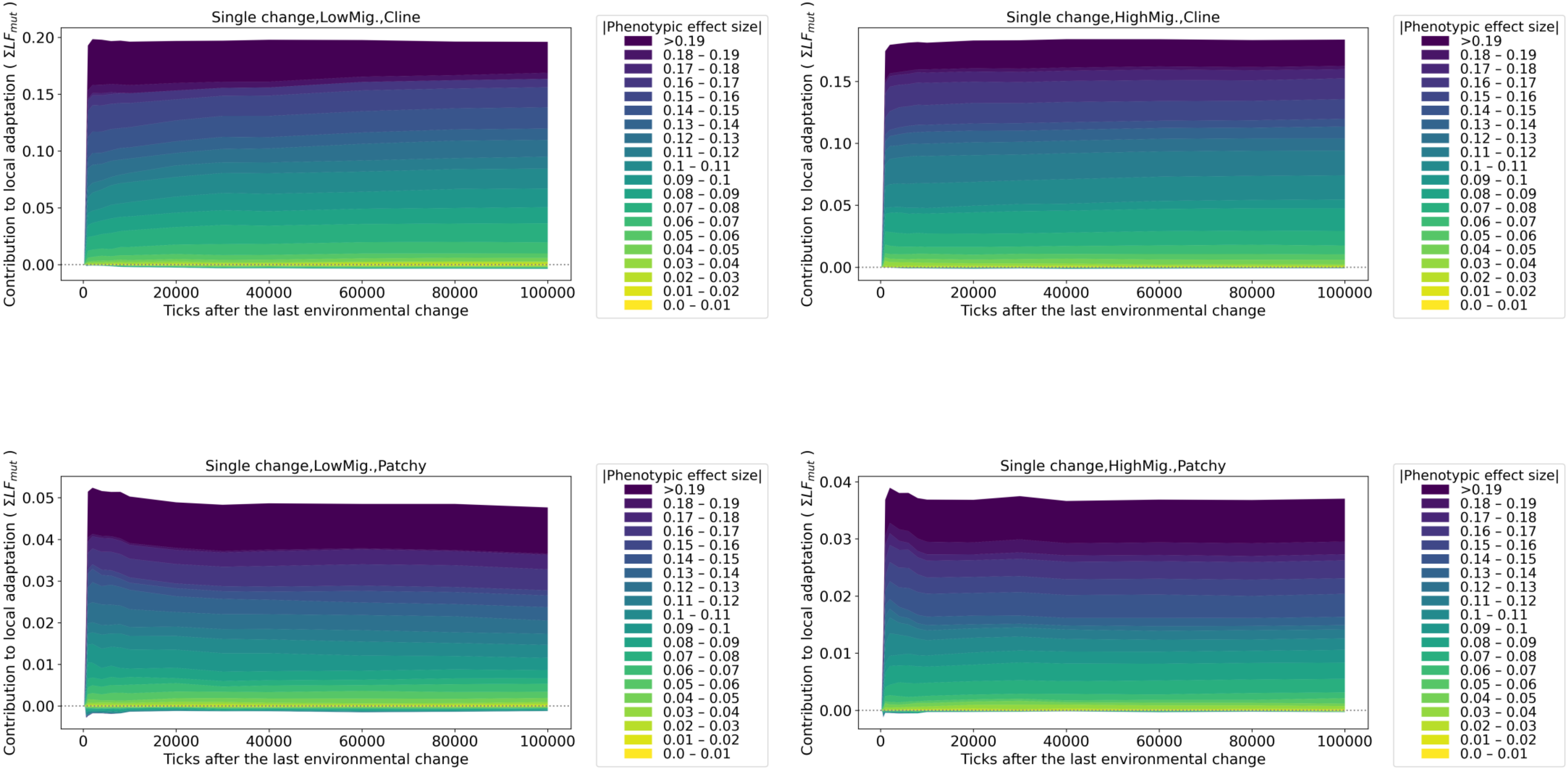

### Low polygenicity

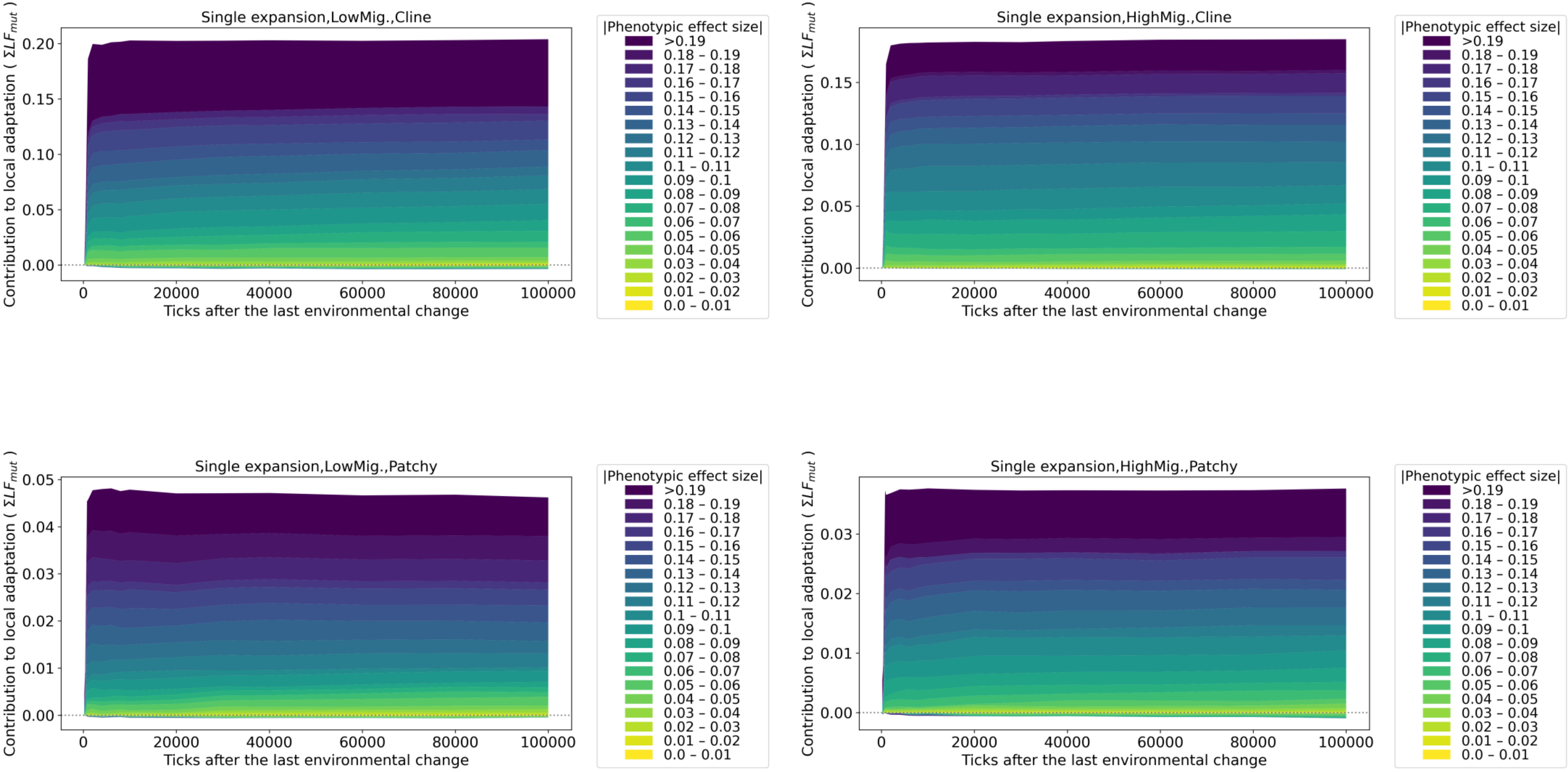

### Low polygenicity

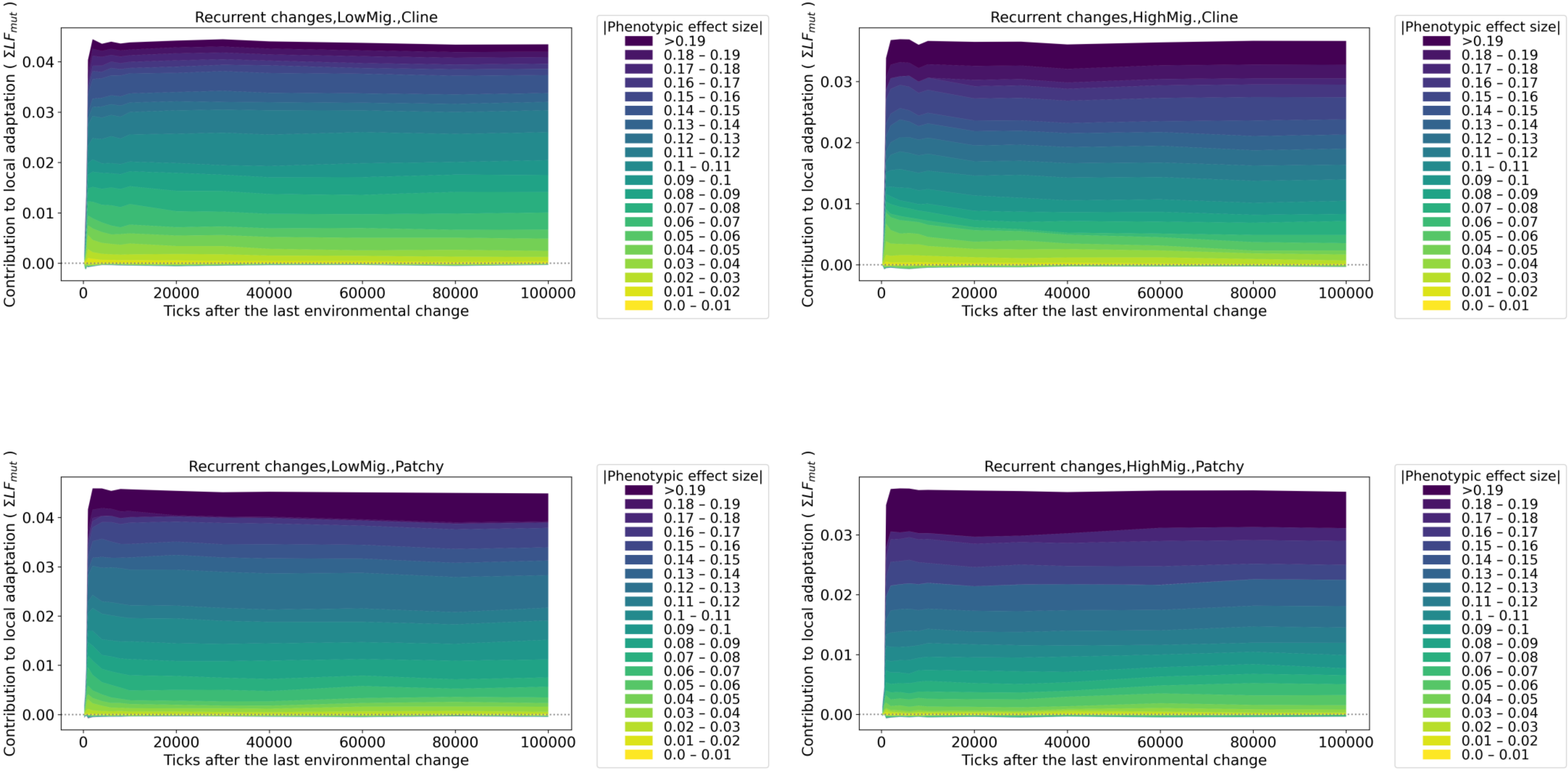

### Low polygenicity

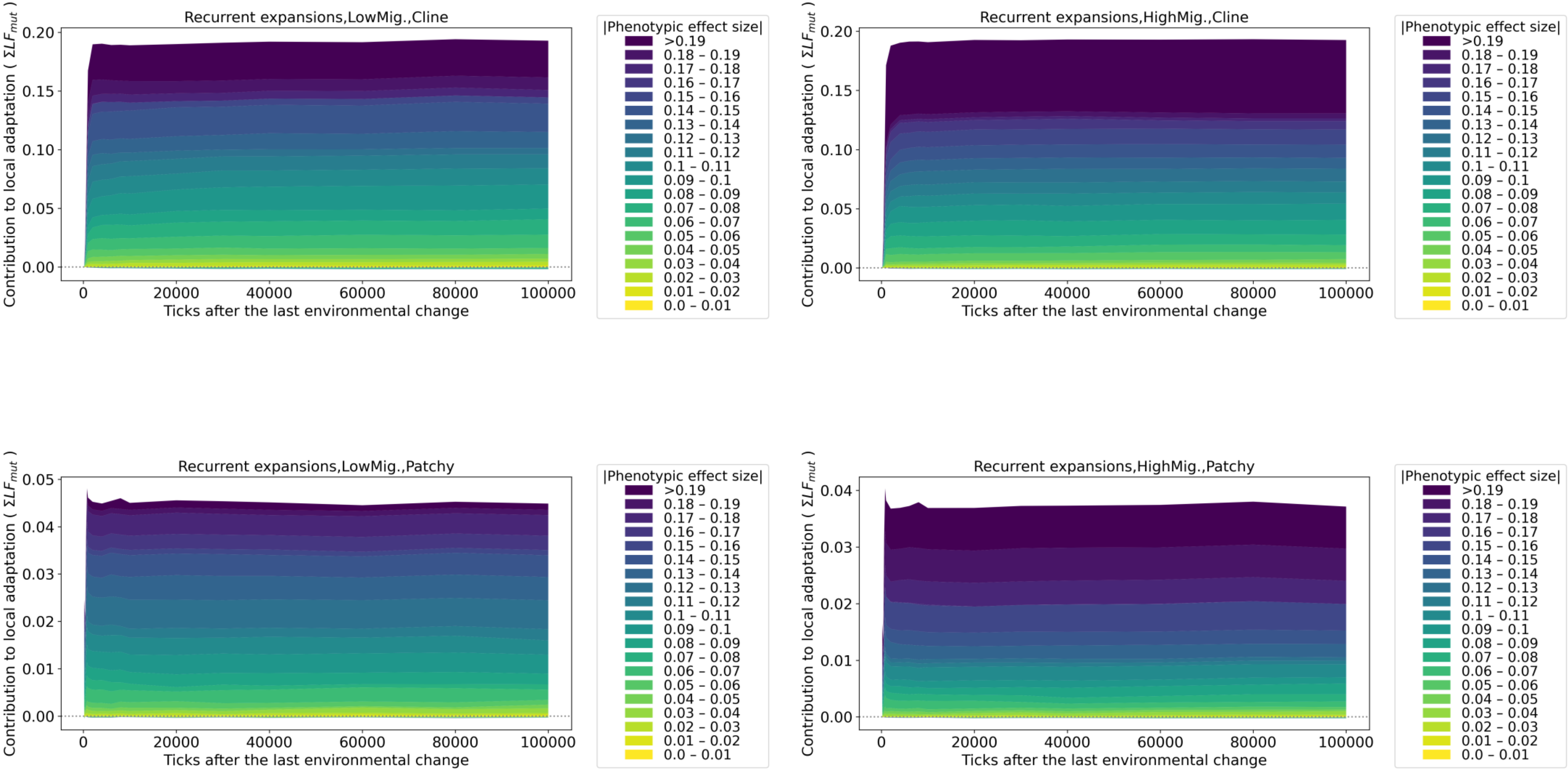
