## Supplementary Figure S6 for "The genetic architecture of local adaptation is historically contingent"

**Supplementary figure S6.** The relative contribution from alleles arising at different times to local adaptation was affected by multiple genetic and ecological parameters. Samples were taken at 10,000 simulated years after the most recent environmental change, with 200 SLiM simulation replicates. The red dashed vertical lines indicate the time of the (last) environmental change. For models with recurrent environmental changes or recurrent population expansions, grey dashed vertical lines indicate the starting times of the past ten rounds of environmental change. Contributions from alleles of different phenotypic effect size categories are distinguished by color. High polygenicity: The standard deviation of the distribution of phenotypic effect size $\sigma_M = 0.01$, and functional mutation rate $\mu_f = 1 \times 10^{-8}$per base pair per simulated year; Low polygenicity : $\sigma_M = 0.1$, $\mu_f = 1 \times 10^{-10}$; Low-Mig./High-Mig.: a lower/higher migration level (corresponding to average overall $F_{ST}$ values of 0.112±0.008 and 0.038±0.002, respectively).

### High polygenicity

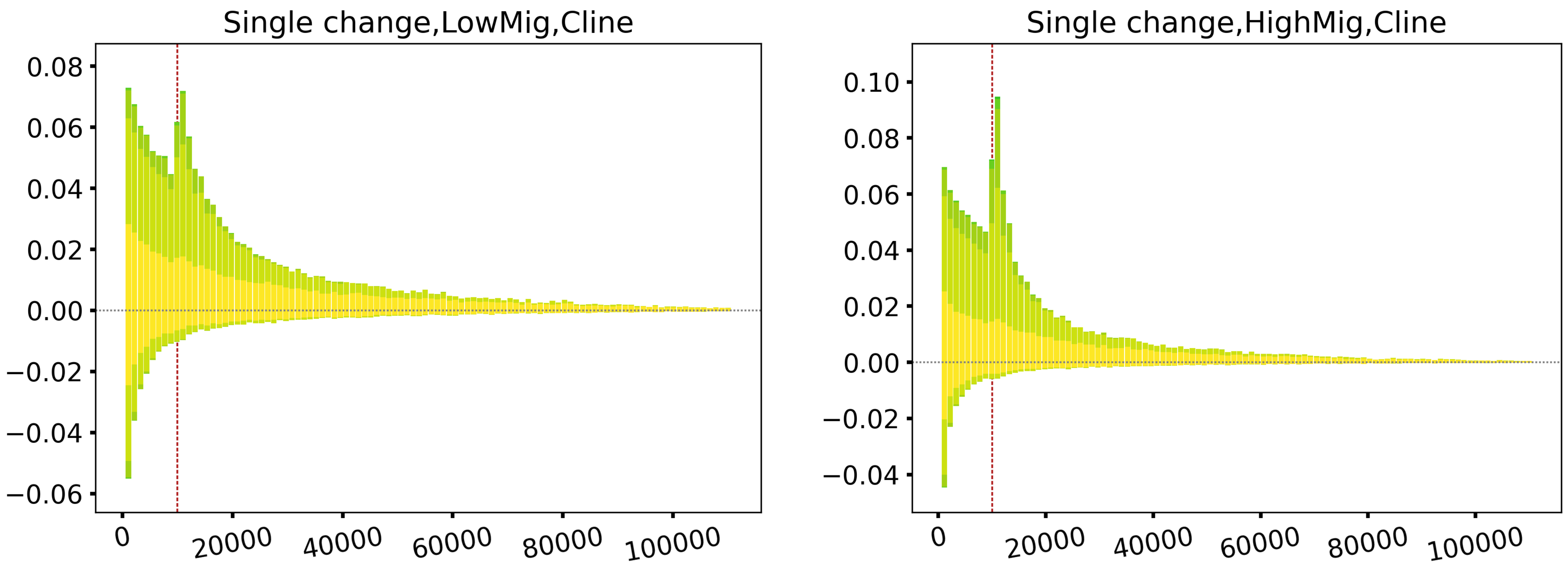


### Low polygenicity

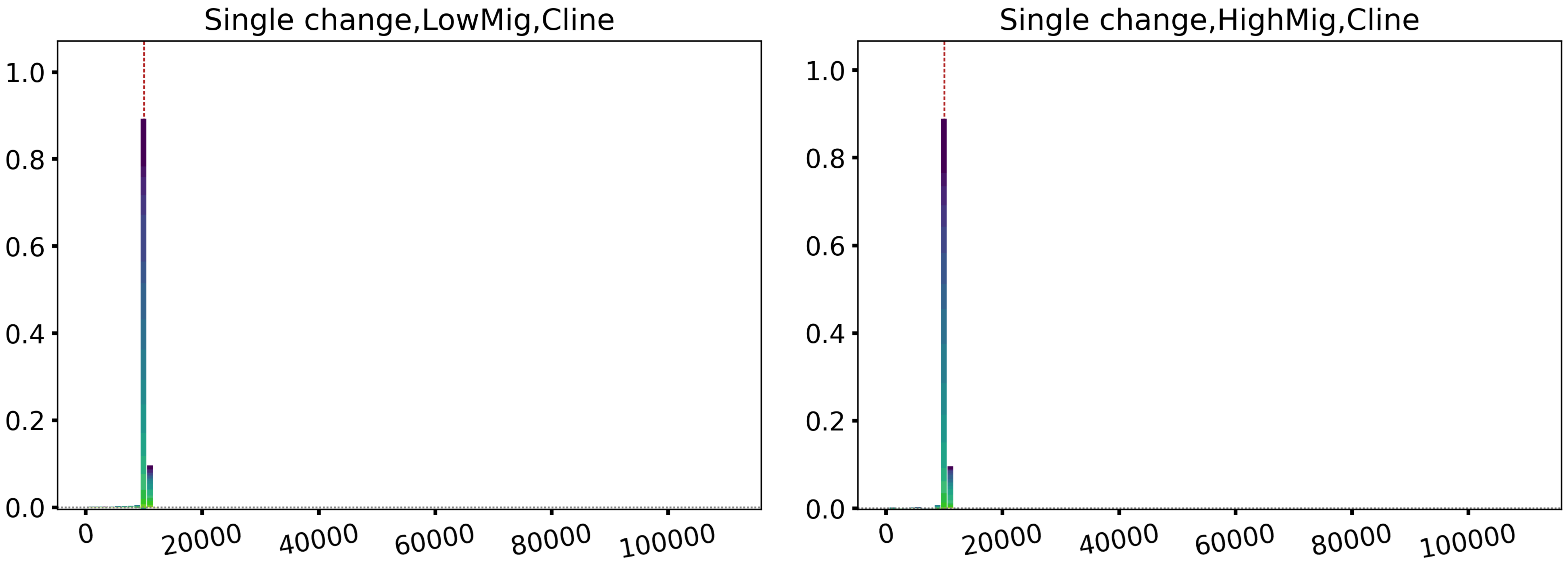

#### High polygenicity

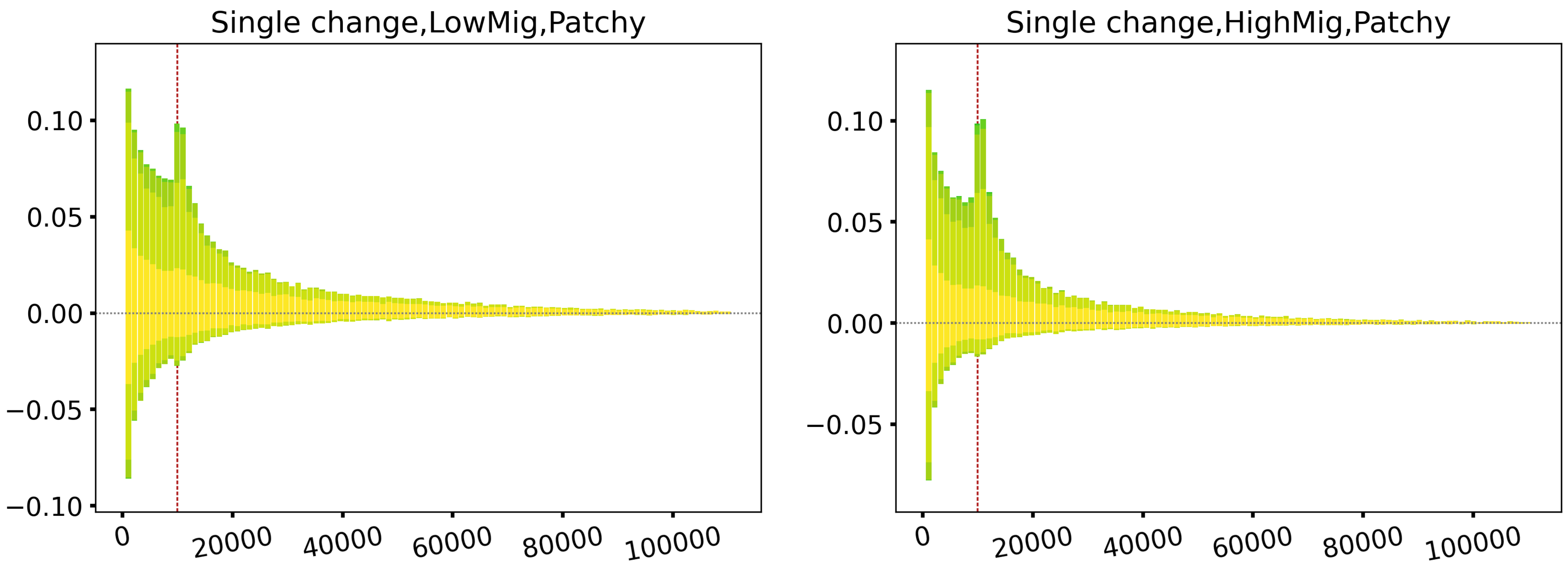


#### Low polygenicity

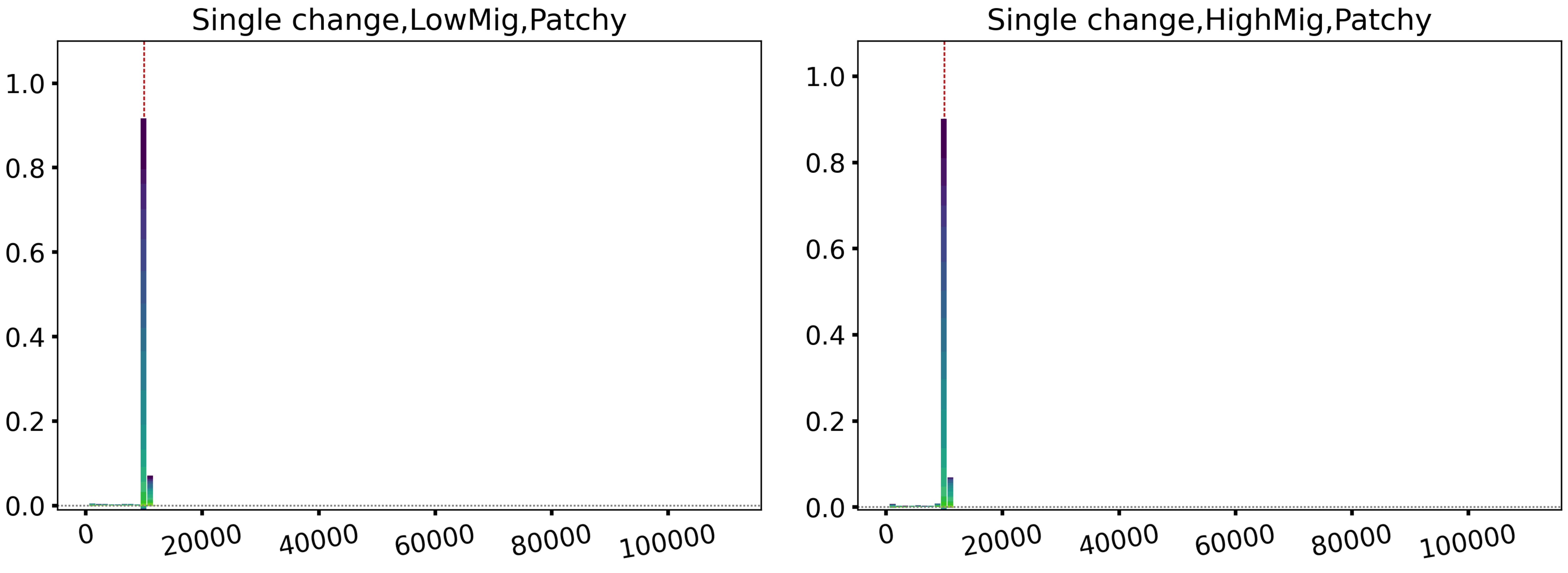

### High polygenicity

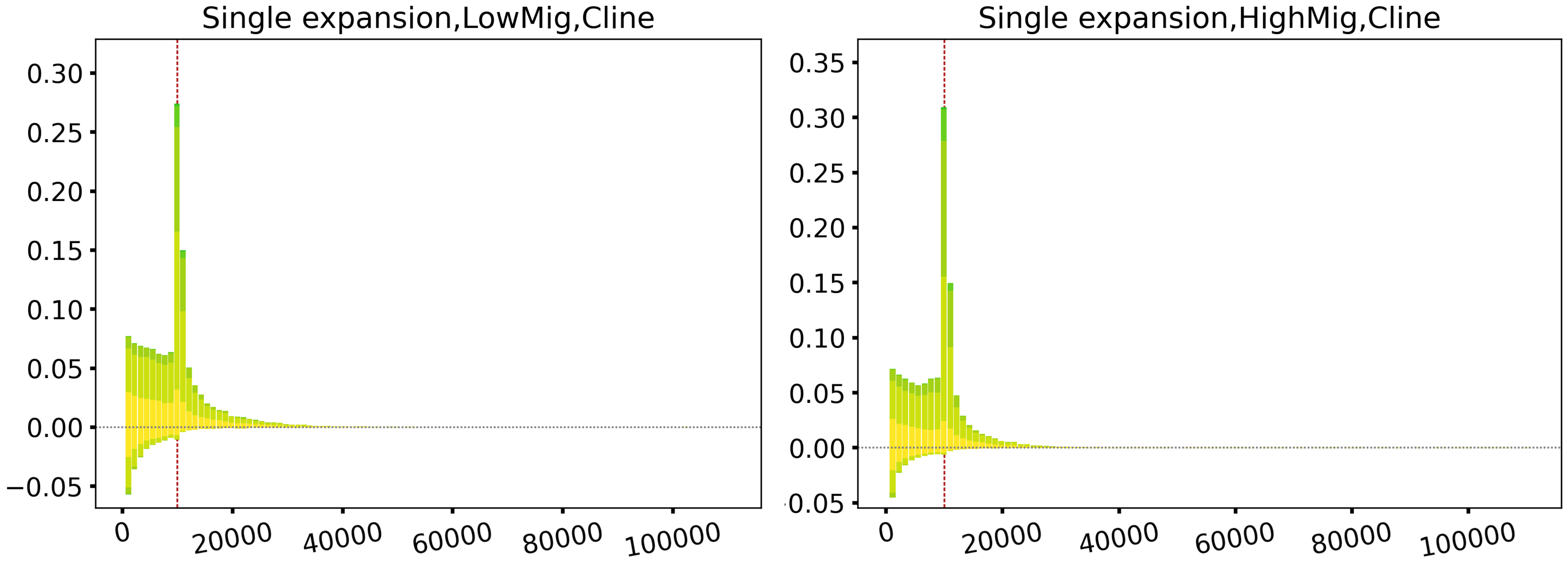


### Low polygenicity

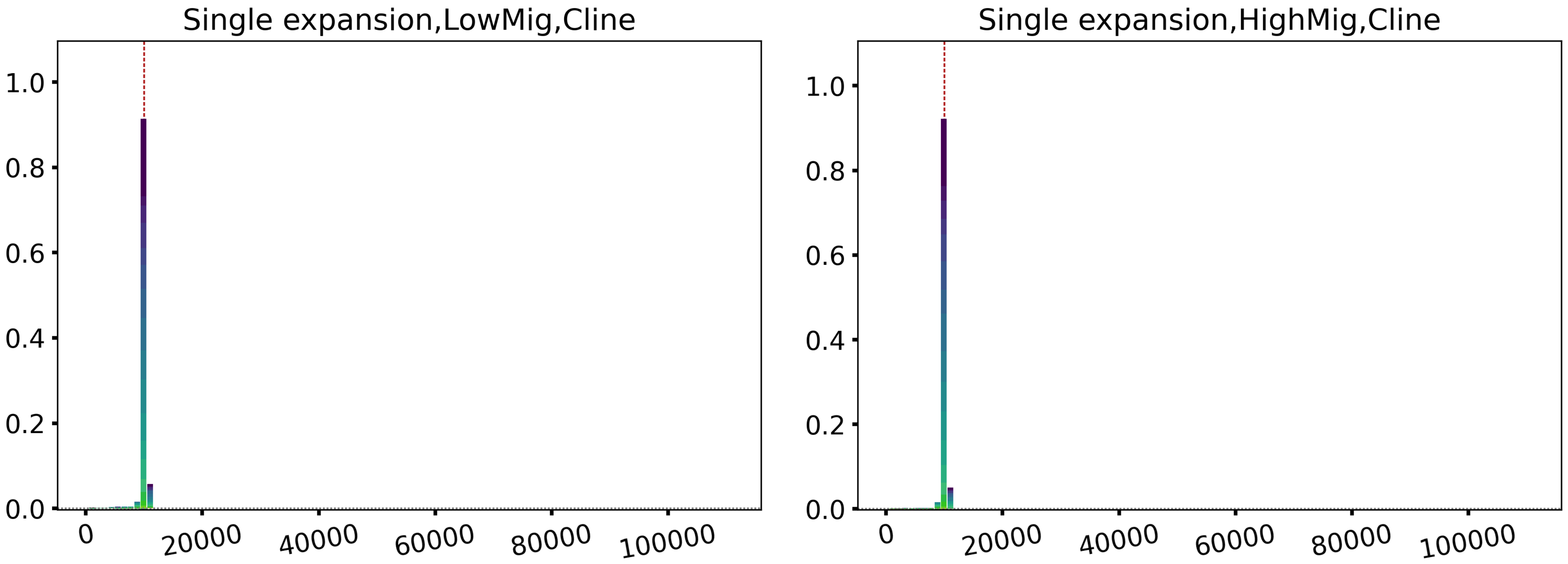

### High polygenicity

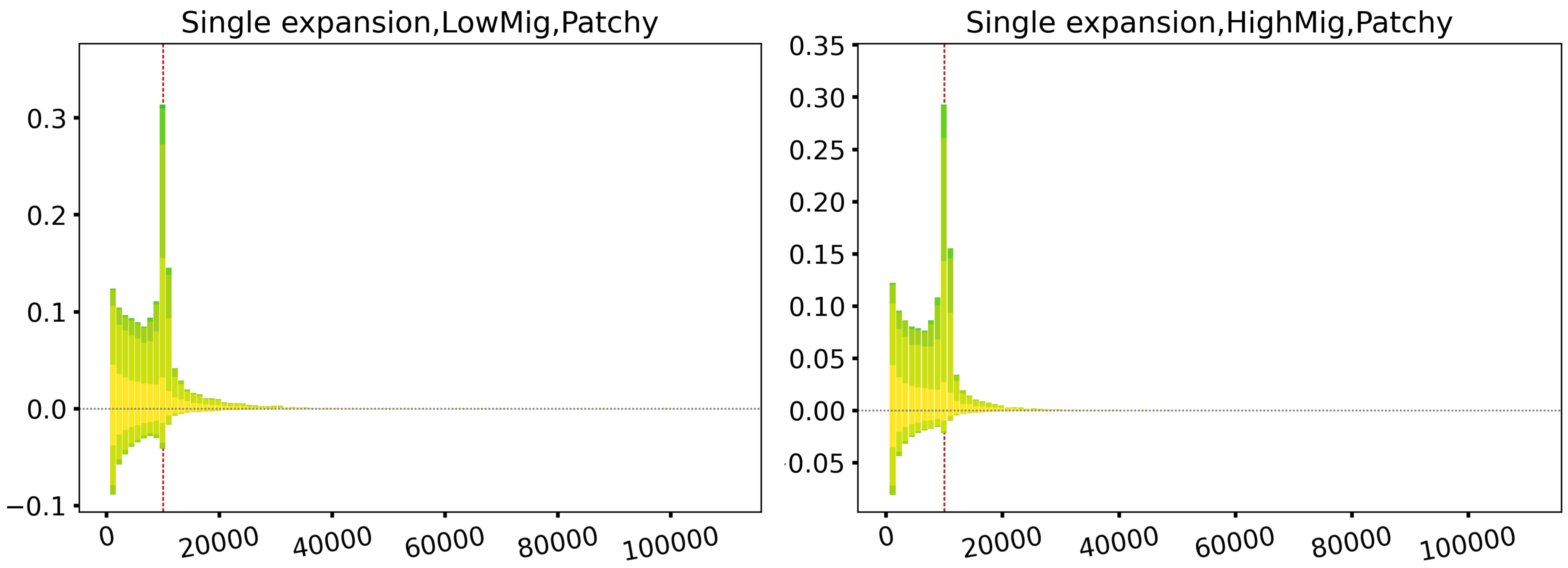


### Low polygenicity

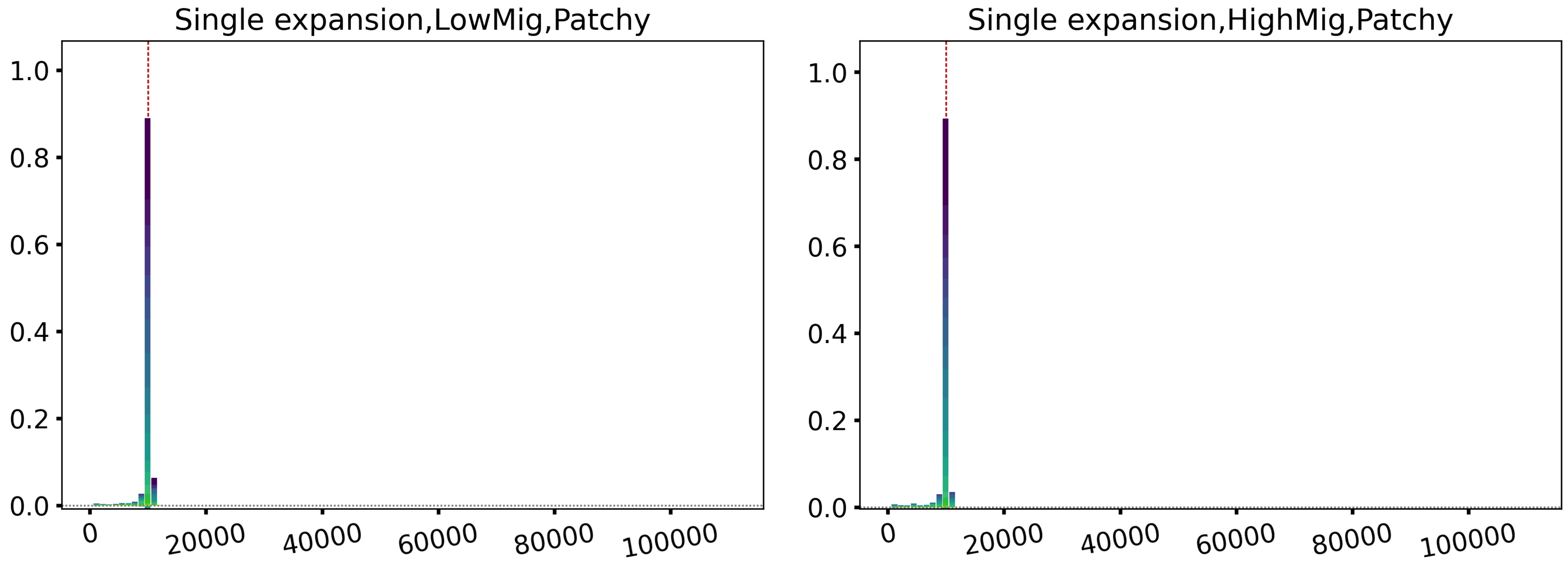

### High polygenicity

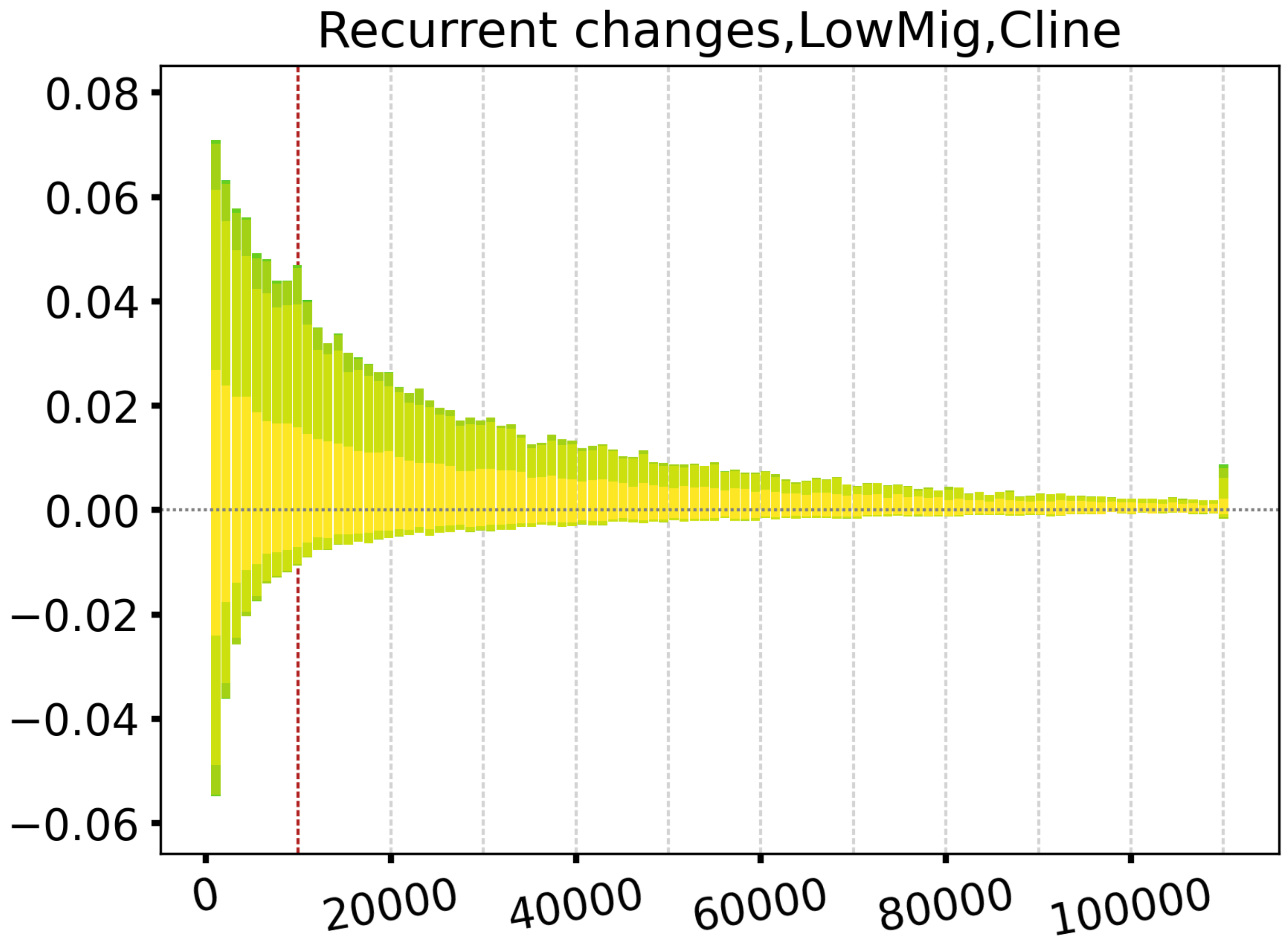


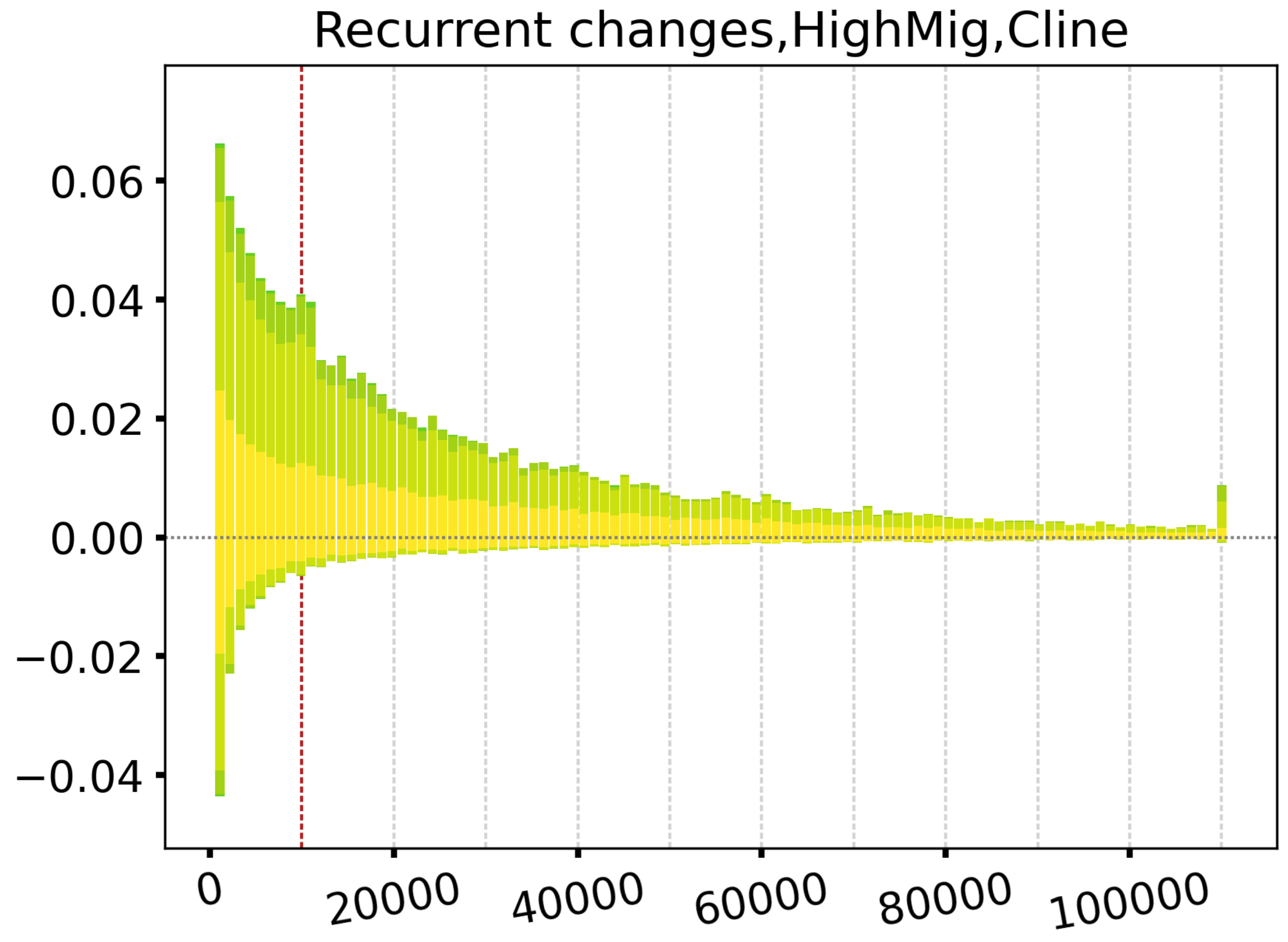


### Low polygenicity

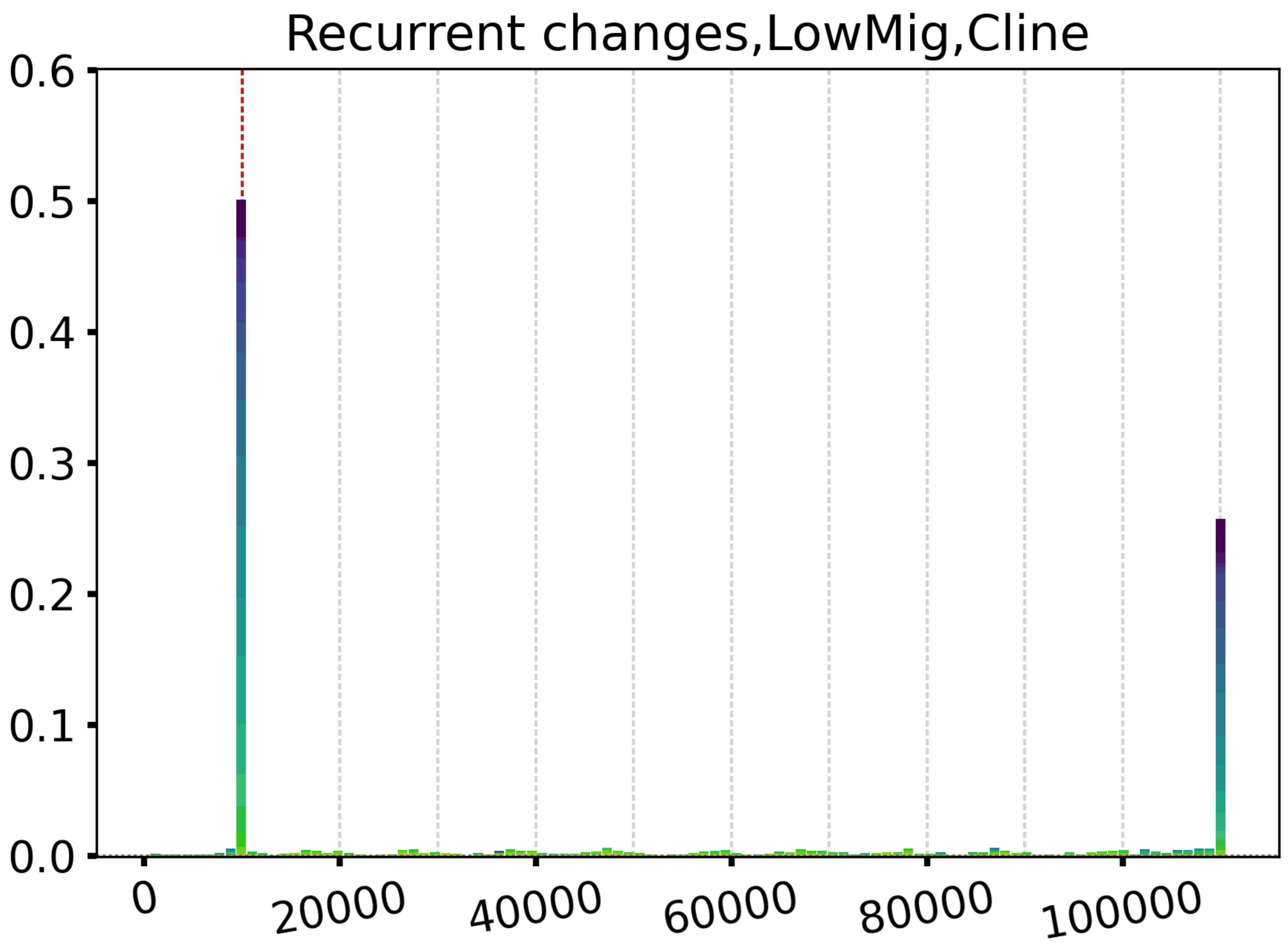


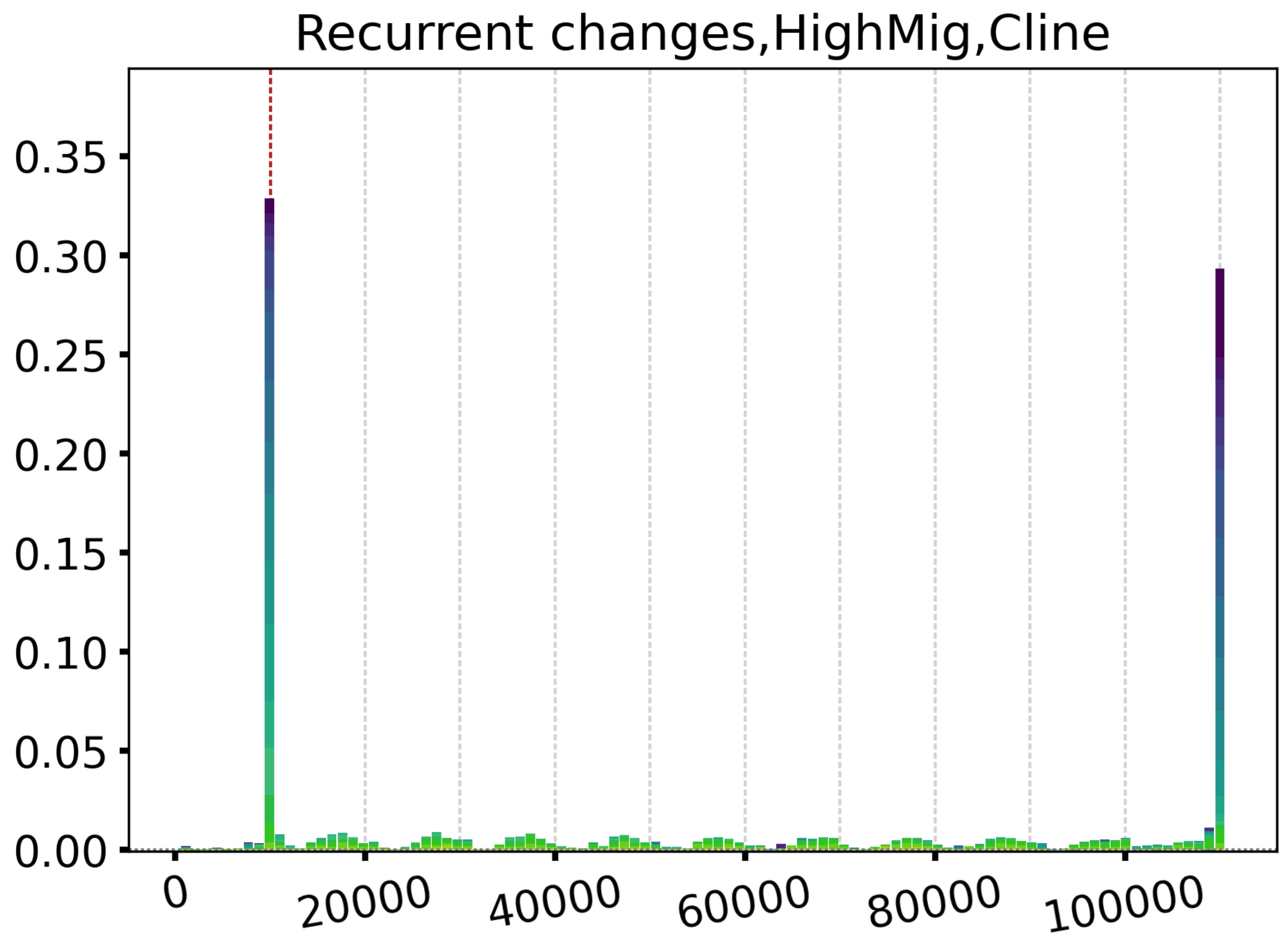

#### High polygenicity

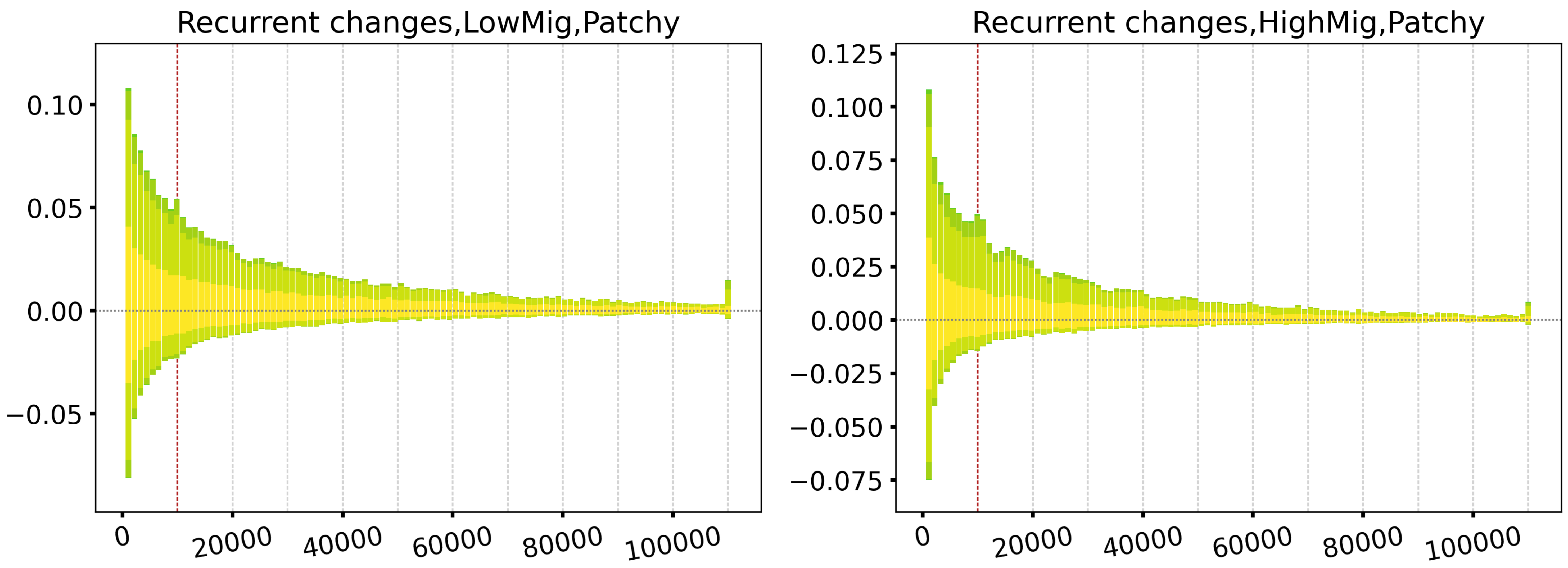


#### Low polygenicity

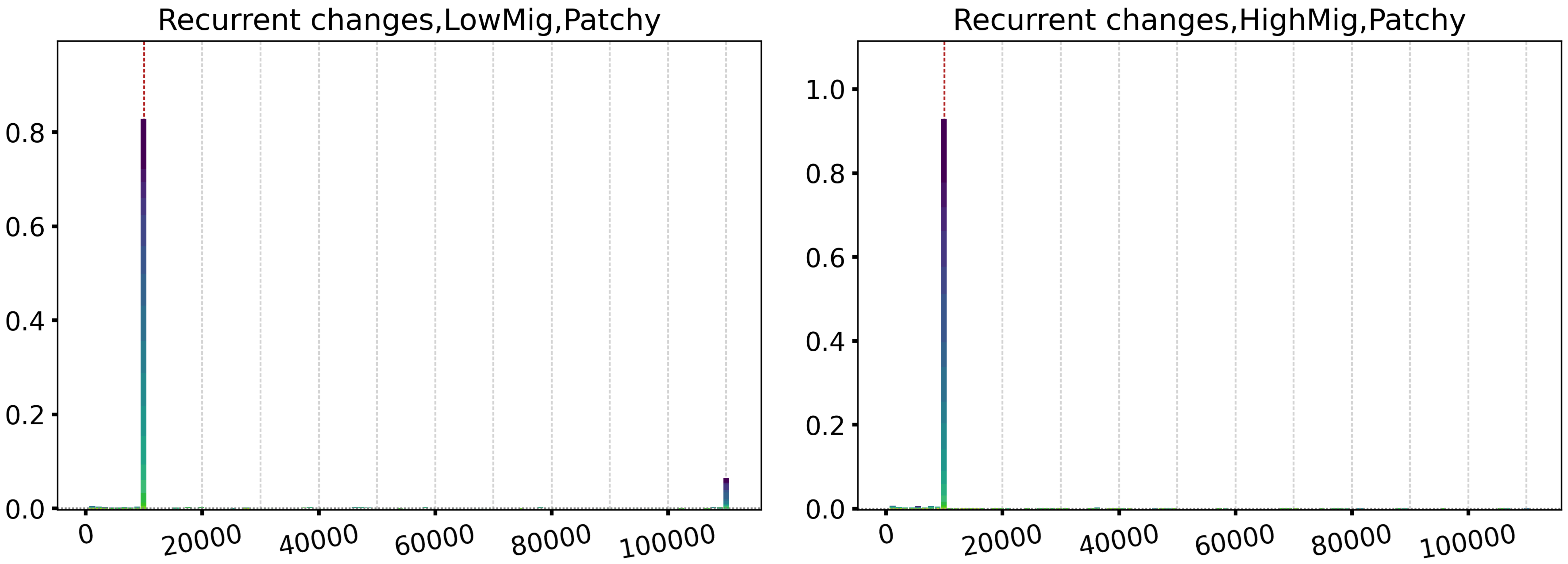

#### High polygenicity

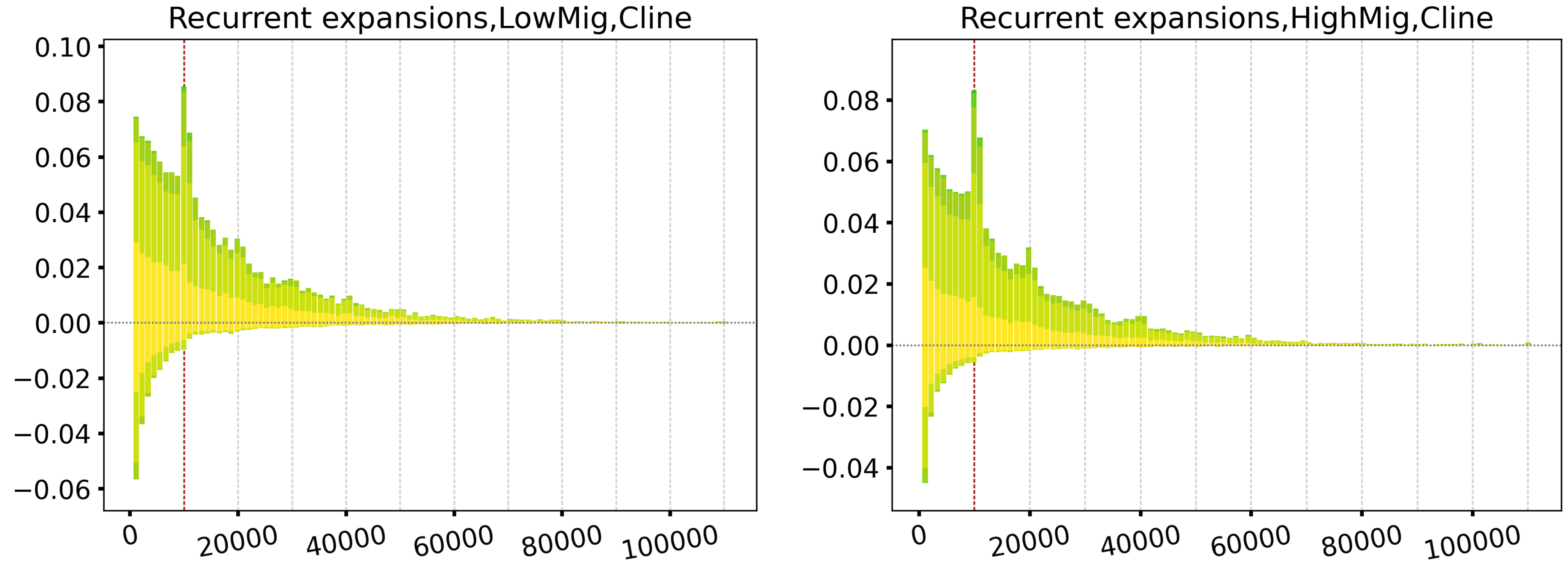


#### Low polygenicity

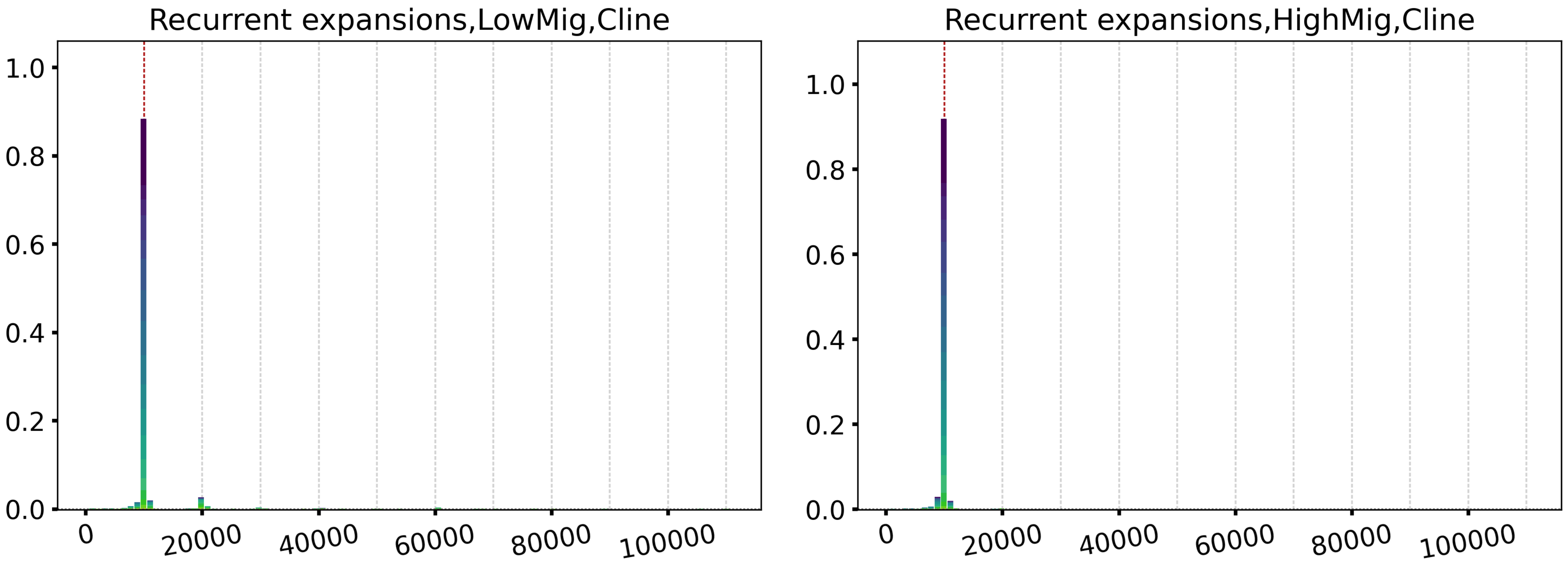

### High polygenicity

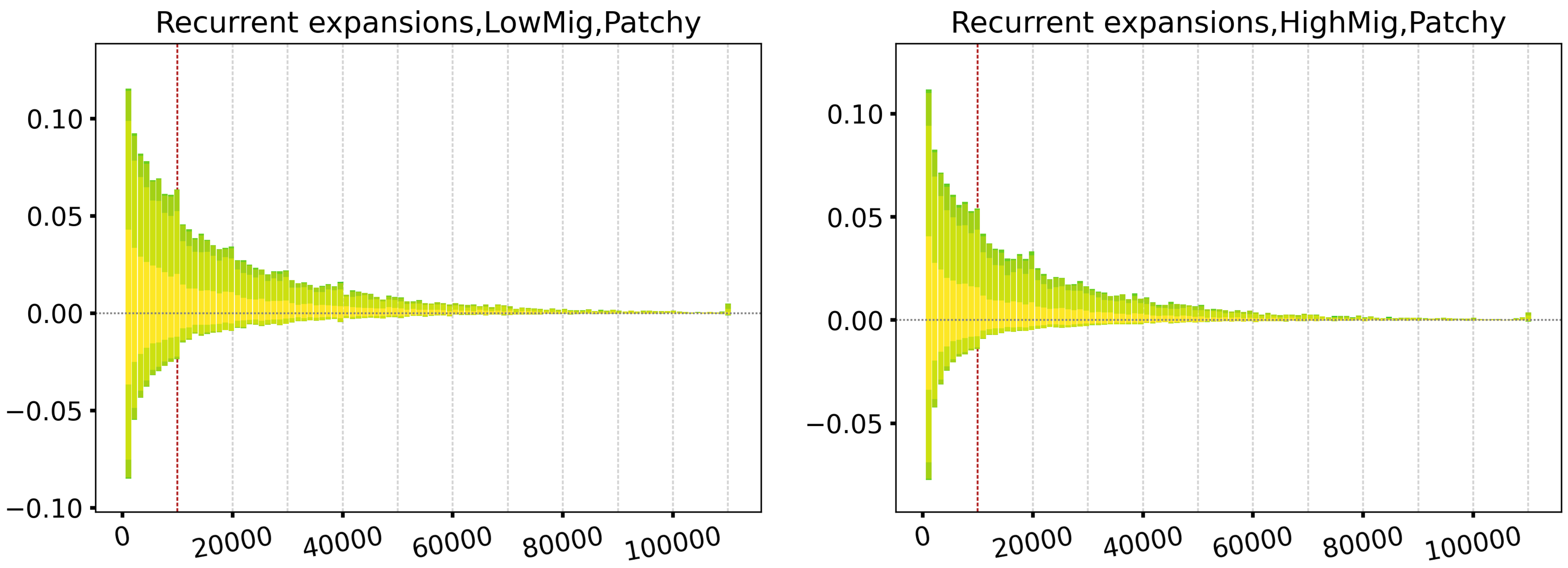


### Low polygenicity

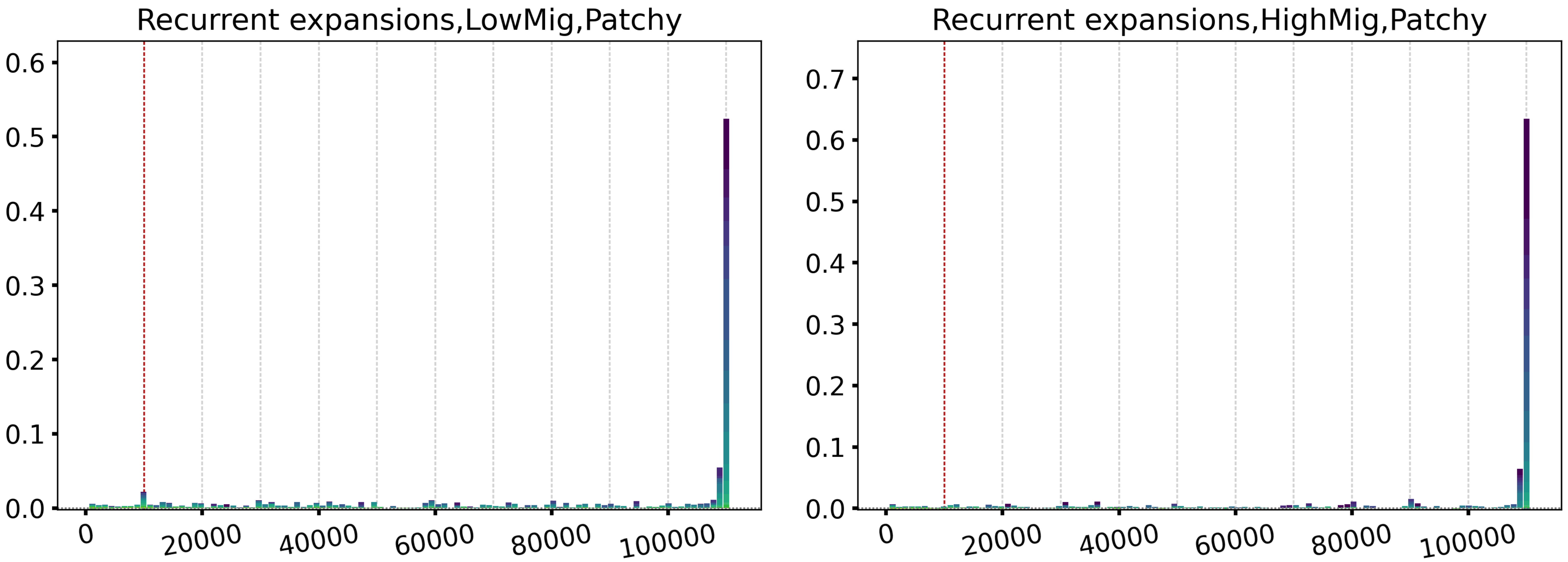

### High polygenicity

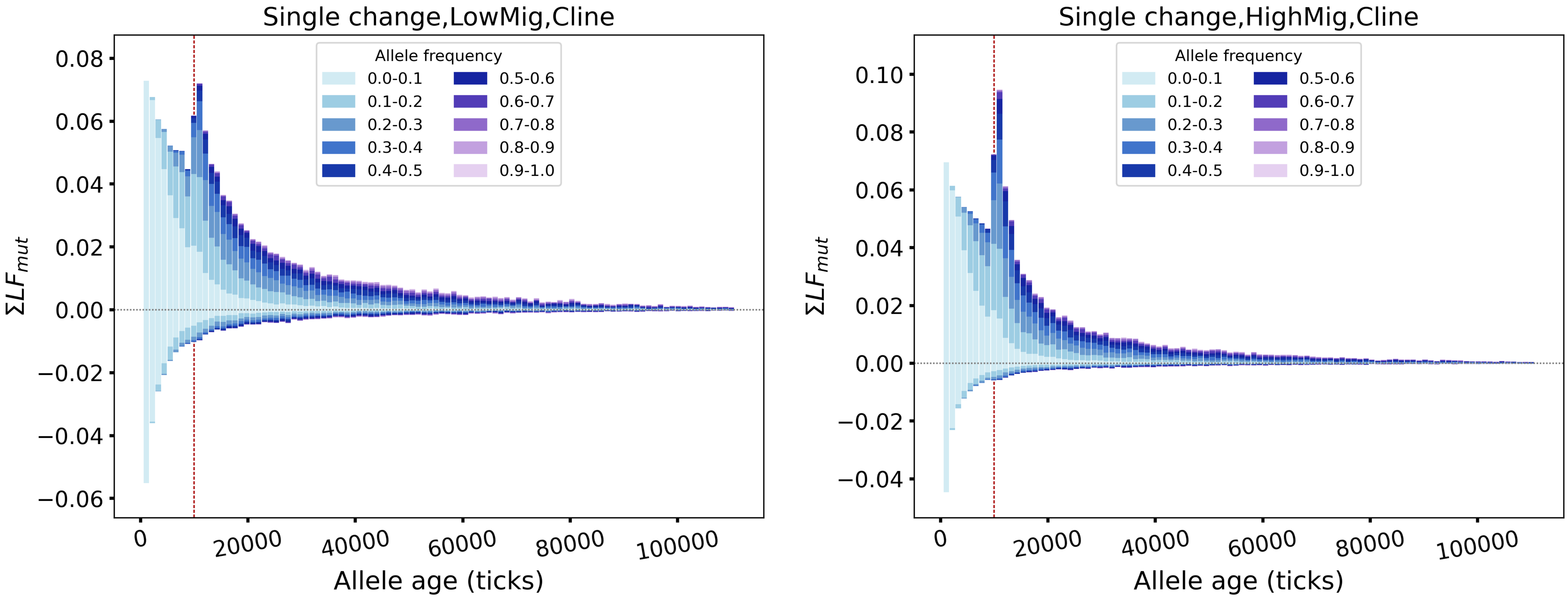


### Low polygenicity

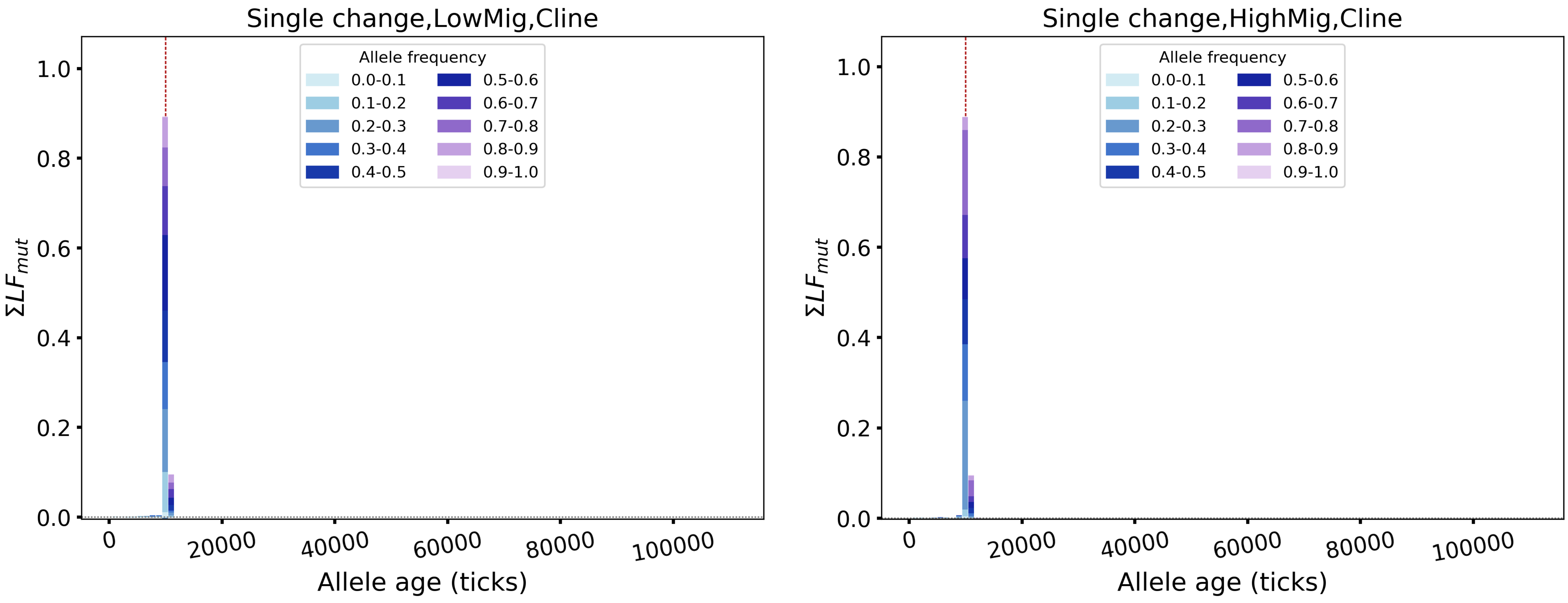

### High polygenicity

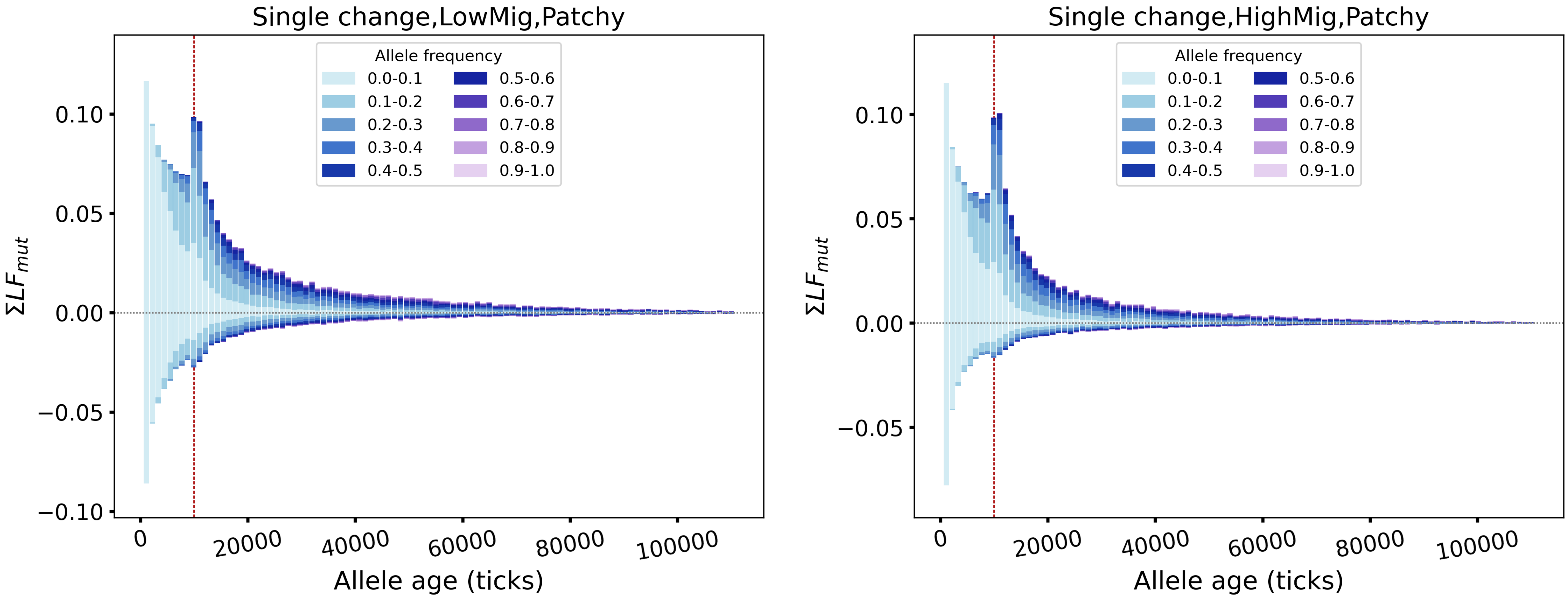


### Low polygenicity

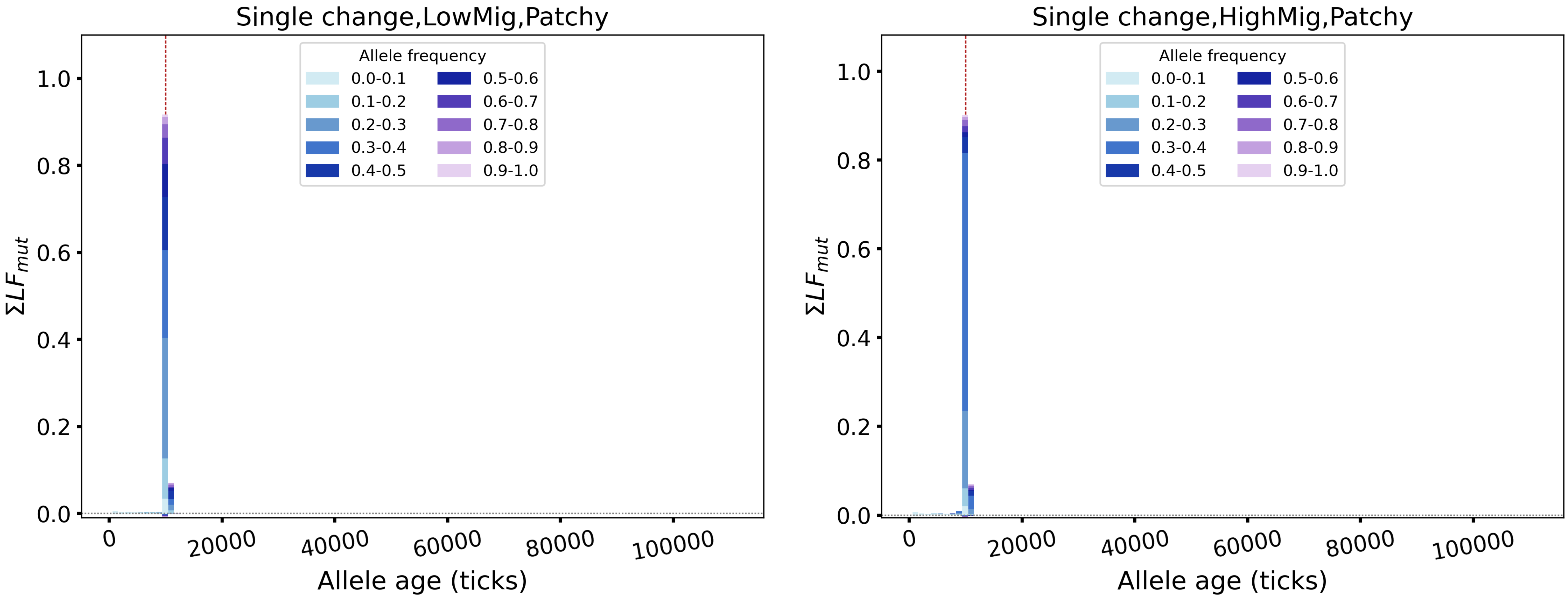

### High polygenicity

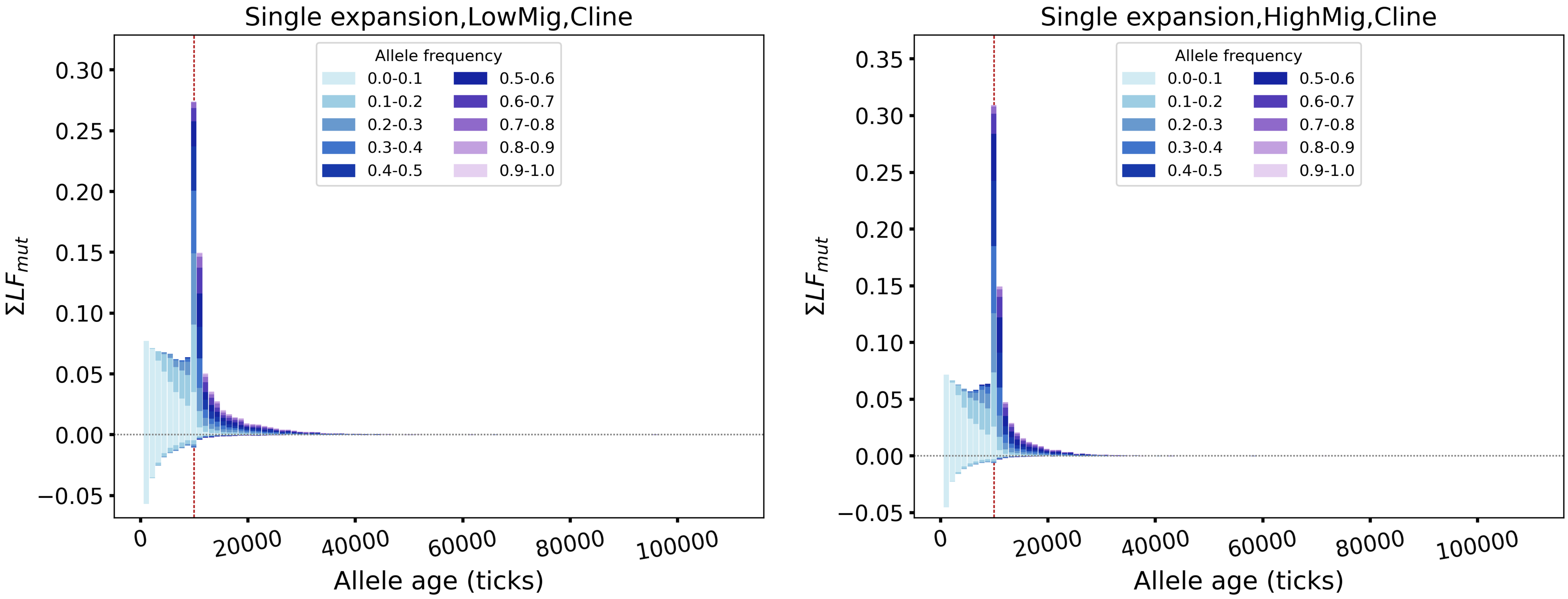


### Low polygenicity

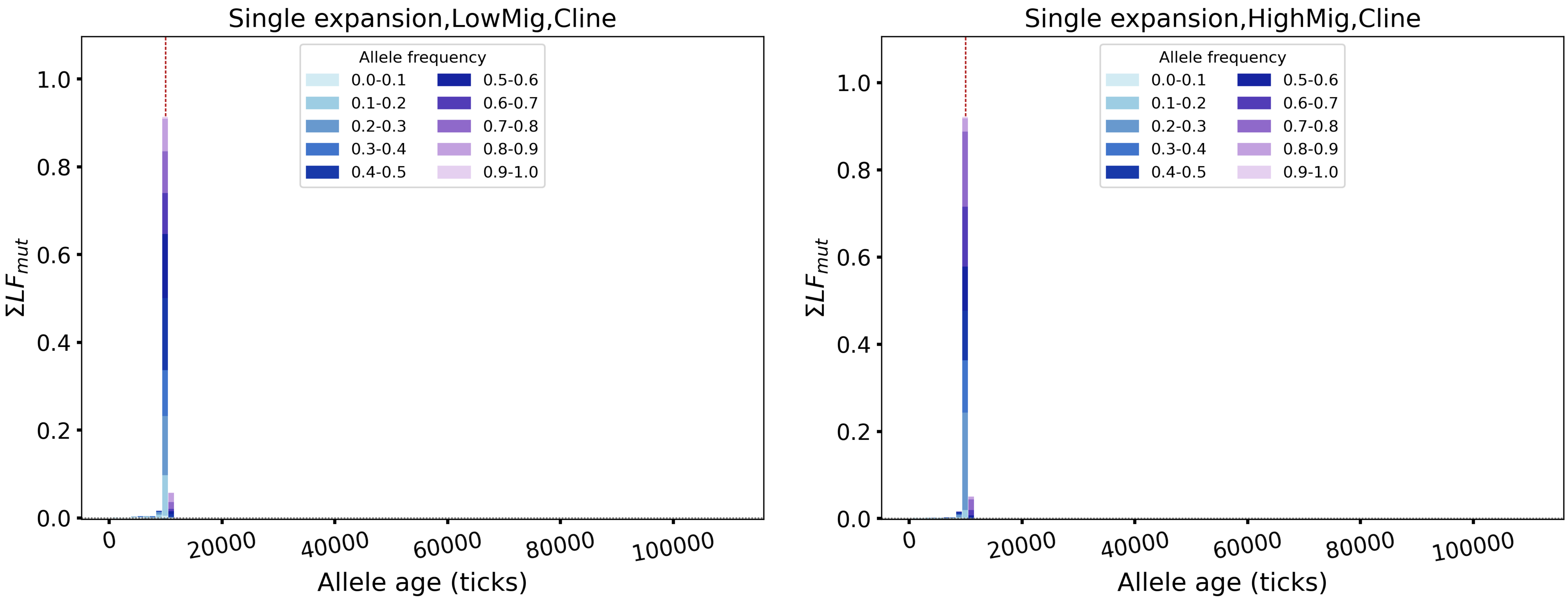

### High polygenicity

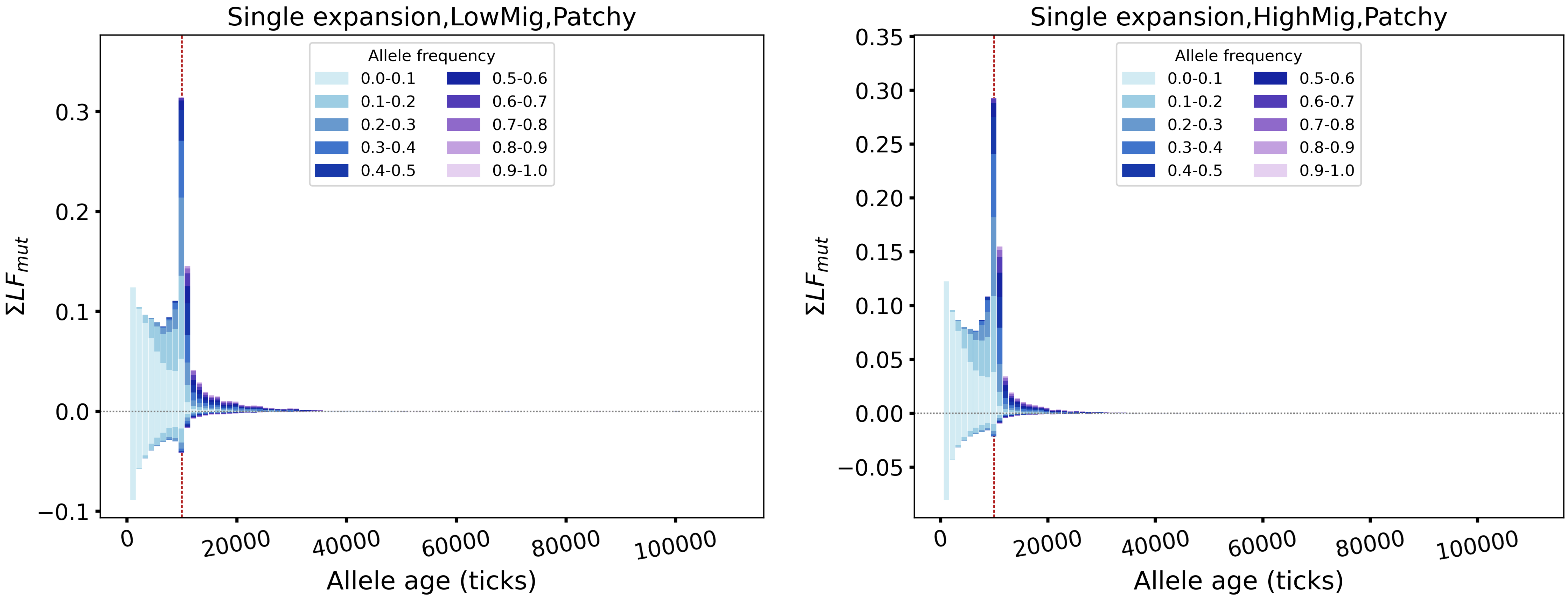


### Low polygenicity

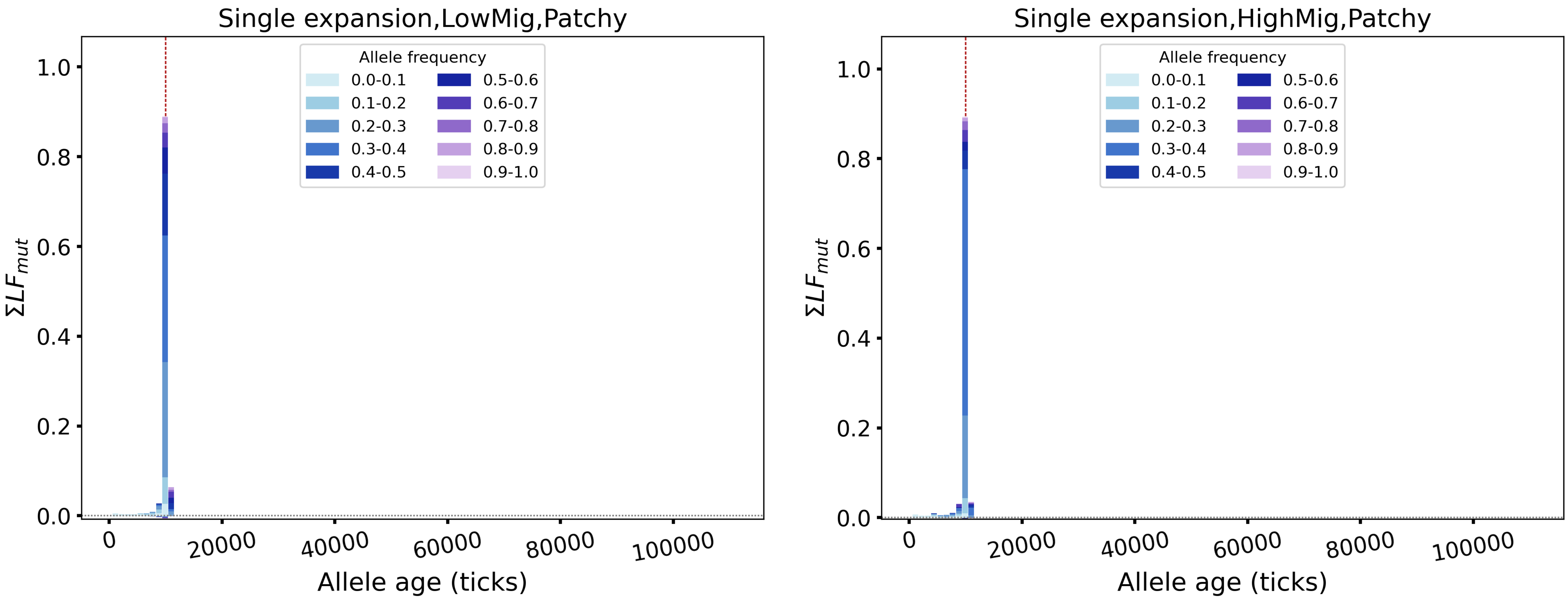

### High polygenicity

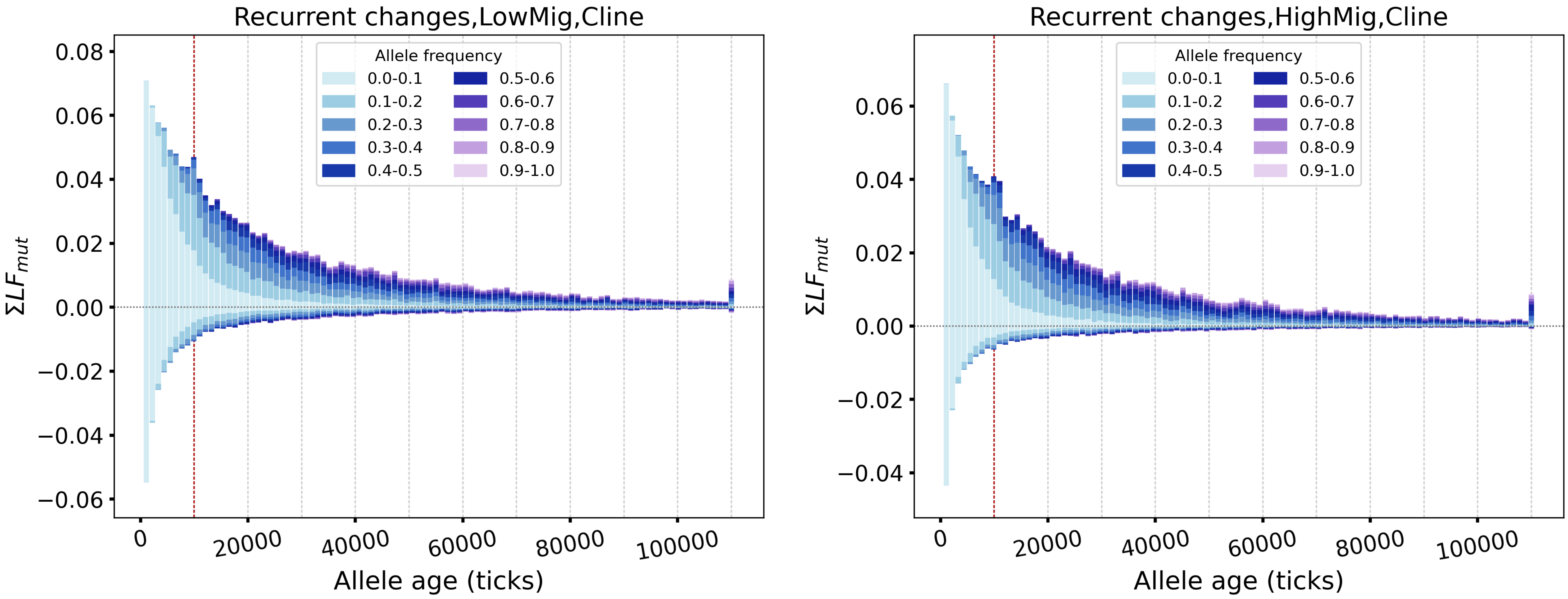


### Low polygenicity

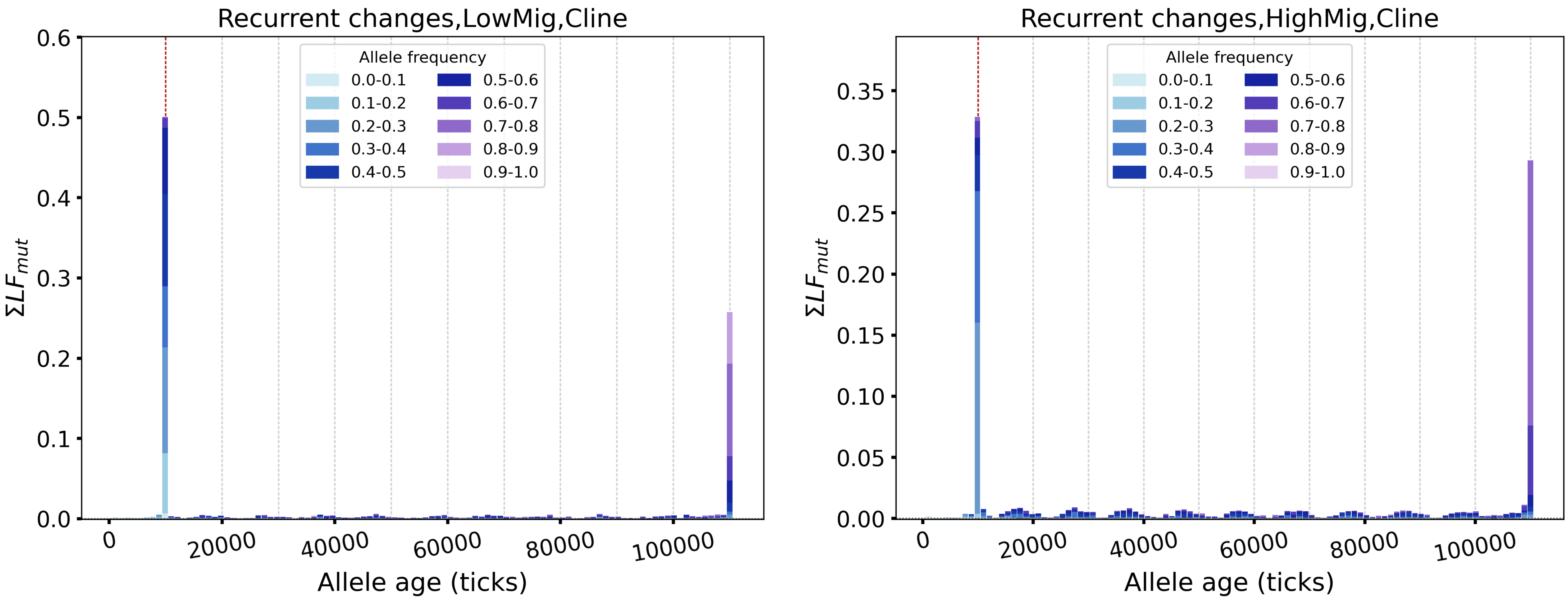

### High polygenicity

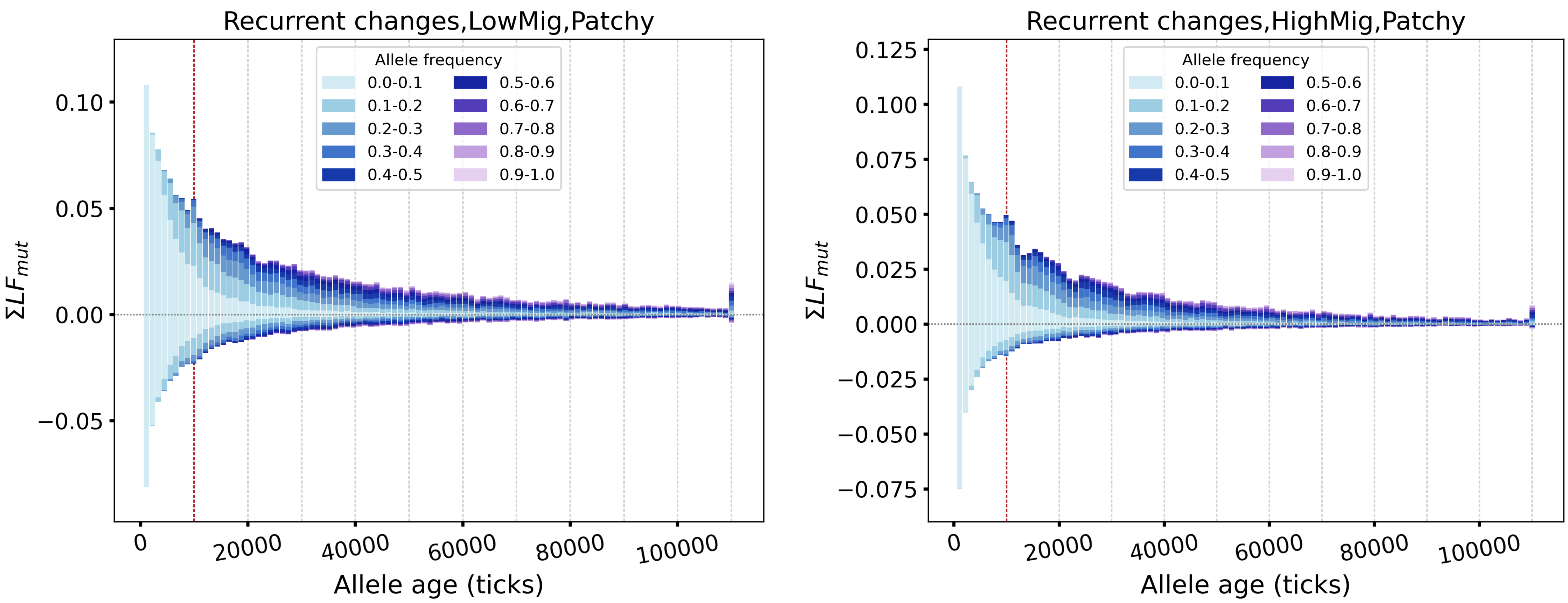


### Low polygenicity

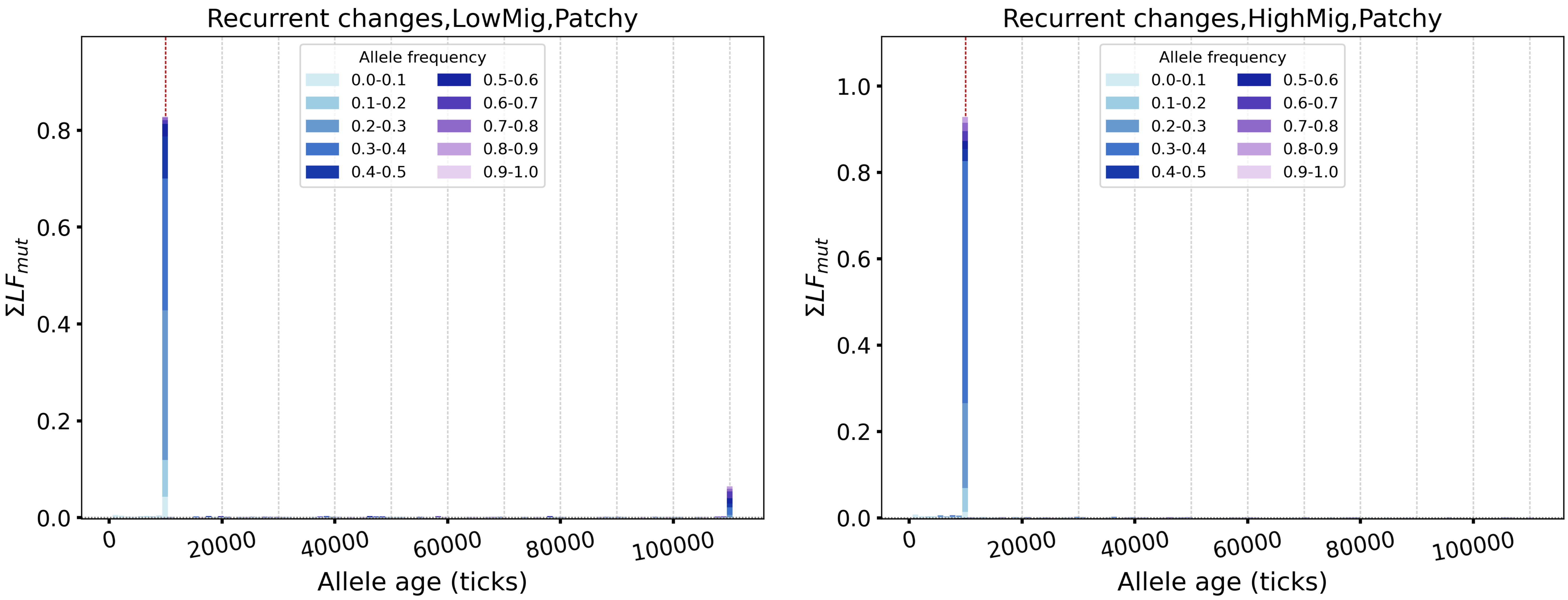

### High polygenicity

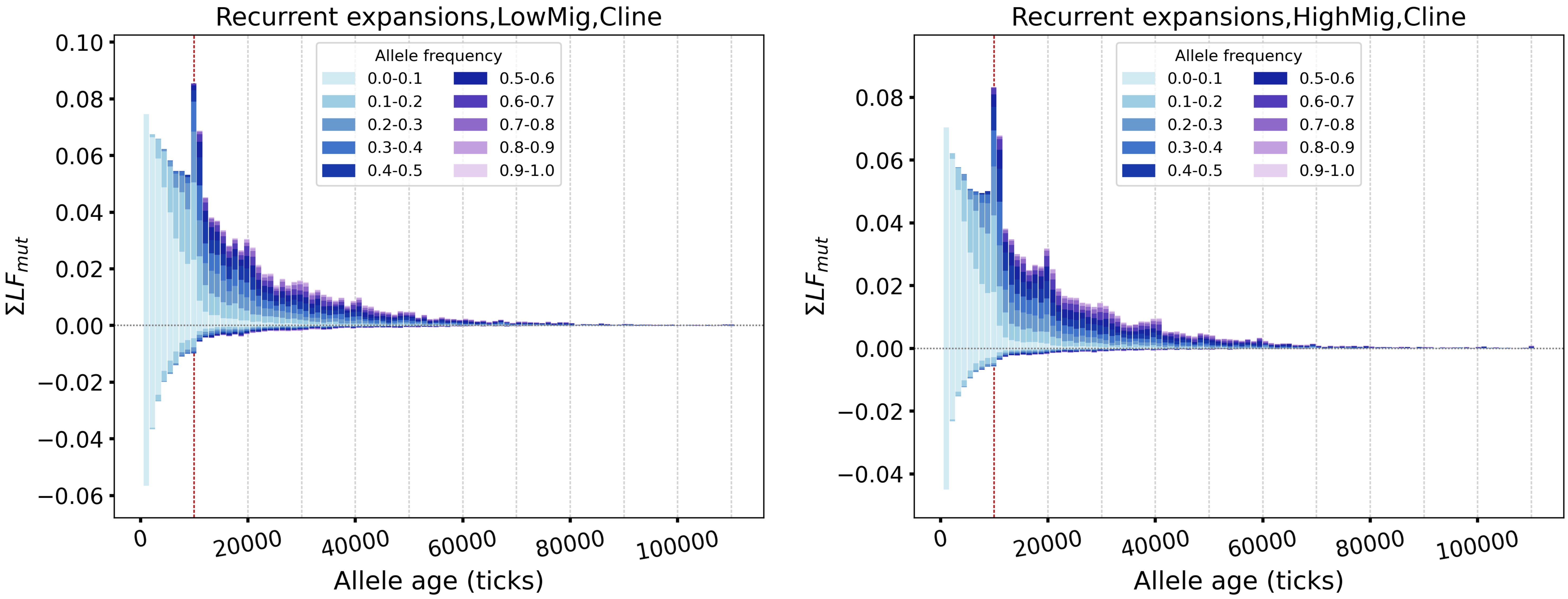


### Low polygenicity

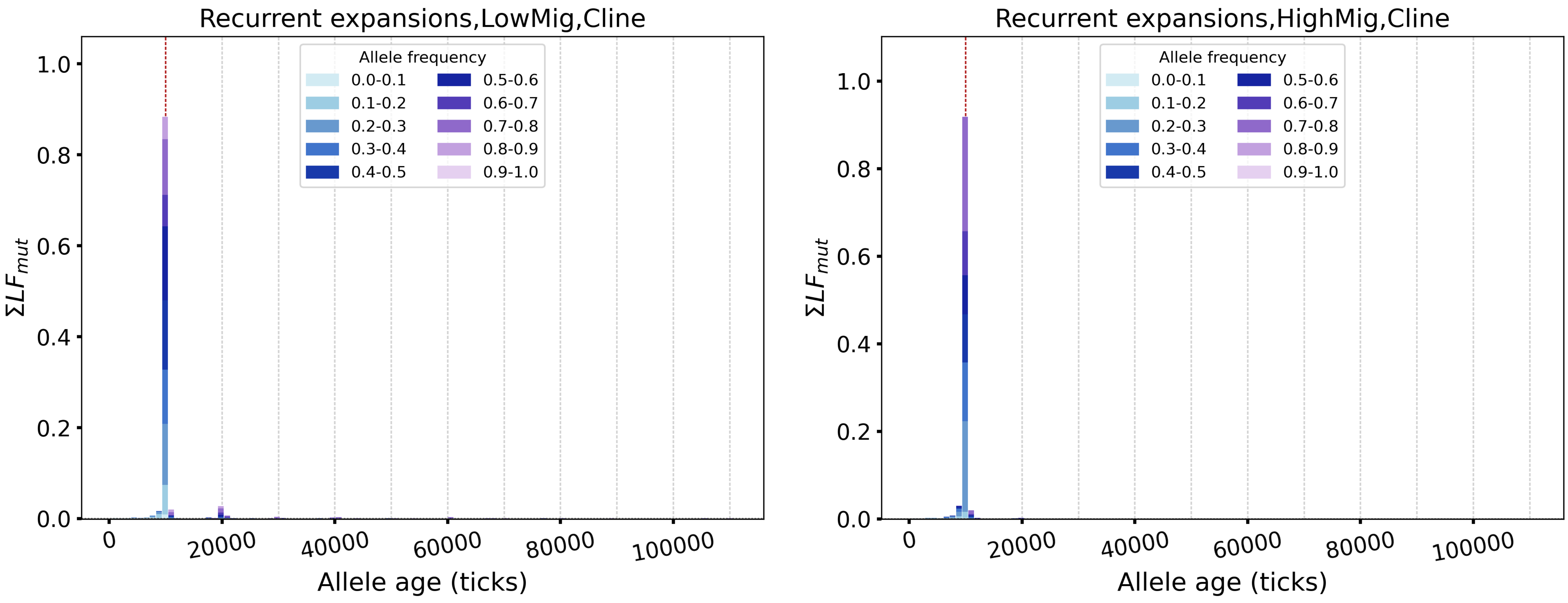

### High polygenicity

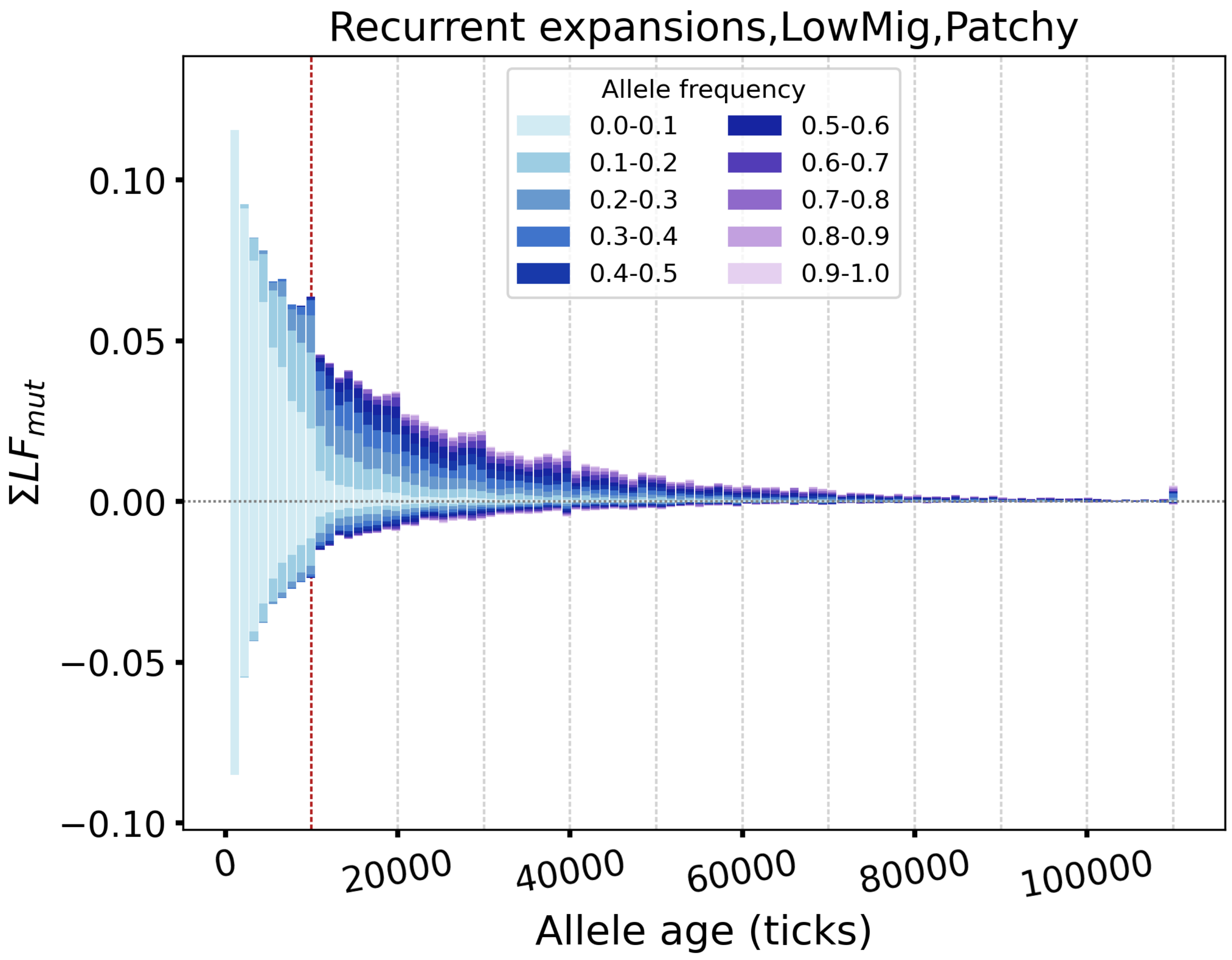


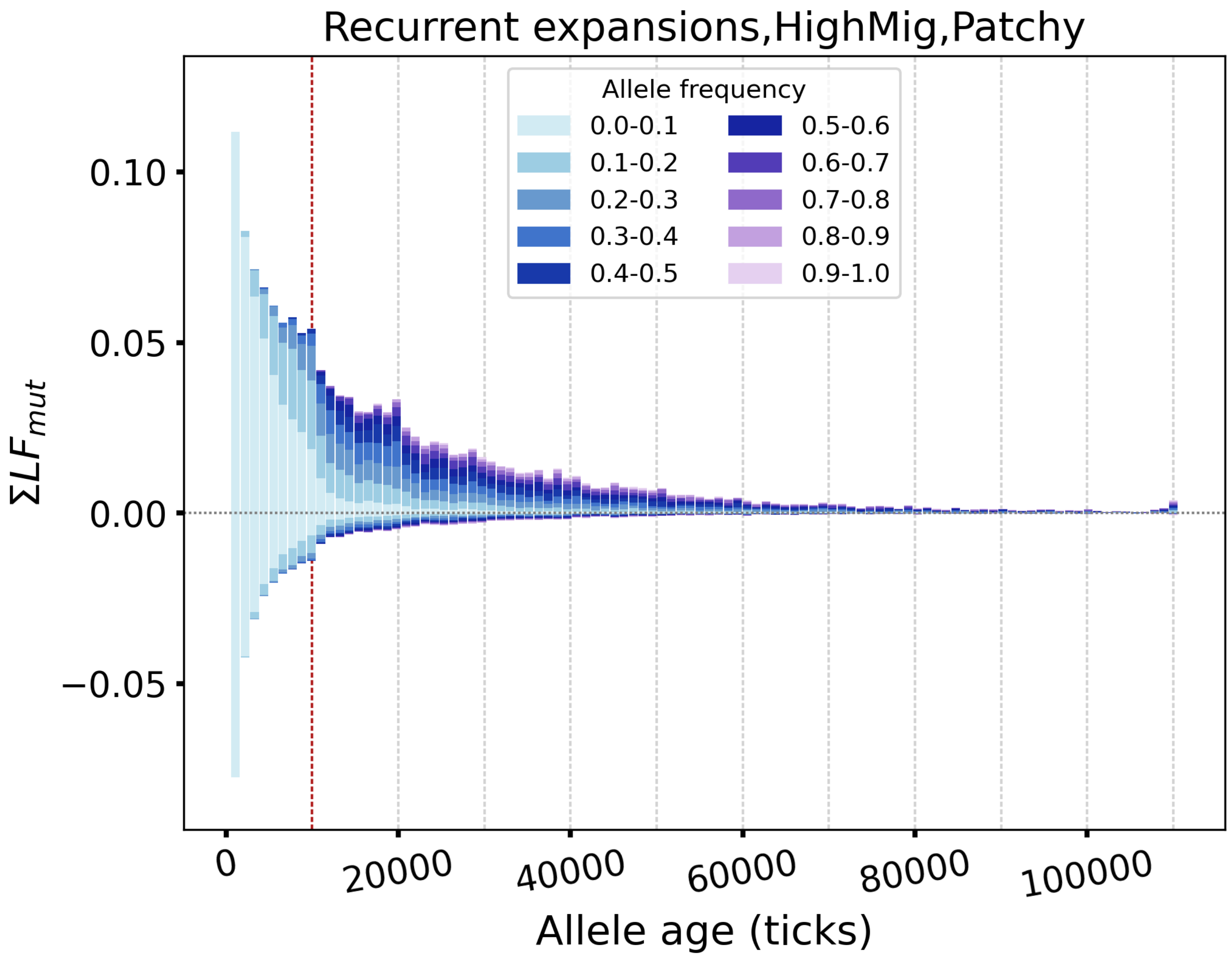


### Low polygenicity

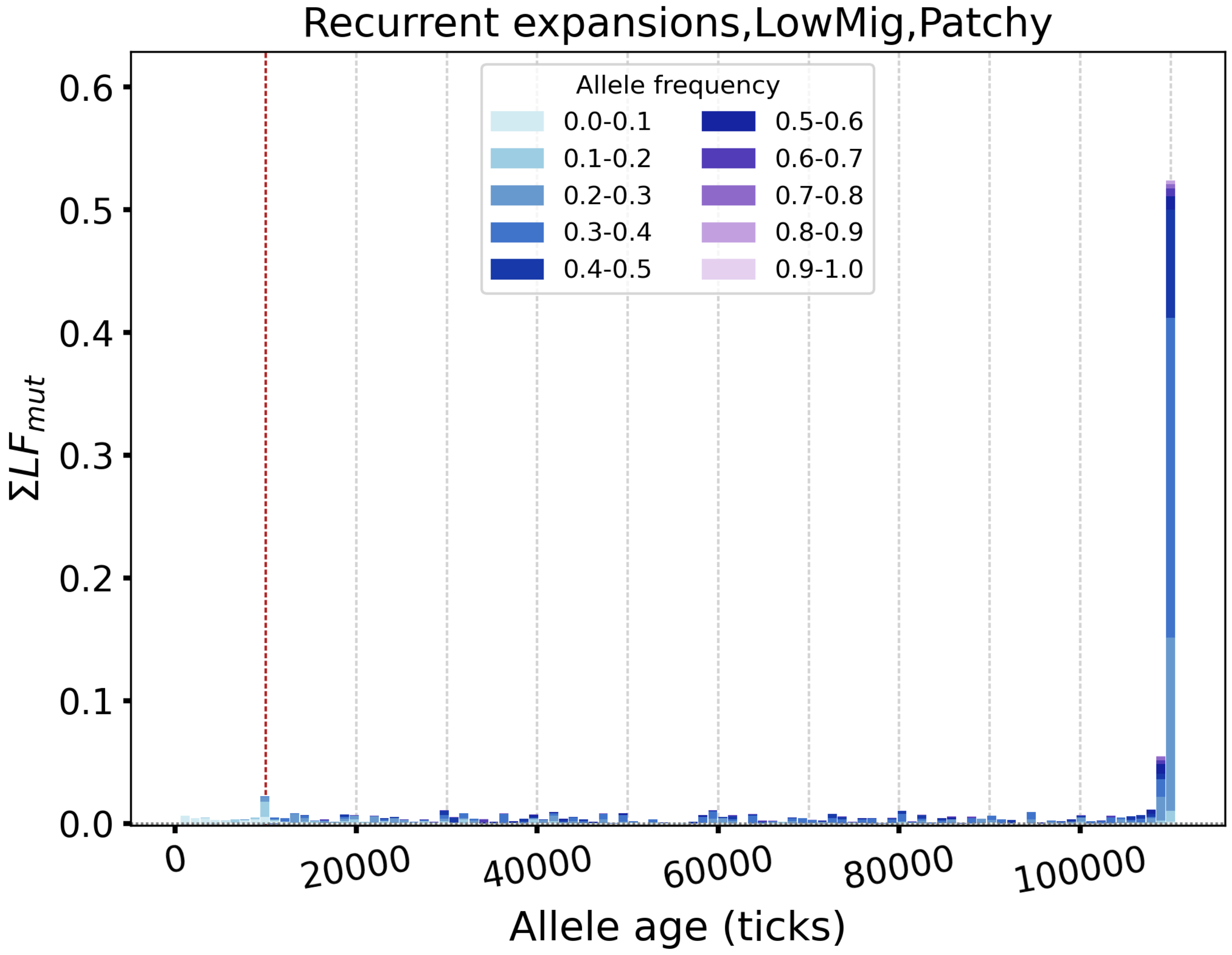


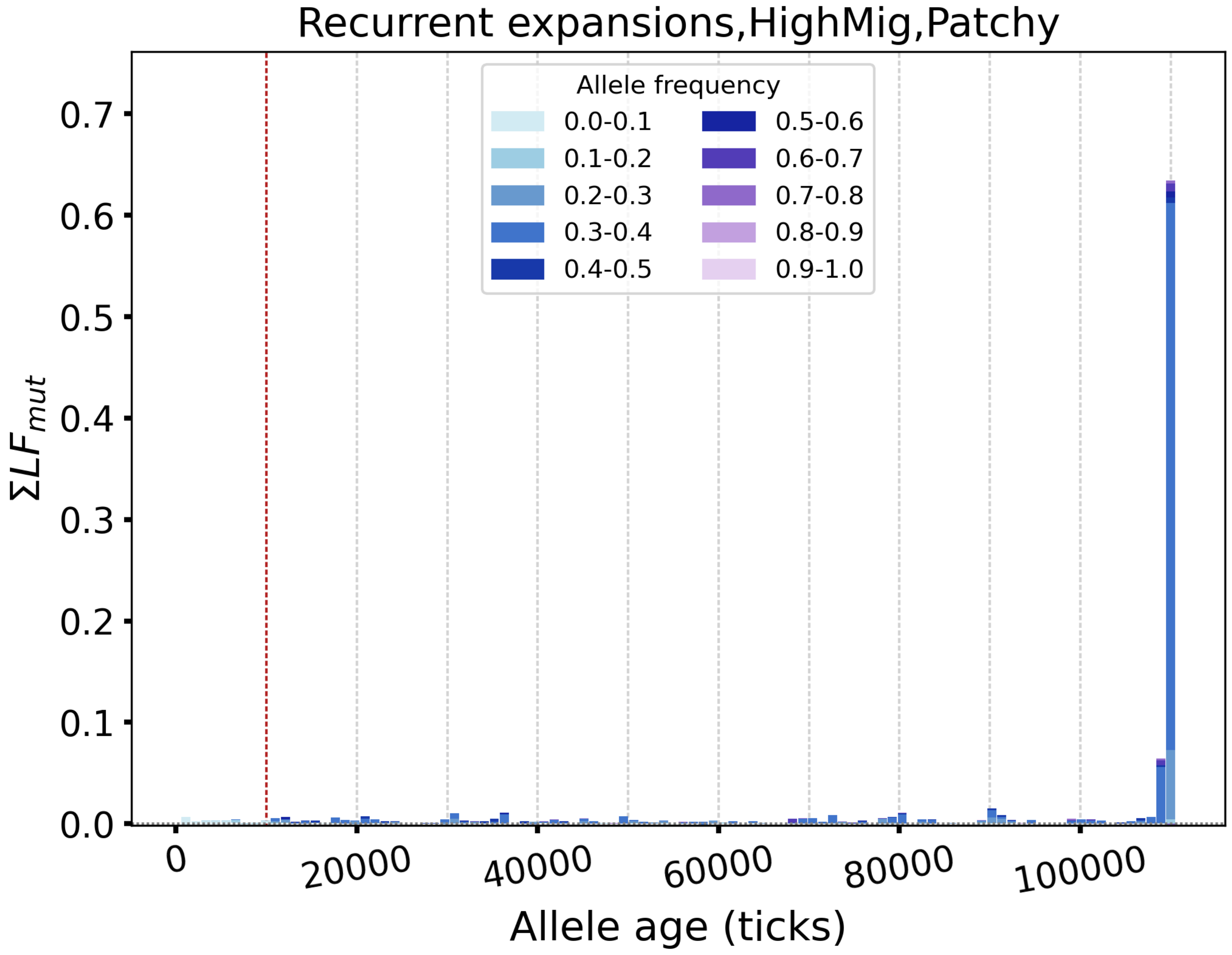
