## Supplementary Figure S7 for "The genetic architecture of local adaptation is historically contingent"

**Supplementary figure S7.** The relationship between allele age and false positive rate (FPR=false positives/(false positives + true negatives)) of neutral alleles in genotype-environment association analysis. The FPR of neutral alleles in each age category was calculated as the number of neutral alleles that had a p-value < $1 \times 10^{-10}$ in Kendall's rank correlation test divided by the total number of neutral alleles within each age category, using results of 200 simulation replicates. Alleles with minor allele frequency < 0.05 were excluded from the analysis. Allele ages are shown in thousand SLiM ticks. One tick is equivalent to a simulated year, in which all individuals reproduced once. High polygenicity: The standard deviation of the distribution of phenotypic effect size $\sigma_M = 0.01$, and functional mutation rate $\mu_f = 1 \times 10^{-8}$per base pair per simulated year; Low polygenicity : $\sigma_M = 0.1$, $\mu_f = 1 \times 10^{-10}$; Low-Mig./High-Mig.: a lower/higher migration level (corresponding to average overall $F_{ST}$ values of 0.112$\pm$0.008 and 0.038$\pm$0.002, respectively).

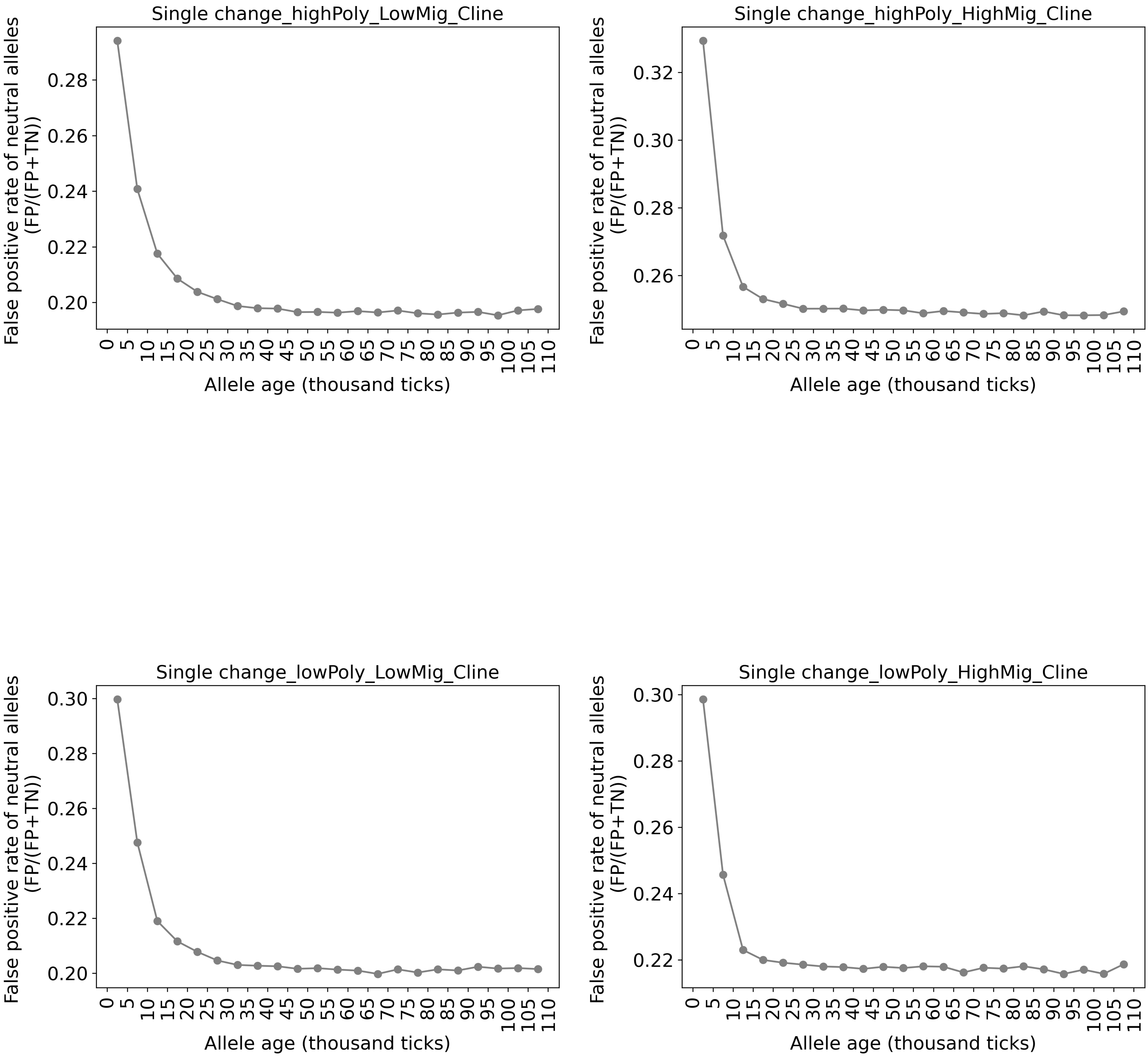
Single change_highPoly_LowMig_Cline
Single change_highPoly_HighMig_Cline
Single change_lowPoly_LowMig_Cline
Single change_lowPoly_HighMig_Cline
False positive rate of neutral alleles (FP/(FP+TN))
Allele age (thousand ticks)

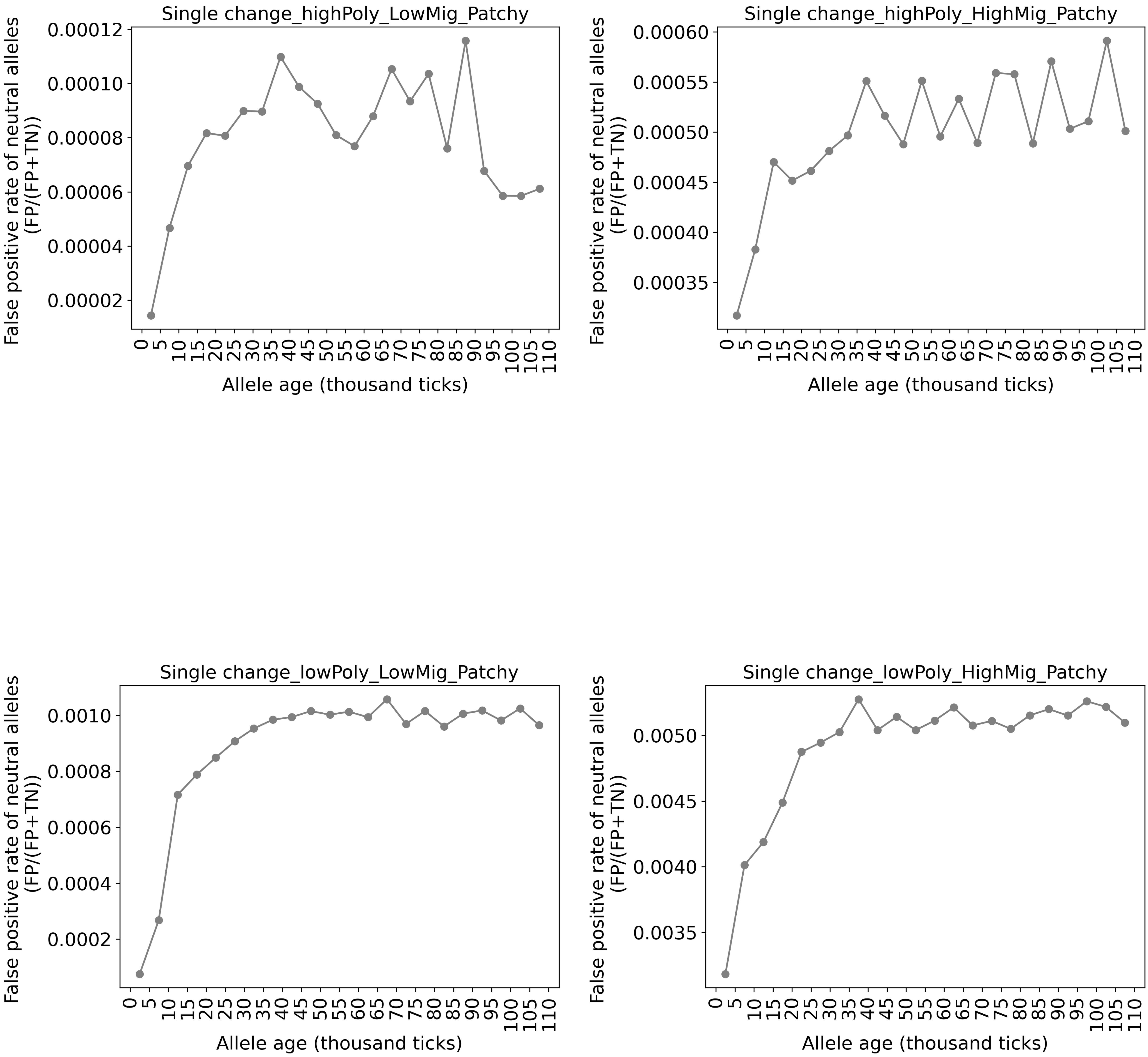
Single change_highPoly_LowMig_Patchy
False positive rate of neutral alleles (FP/(FP+TN))
0.00012
0.00010
0.00008
0.00006
0.00004
0.00002
0 5 10 15 20 25 30 35 40 45 50 55 60 65 70 75 80 85 90 95 100 105 110
Allele age (thousand ticks)
Single change_highPoly_HighMig_Patchy
False positive rate of neutral alleles (FP/(FP+TN))
0.00060
0.00055
0.00050
0.00045
0.00040
0.00035
0 5 10 15 20 25 30 35 40 45 50 55 60 65 70 75 80 85 90 95 100 105 110
Allele age (thousand ticks)
Single change_lowPoly_LowMig_Patchy
False positive rate of neutral alleles (FP/(FP+TN))
0.0010
0.0008
0.0006
0.0004
0.0002
0 5 10 15 20 25 30 35 40 45 50 55 60 65 70 75 80 85 90 95 100 105 110
Allele age (thousand ticks)
Single change_lowPoly_HighMig_Patchy
False positive rate of neutral alleles (FP/(FP+TN))
0.0050
0.0045
0.0040
0.0035
0 5 10 15 20 25 30 35 40 45 50 55 60 65 70 75 80 85 90 95 100 105 110
Allele age (thousand ticks)

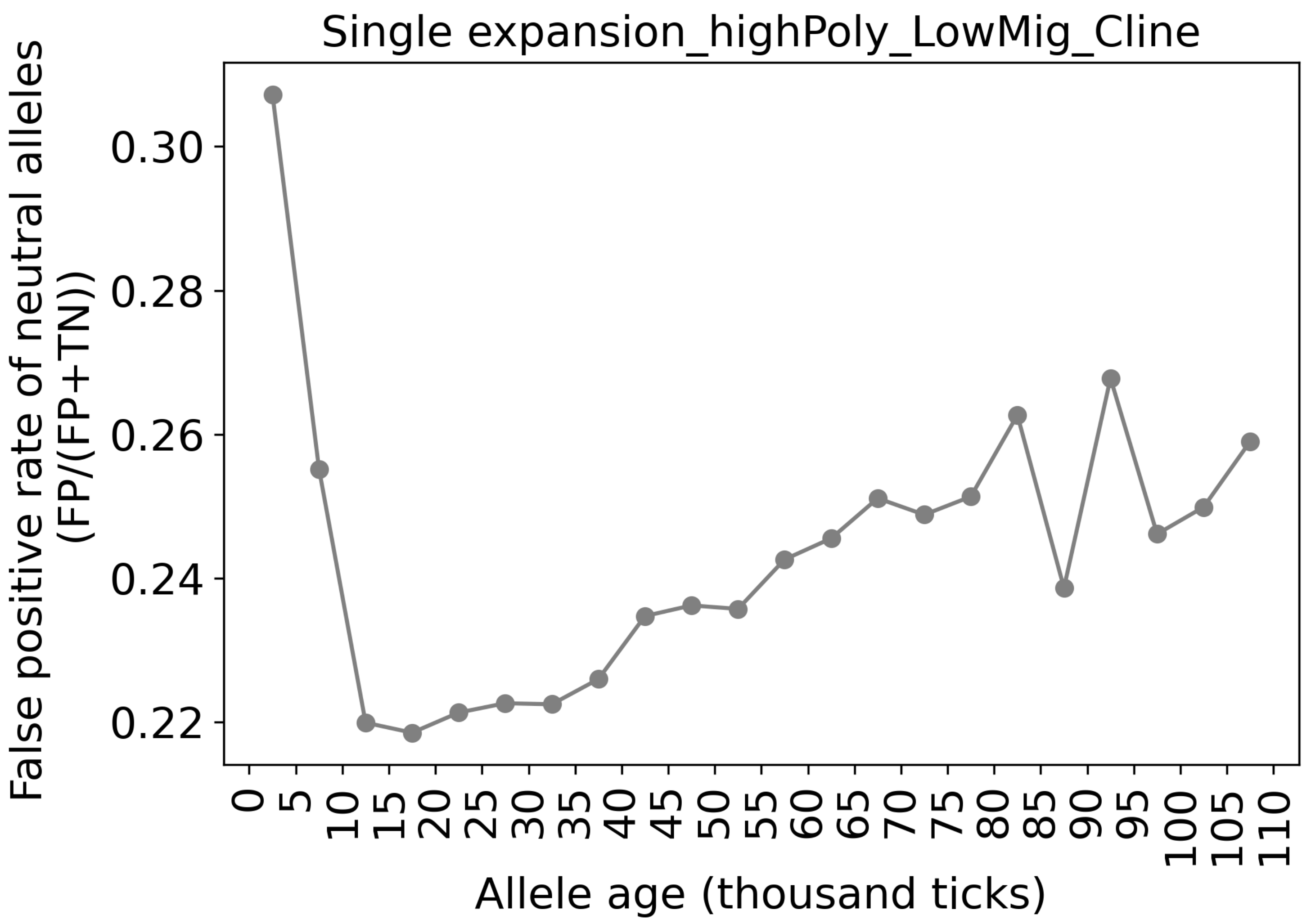
Single expansion_highPoly_LowMig_Cline
False positive rate of neutral alleles
(FP/(FP+TN))
Allele age (thousand ticks)

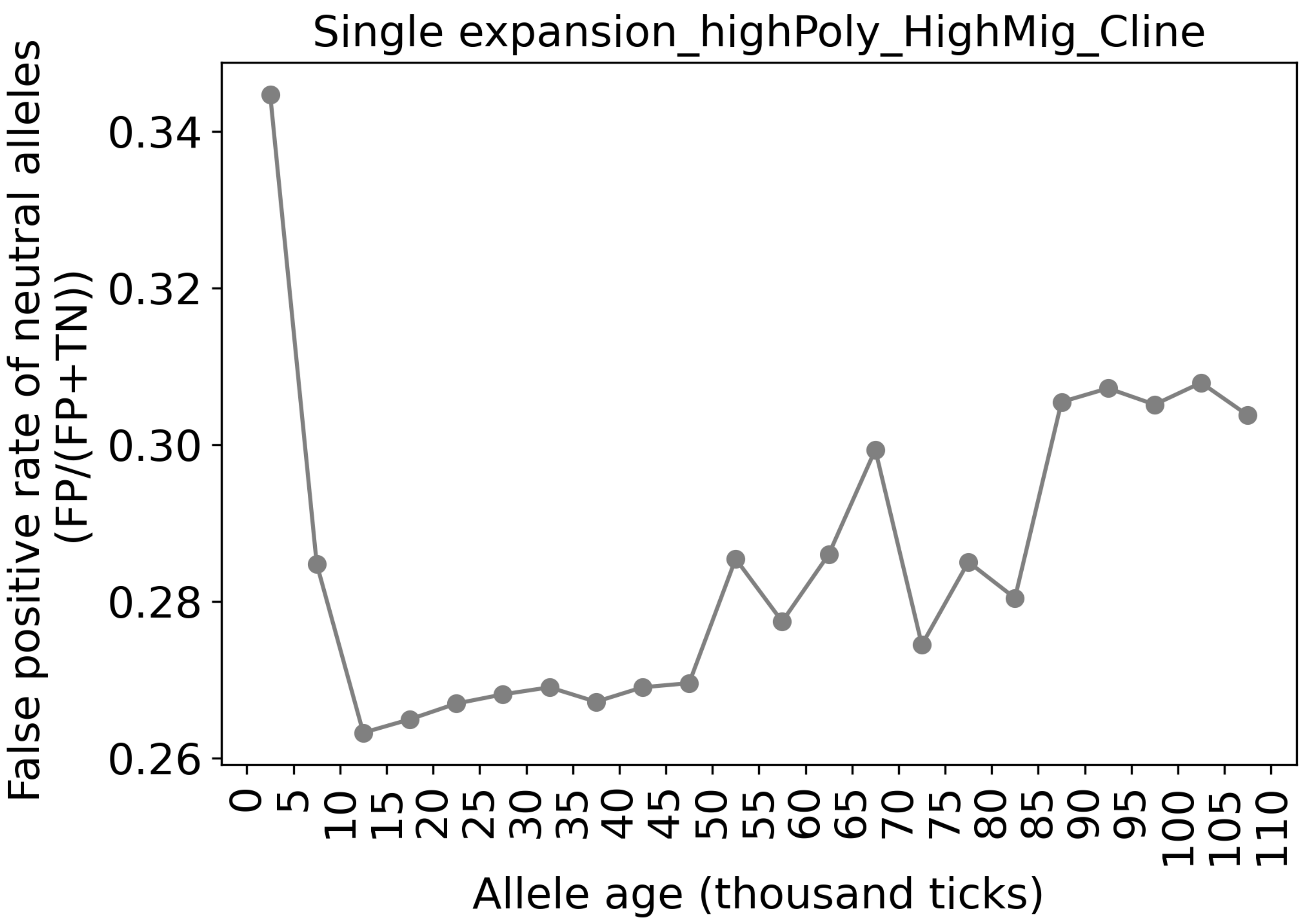
Single expansion_highPoly_HighMig_Cline
False positive rate of neutral alleles
(FP/(FP+TN))
Allele age (thousand ticks)

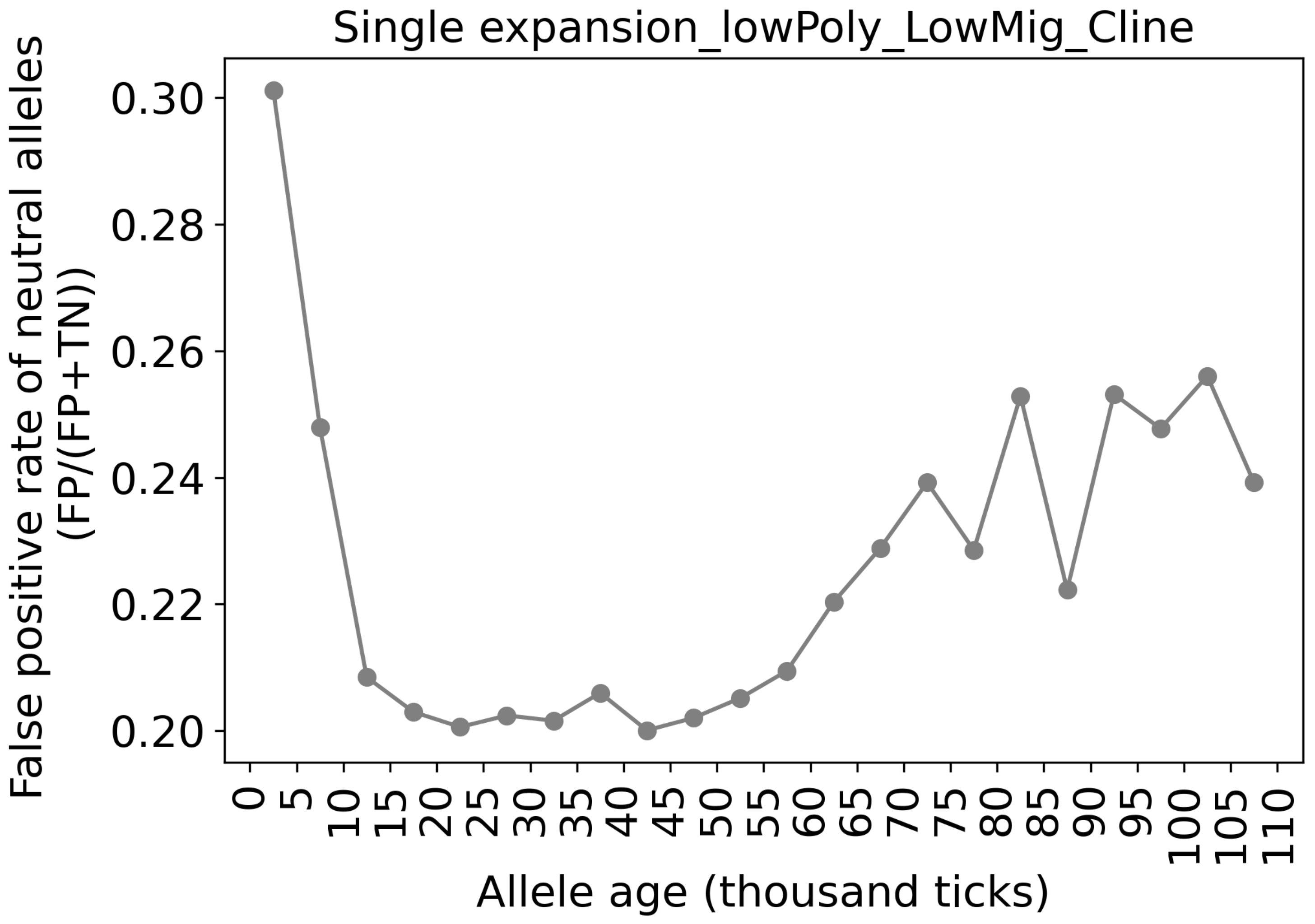
Single expansion_lowPoly_LowMig_Cline
False positive rate of neutral alleles
(FP/(FP+TN))
Allele age (thousand ticks)

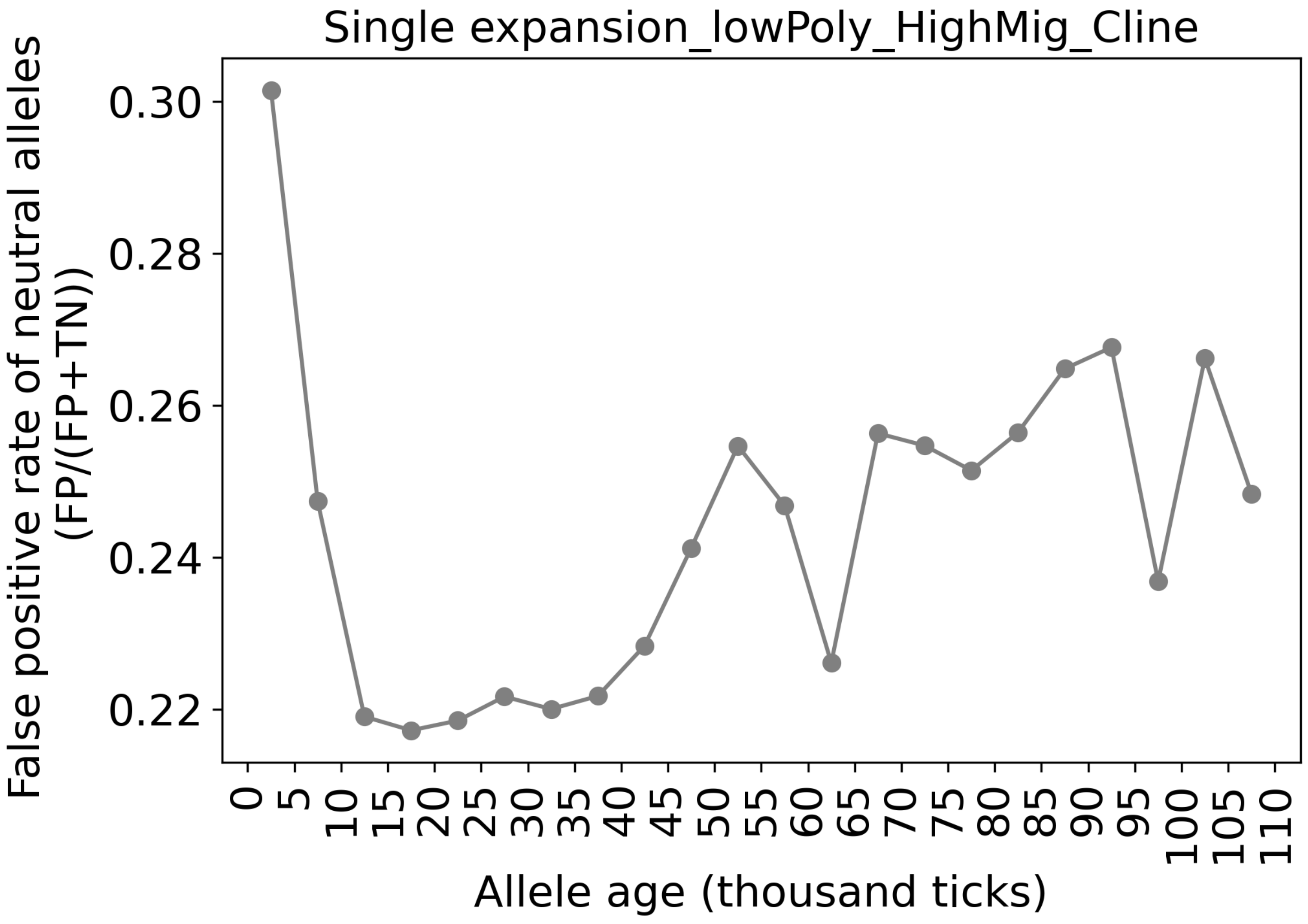
Single expansion_lowPoly_HighMig_Cline
False positive rate of neutral alleles
(FP/(FP+TN))
Allele age (thousand ticks)

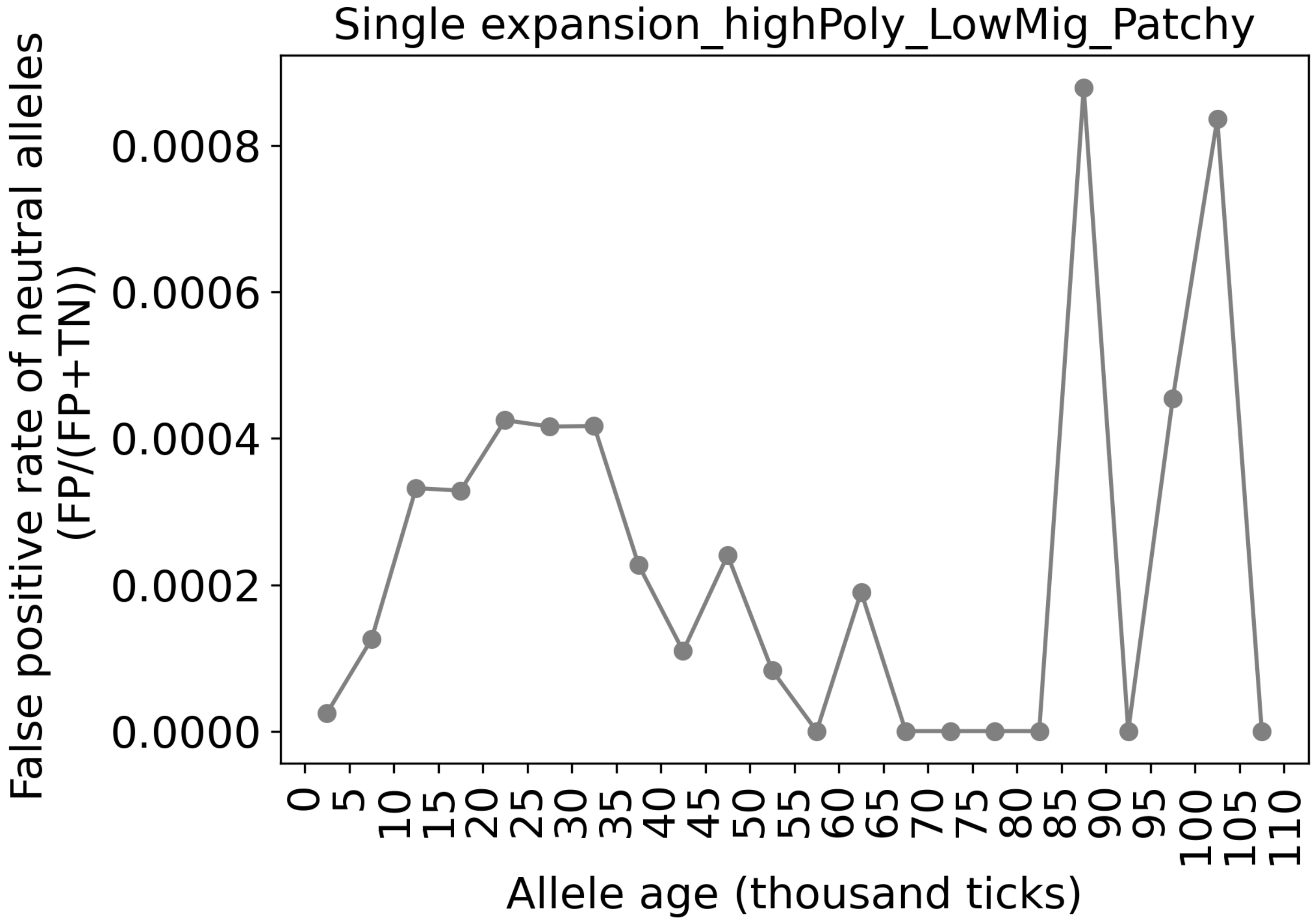
Single expansion_highPoly_LowMig_Patchy
False positive rate of neutral alleles (FP/(FP+TN))
0.0000
0.0002
0.0004
0.0006
0.0008
0 5 10 15 20 25 30 35 40 45 50 55 60 65 70 75 80 85 90 95 100 105 110
Allele age (thousand ticks)

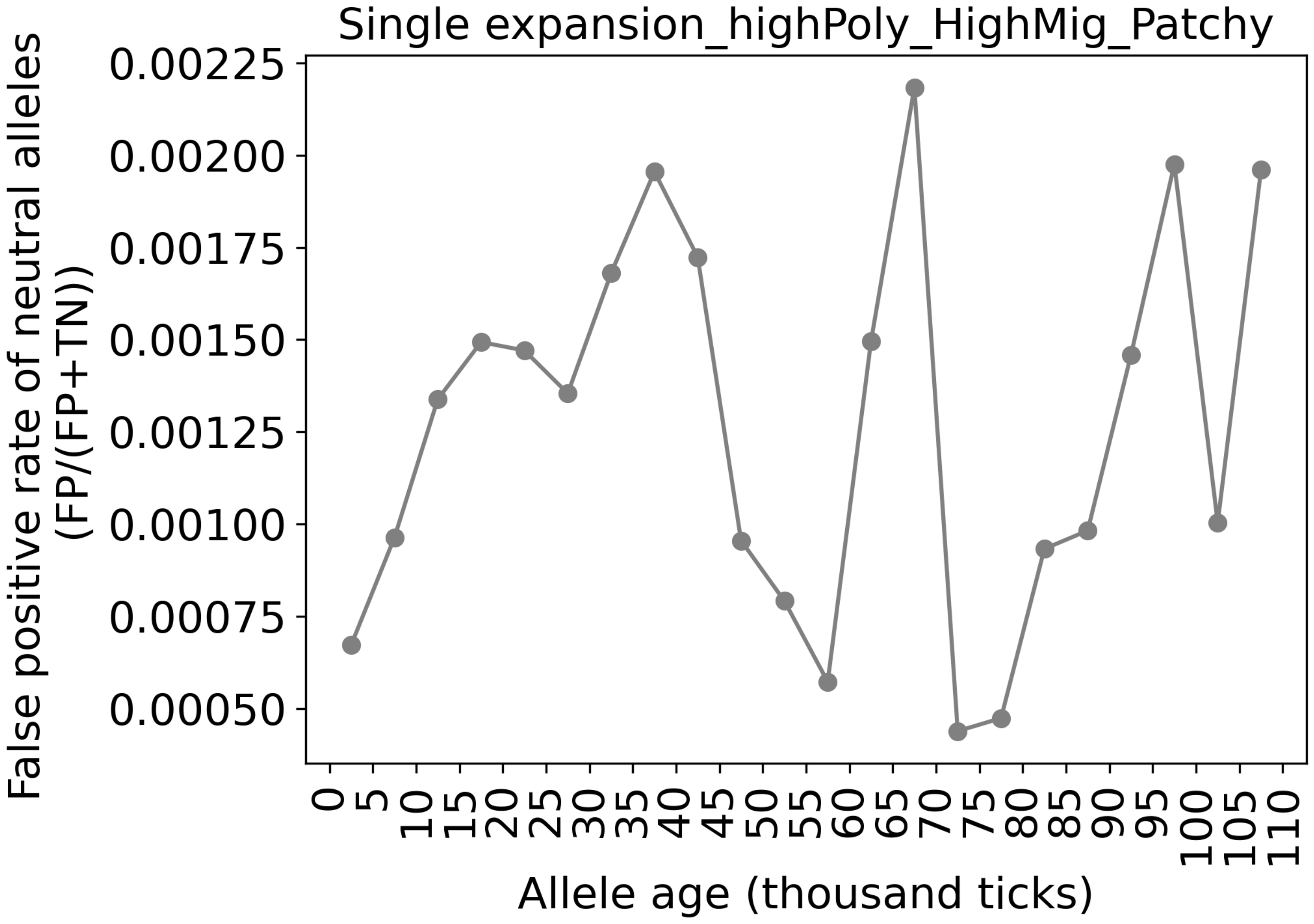
Single expansion_highPoly_HighMig_Patchy
False positive rate of neutral alleles (FP/(FP+TN))
0.00050
0.00075
0.00100
0.00125
0.00150
0.00175
0.00200
0.00225
0 5 10 15 20 25 30 35 40 45 50 55 60 65 70 75 80 85 90 95 100 105 110
Allele age (thousand ticks)

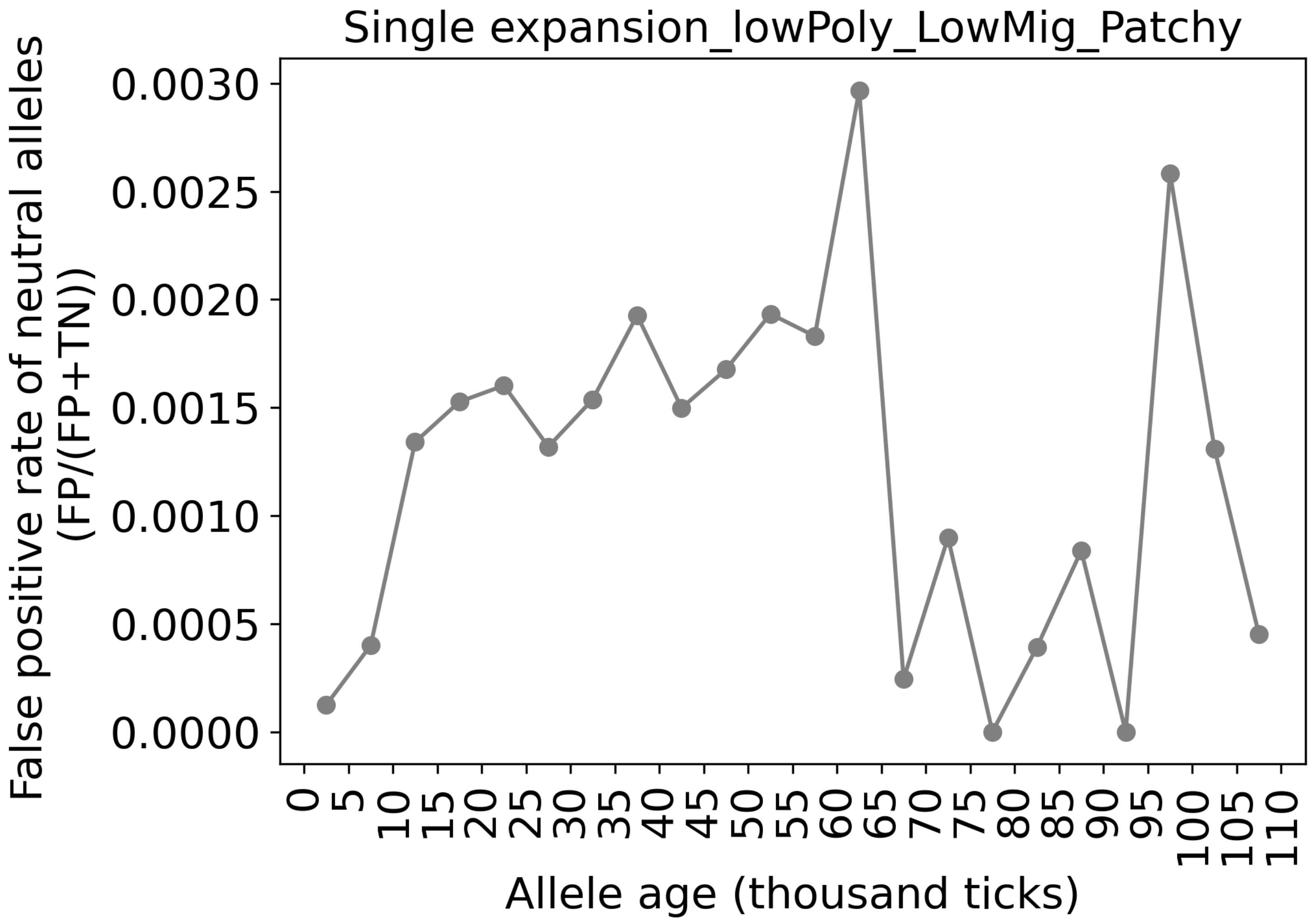
Single expansion_lowPoly_LowMig_Patchy
False positive rate of neutral alleles (FP/(FP+TN))
0.0000
0.0005
0.0010
0.0015
0.0020
0.0025
0.0030
0 5 10 15 20 25 30 35 40 45 50 55 60 65 70 75 80 85 90 95 100 105 110
Allele age (thousand ticks)

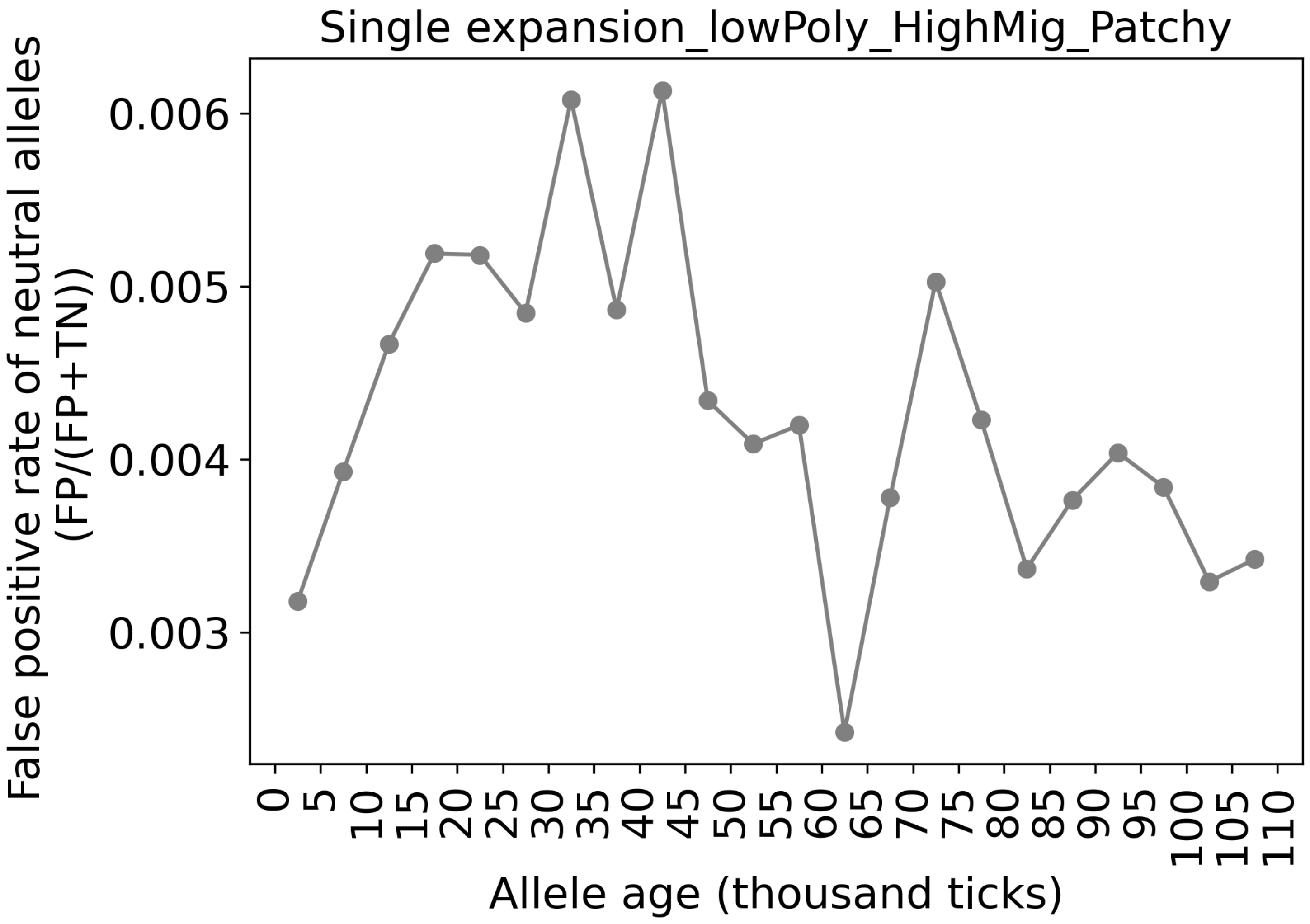
Single expansion_lowPoly_HighMig_Patchy
False positive rate of neutral alleles (FP/(FP+TN))
0.003
0.004
0.005
0.006
0 5 10 15 20 25 30 35 40 45 50 55 60 65 70 75 80 85 90 95 100 105 110
Allele age (thousand ticks)

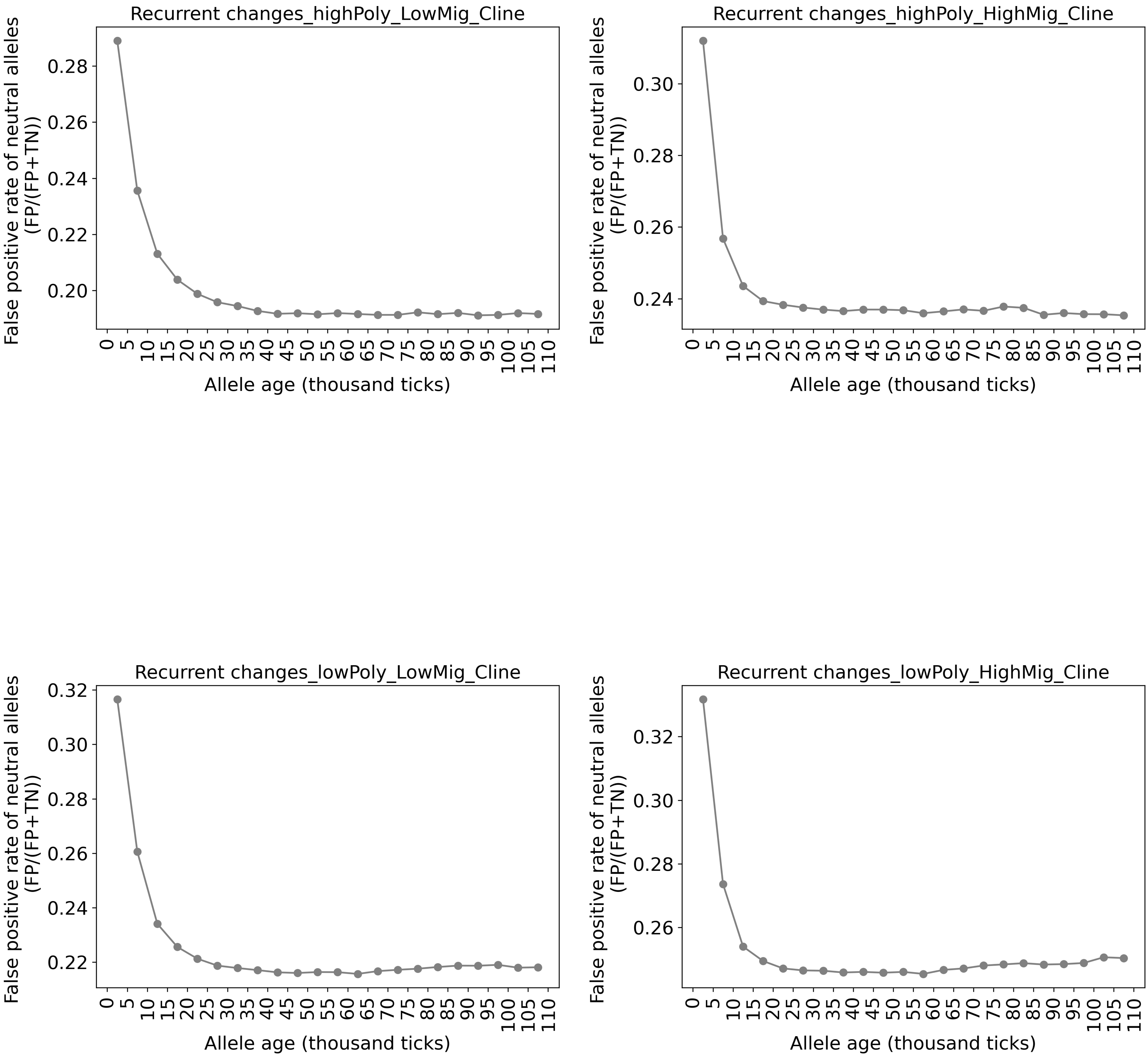
Recurrent changes_highPoly_LowMig_Cline
Recurrent changes_highPoly_HighMig_Cline
Recurrent changes_lowPoly_LowMig_Cline
Recurrent changes_lowPoly_HighMig_Cline
False positive rate of neutral alleles (FP/(FP+TN))
Allele age (thousand ticks)

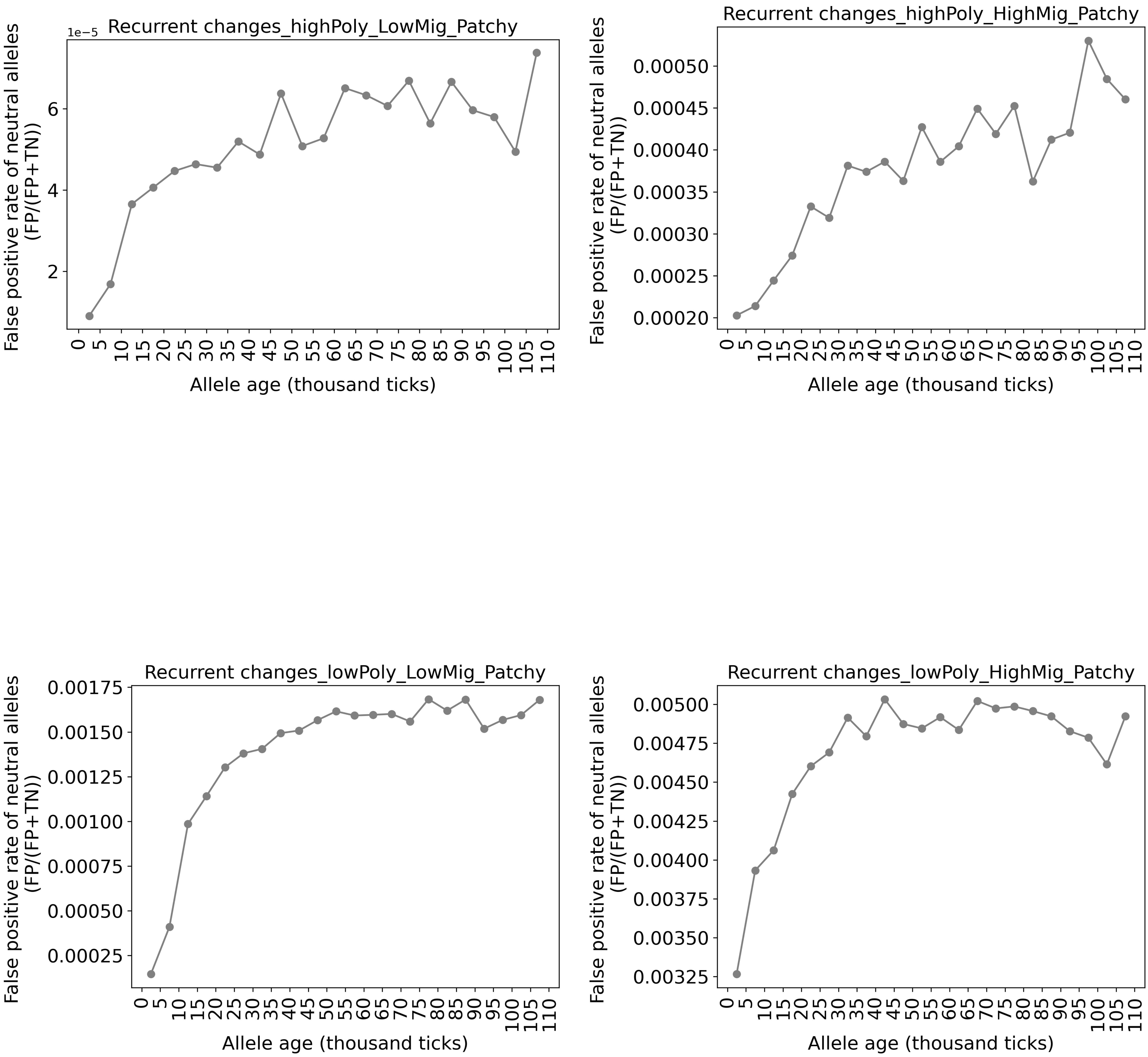
Recurrent changes_highPoly_LowMig_Patchy
1e−5
False positive rate of neutral alleles (FP/(FP+TN))
Allele age (thousand ticks)
Recurrent changes_highPoly_HighMig_Patchy
False positive rate of neutral alleles (FP/(FP+TN))
Allele age (thousand ticks)
Recurrent changes_lowPoly_LowMig_Patchy
False positive rate of neutral alleles (FP/(FP+TN))
Allele age (thousand ticks)
Recurrent changes_lowPoly_HighMig_Patchy
False positive rate of neutral alleles (FP/(FP+TN))
Allele age (thousand ticks)

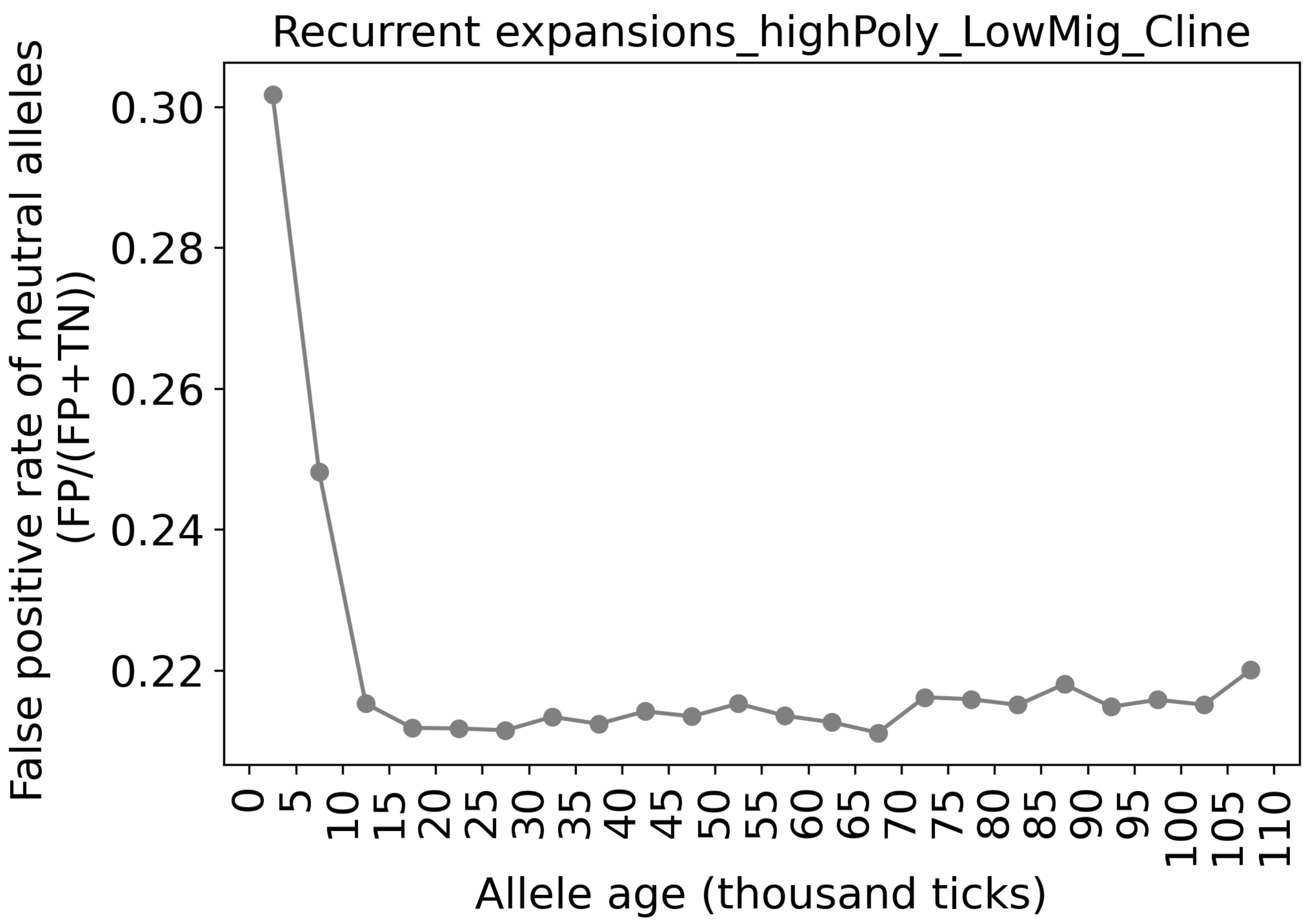
Recurrent expansions_highPoly_LowMig_Cline
False positive rate of neutral alleles (FP/(FP+TN))
0.30
0.28
0.26
0.24
0.22
0 5 10 15 20 25 30 35 40 45 50 55 60 65 70 75 80 85 90 95 100 105 110
Allele age (thousand ticks)

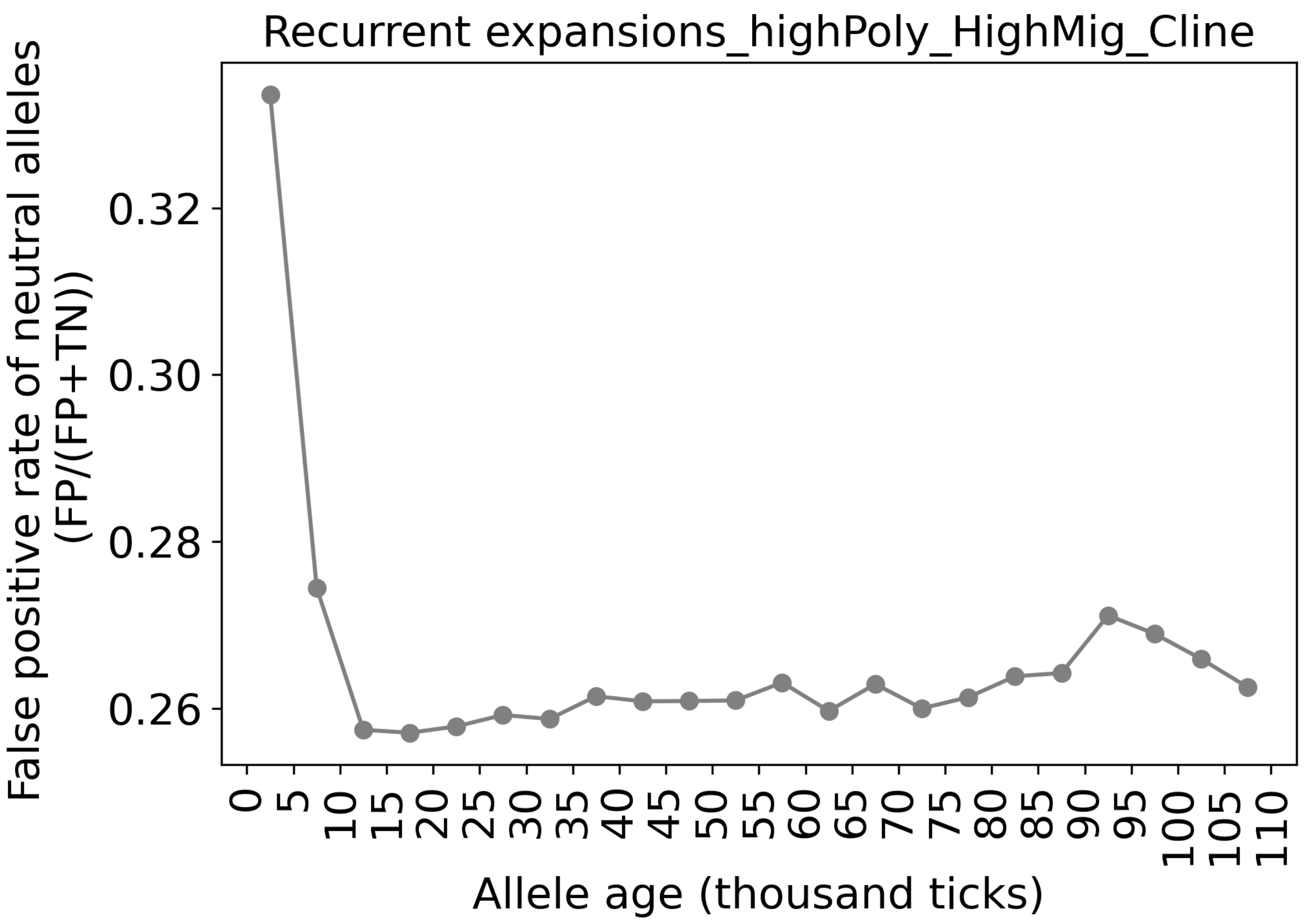
Recurrent expansions_highPoly_HighMig_Cline
False positive rate of neutral alleles (FP/(FP+TN))
0.32
0.30
0.28
0.26
0 5 10 15 20 25 30 35 40 45 50 55 60 65 70 75 80 85 90 95 100 105 110
Allele age (thousand ticks)

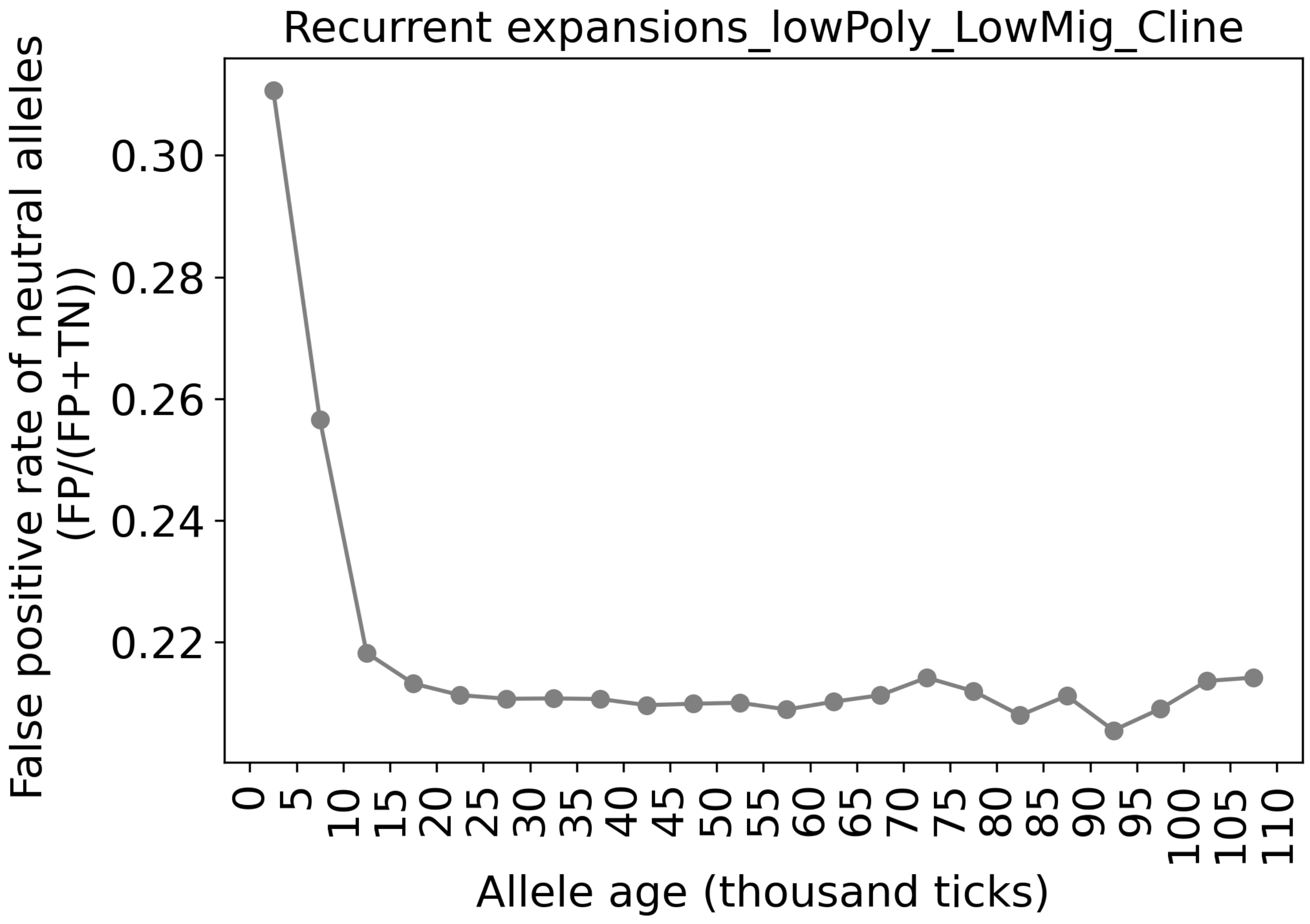
Recurrent expansions_lowPoly_LowMig_Cline
False positive rate of neutral alleles (FP/(FP+TN))
0.30
0.28
0.26
0.24
0.22
0 5 10 15 20 25 30 35 40 45 50 55 60 65 70 75 80 85 90 95 100 105 110
Allele age (thousand ticks)

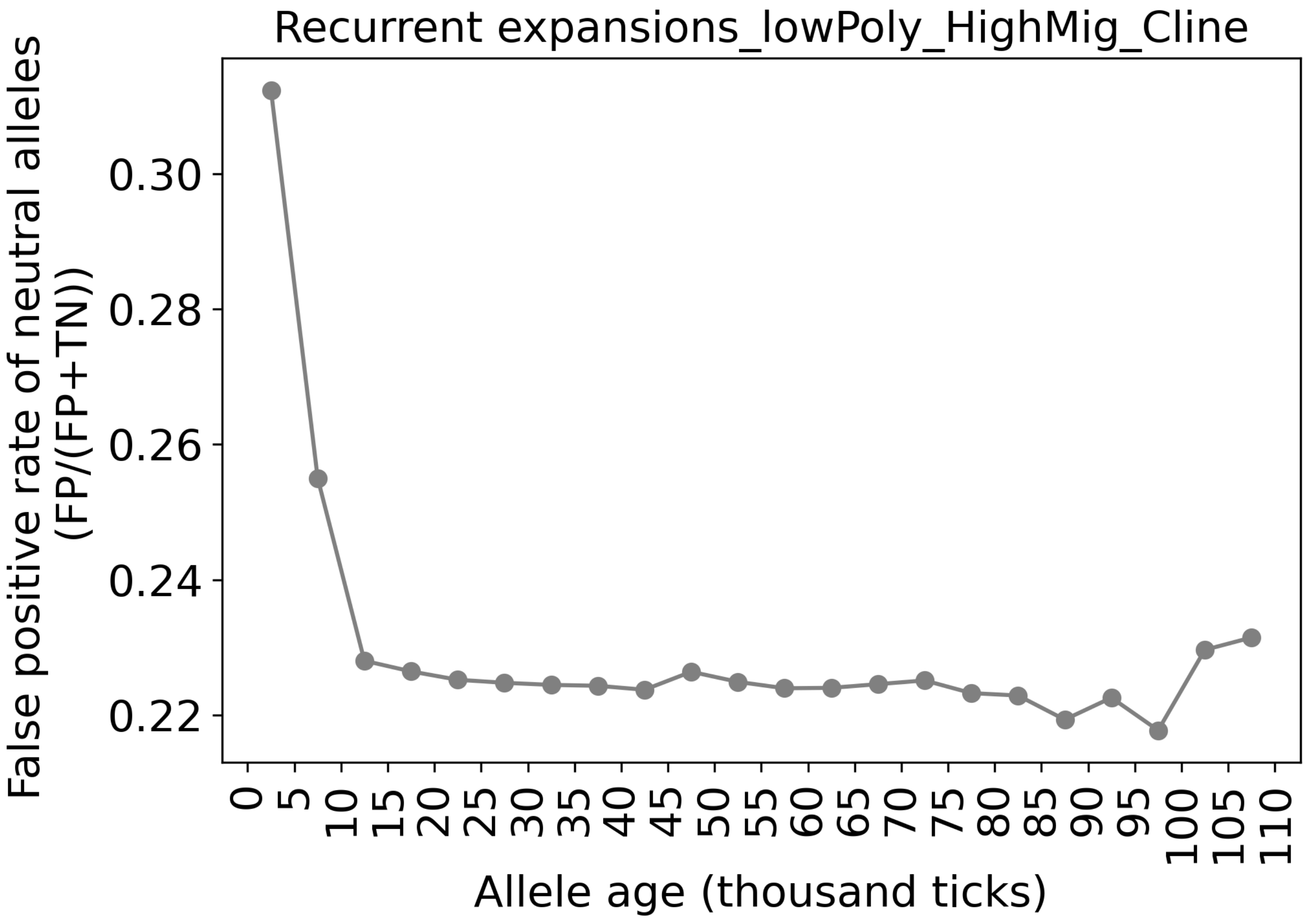
Recurrent expansions_lowPoly_HighMig_Cline
False positive rate of neutral alleles (FP/(FP+TN))
0.30
0.28
0.26
0.24
0.22
0 5 10 15 20 25 30 35 40 45 50 55 60 65 70 75 80 85 90 95 100 105 110
Allele age (thousand ticks)

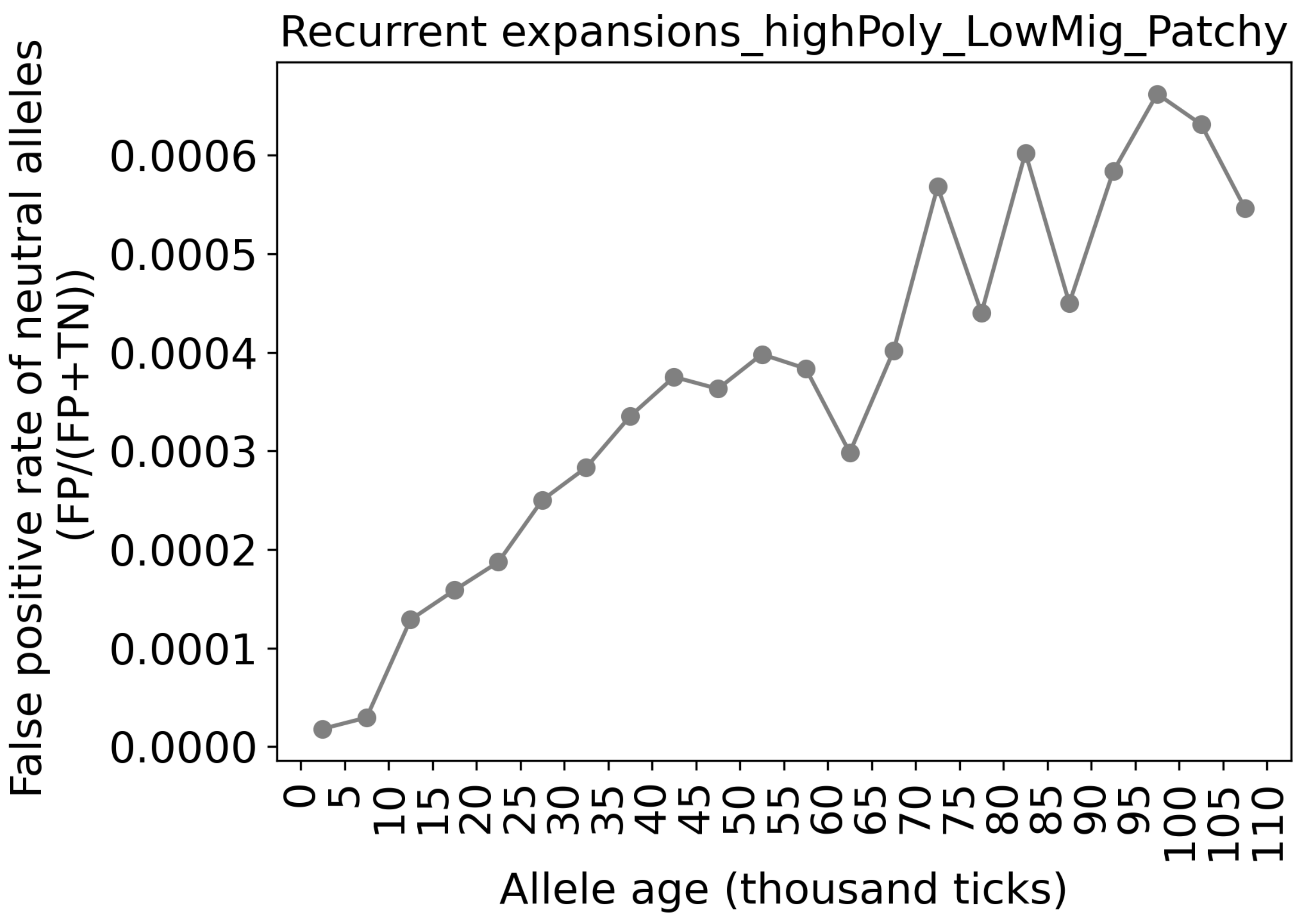
Recurrent expansions_highPoly_LowMig_Patchy
False positive rate of neutral alleles (FP/(FP+TN))
Allele age (thousand ticks)

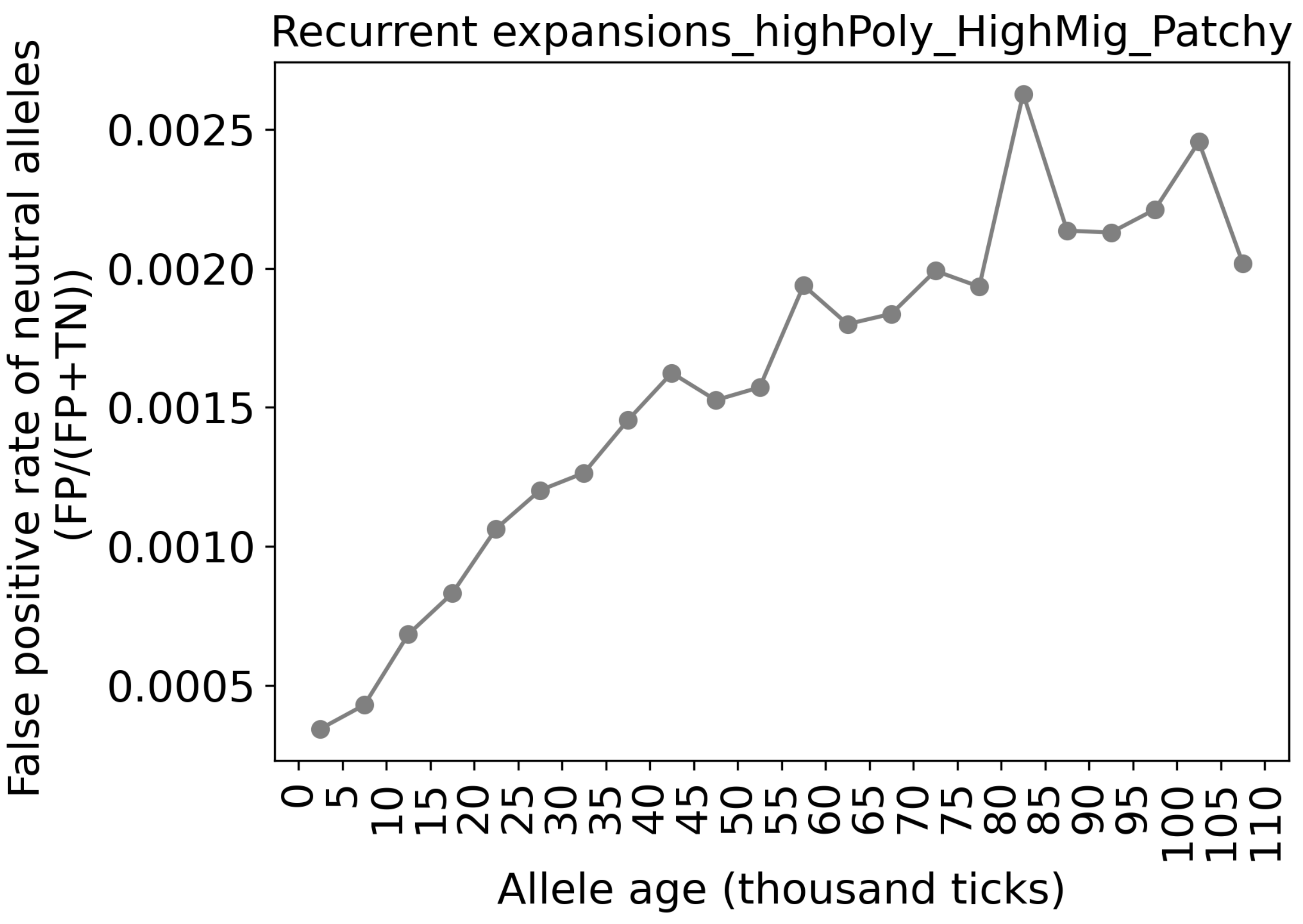
Recurrent expansions_highPoly_HighMig_Patchy
False positive rate of neutral alleles (FP/(FP+TN))
Allele age (thousand ticks)

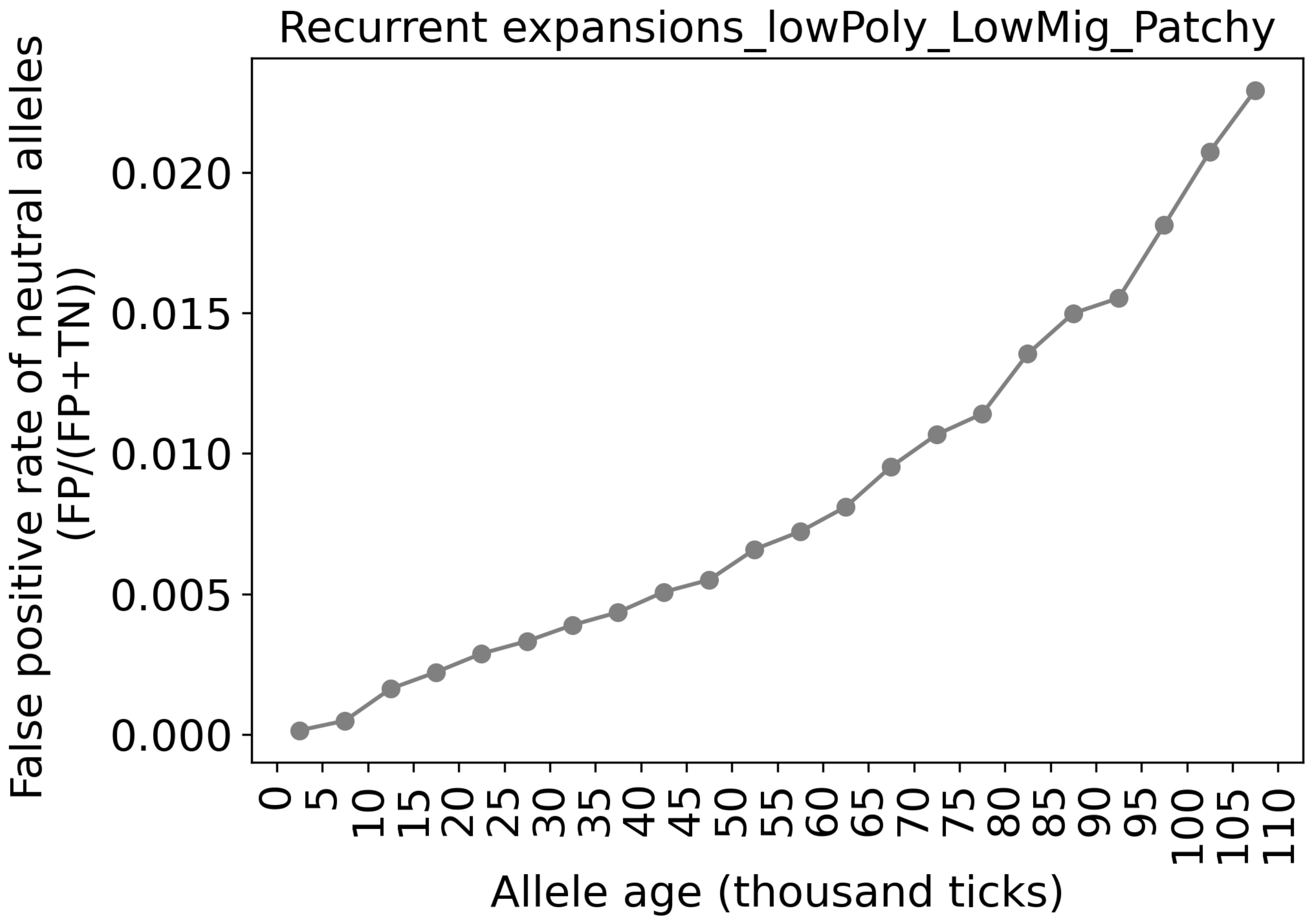
Recurrent expansions_lowPoly_LowMig_Patchy
False positive rate of neutral alleles (FP/(FP+TN))
Allele age (thousand ticks)

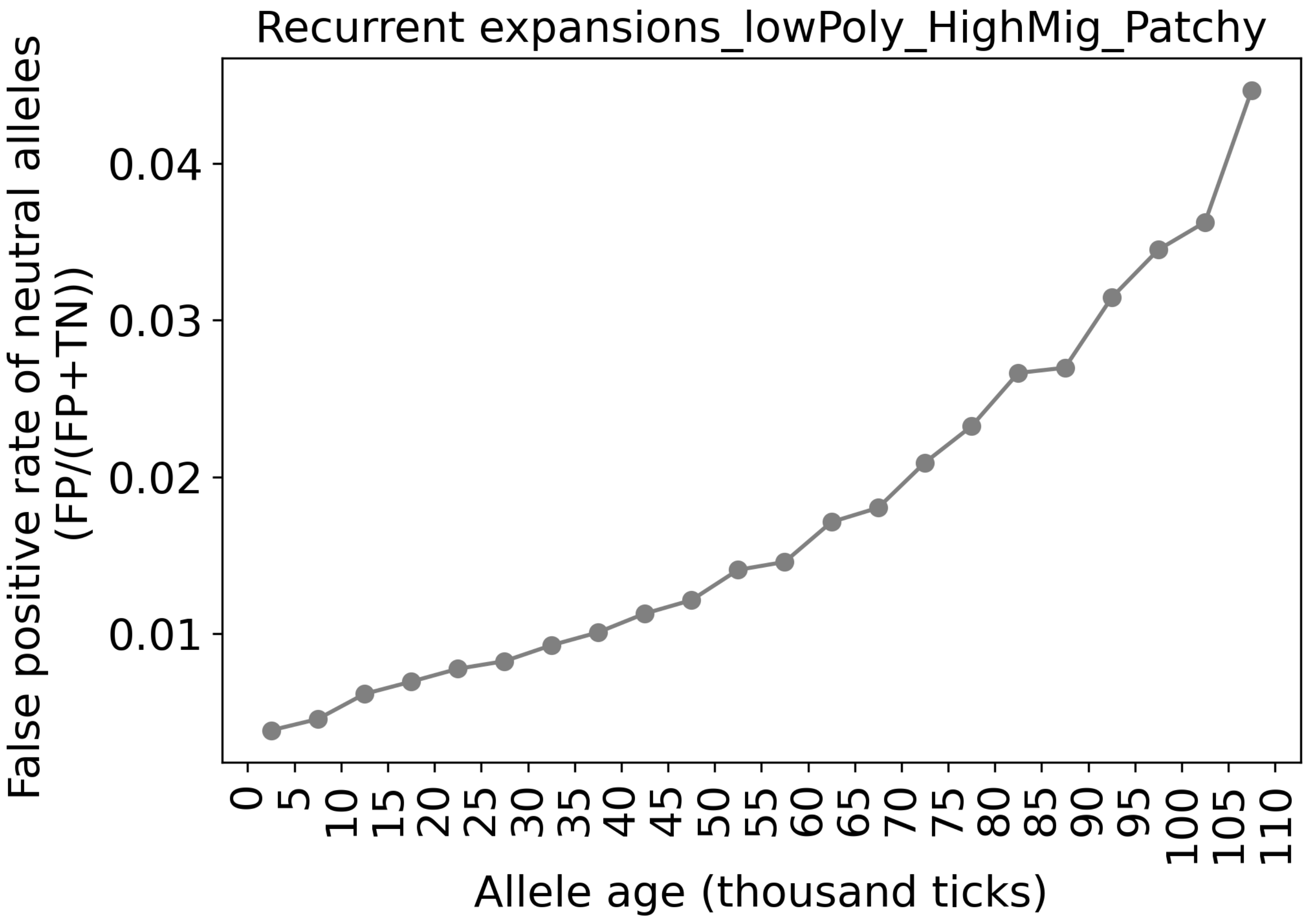
Recurrent expansions_lowPoly_HighMig_Patchy
False positive rate of neutral alleles (FP/(FP+TN))
Allele age (thousand ticks)
