## Supplementary Figure S8 for "The genetic architecture of local adaptation is historically contingent"

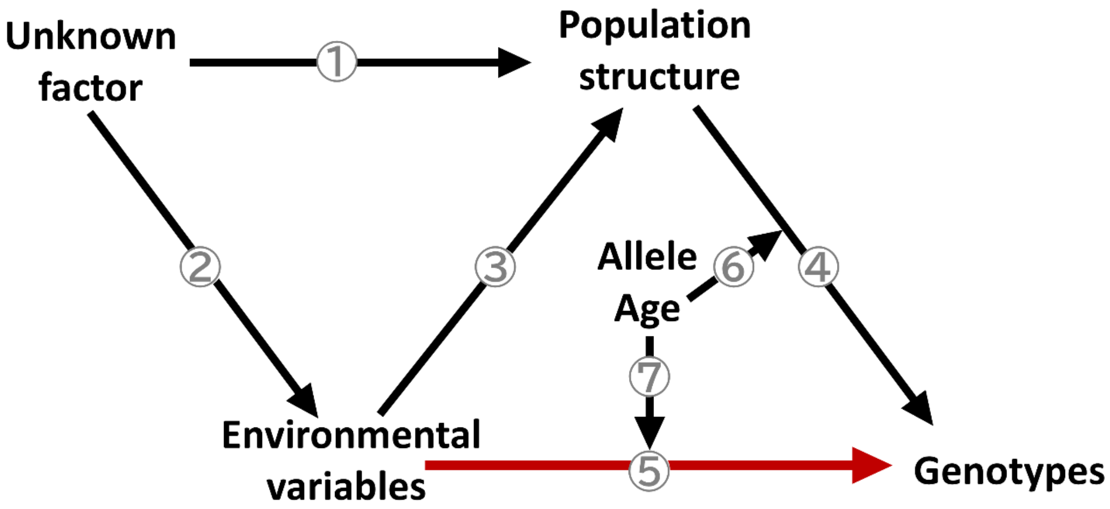


**Supplementary figure S8.** Hypothesized causal relationships among variables in genotype environment association studies (GEA), including environmental variables (E), genotypes (G), population structure, allele age, and an unknown confounding factor (U). The red arrow: Target causal effects of GEA; Black arrows: Other presumed causal effects. Arrows 1,2: The unknown confounding factor U generates a non-causal association between environmental variables and population structure, e.g., both isolation by distance and autocorrelation of environmental factors depend on the space of a surface. Arrow 3: Environmental variables may also directly affect population structure, e.g., environmental variables can create isolation by adaptation with linked selection, or affect the direction of the main axis of range expansion. Arrow 4: Population structure affects the genotypes (spatial pattern of allele frequency); Through this arrow, population structure mediates the confounding effect of U on G, but also partially mediates the target effects of E on G. Arrow 5: Direct effect of environmental variables on genotypes. Arrow 6: Allele age moderates the effects of population structure on genotypes. Arrow 7: Allele age also modifies the effect of E on G, as alleles of different ages have different probabilities of participating in local adaptation.
