## Supplementary Figure S9 for "The genetic architecture of local adaptation is historically contingent"

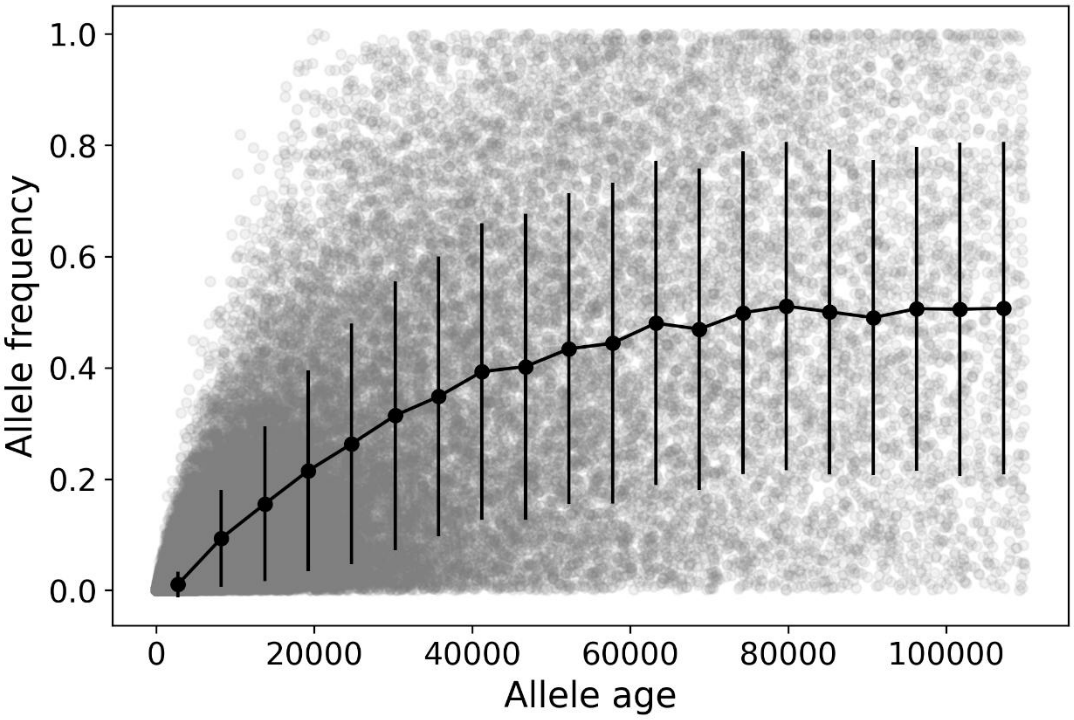


**Supplementary figure S9.** The relationship of age and frequency of 100,000 randomly sampled neutral segregating alleles (0<frequency<1) from a SLiM-simulated population (Model MNeuCon) with constant population size, no selection, and a low migration level (corresponding to average overall $F_{ST}$ values of 0.112±0.008). The average allele frequency and standard deviation of allele age in 20 equally divided time intervals are shown by black dots and error bars, respectively.
