## Supplementary Table S1-S3 for "The genetic architecture of local adaptation is historically contingent"

**Table S1** Age statistics of SLiM simulated individuals at tick 10,000

| **Model** | **Median** | **95th percentile** | **Maximum** | **Percentage of individuals older than 200 years old (%)** |
|---|---|---|---|---|
| MRecCon_HighPoly_HighMig_Cline | 4 | 23 | 1972 | 0.018 |
| MRecCon_HighPoly_HighMig_Patchy | 4 | 26 | 3583 | 0.048 |
| MRecCon_HighPoly_LowMig_Cline | 4 | 22 | 833 | 0.007 |
| MRecCon_HighPoly_LowMig_Patchy | 4 | 24 | 949 | 0.011 |
| MRecCon_LowPoly_HighMig_Cline | 4 | 23 | 903 | 0.021 |
| MRecCon_LowPoly_HighMig_Patchy | 4 | 26 | 2131 | 0.047 |
| MRecCon_LowPoly_LowMig_Cline | 4 | 22 | 1178 | 0.005 |
| MRecCon_LowPoly_LowMig_Patchy | 4 | 24 | 1365 | 0.010 |
| MRecExp_HighPoly_HighMig_Cline | 4 | 23 | 1553 | 0.021 |
| MRecExp_HighPoly_HighMig_Patchy | 4 | 26 | 3716 | 0.048 |
| MRecExp_HighPoly_LowMig_Cline | 4 | 22 | 947 | 0.008 |
| MRecExp_HighPoly_LowMig_Patchy | 4 | 24 | 651 | 0.013 |
| MRecExp_LowPoly_HighMig_Cline | 4 | 23 | 1075 | 0.019 |
| MRecExp_LowPoly_HighMig_Patchy | 4 | 26 | 2771 | 0.042 |
| MRecExp_LowPoly_LowMig_Cline | 4 | 22 | 934 | 0.007 |
| MRecExp_LowPoly_LowMig_Patchy | 4 | 23 | 939 | 0.009 |
| MSelCon_HighPoly_HighMig_Cline | 4 | 23 | 1304 | 0.023 |
| MSelCon_HighPoly_HighMig_Patchy | 4 | 26 | 1999 | 0.047 |
| MSelCon_HighPoly_LowMig_Cline | 4 | 22 | 618 | 0.008 |
| MSelCon_HighPoly_LowMig_Patchy | 4 | 24 | 927 | 0.010 |
| MSelCon_LowPoly_HighMig_Cline | 4 | 23 | 1134 | 0.019 |
| MSelCon_LowPoly_HighMig_Patchy | 4 | 26 | 3462 | 0.046 |
| MSelCon_LowPoly_LowMig_Cline | 4 | 22 | 770 | 0.005 |
| MSelCon_LowPoly_LowMig_Patchy | 4 | 24 | 742 | 0.013 |
| MSelExp_HighPoly_HighMig_Cline | 4 | 23 | 1996 | 0.021 |
| MSelExp_HighPoly_HighMig_Patchy | 4 | 26 | 2936 | 0.044 |
| MSelExp_HighPoly_LowMig_Cline | 4 | 22 | 799 | 0.007 |
| MSelExp_HighPoly_LowMig_Patchy | 4 | 24 | 918 | 0.011 |
| MSelExp_LowPoly_HighMig_Cline | 4 | 23 | 1101 | 0.017 |
| MSelExp_LowPoly_HighMig_Patchy | 4 | 26 | 2213 | 0.047 |
| MSelExp_LowPoly_LowMig_Cline | 4 | 22 | 690 | 0.007 |
| MSelExp_LowPoly_LowMig_Patchy | 4 | 24 | 941 | 0.012 |

**Table S2** $F_{ST}$ and the average mean fitness difference between local and foreign individuals ($\overline{LF}$) 10,000 simulated years after the last environmental change from 200 SLiM simulation replicates

| SLiM Model | $F_{ST}$ | | $\overline{LF}$ | |
|---|---|---|---|---|
| | mean | SD | mean | SD |
| MSelCon_HighPoly_HighMig_Cline | 0.0372 | 0.0012 | 0.1824 | 0.0013 |
| MSelCon_HighPoly_LowMig_Cline | 0.1031 | 0.0026 | 0.1874 | 0.0014 |
| MSelCon_LowPoly_HighMig_Cline | 0.0387 | 0.0015 | 0.1851 | 0.0015 |
| MSelCon_LowPoly_LowMig_Cline | 0.1079 | 0.0034 | 0.1892 | 0.0013 |
| MSelExp_HighPoly_HighMig_Cline | 0.0375 | 0.0012 | 0.1828 | 0.0015 |
| MSelExp_HighPoly_LowMig_Cline | 0.102 | 0.0026 | 0.1878 | 0.0014 |
| MSelExp_LowPoly_HighMig_Cline | 0.0374 | 0.0014 | 0.1853 | 0.0016 |
| MSelExp_LowPoly_LowMig_Cline | 0.1041 | 0.0033 | 0.1892 | 0.0015 |
| MRecCon_HighPoly_HighMig_Cline | 0.0364 | 0.0011 | 0.1823 | 0.0013 |
| MRecCon_HighPoly_LowMig_Cline | 0.1026 | 0.0025 | 0.1872 | 0.0013 |
| MRecCon_LowPoly_HighMig_Cline | 0.041 | 0.0017 | 0.1846 | 0.0015 |
| MRecCon_LowPoly_LowMig_Cline | 0.1098 | 0.0033 | 0.1888 | 0.0014 |
| MRecExp_HighPoly_HighMig_Cline | 0.0374 | 0.0011 | 0.1826 | 0.0014 |
| MRecExp_HighPoly_LowMig_Cline | 0.103 | 0.0024 | 0.1875 | 0.0013 |
| MRecExp_LowPoly_HighMig_Cline | 0.039 | 0.0016 | 0.185 | 0.0014 |
| MRecExp_LowPoly_LowMig_Cline | 0.1071 | 0.0031 | 0.1891 | 0.0015 |
| MSelCon_HighPoly_HighMig_Patchy | 0.0371 | 0.0012 | 0.0289 | 0.0008 |
| MSelCon_HighPoly_LowMig_Patchy | 0.1179 | 0.0027 | 0.033 | 0.0008 |
| MSelCon_LowPoly_HighMig_Patchy | 0.0378 | 0.0012 | 0.0303 | 0.0011 |
| MSelCon_LowPoly_LowMig_Patchy | 0.1214 | 0.0037 | 0.0344 | 0.0009 |
| MSelExp_HighPoly_HighMig_Patchy | 0.0368 | 0.0012 | 0.0288 | 0.0008 |
| MSelExp_HighPoly_LowMig_Patchy | 0.1155 | 0.0027 | 0.033 | 0.0008 |
| MSelExp_LowPoly_HighMig_Patchy | 0.0372 | 0.0013 | 0.0305 | 0.0012 |
| MSelExp_LowPoly_LowMig_Patchy | 0.1166 | 0.0033 | 0.0345 | 0.0009 |
| MRecCon_HighPoly_HighMig_Patchy | 0.0366 | 0.001 | 0.0291 | 0.0007 |
| MRecCon_HighPoly_LowMig_Patchy | 0.1166 | 0.0029 | 0.033 | 0.0008 |
| MRecCon_LowPoly_HighMig_Patchy | 0.0377 | 0.0012 | 0.0306 | 0.0011 |
| MRecCon_LowPoly_LowMig_Patchy | 0.1204 | 0.0028 | 0.0345 | 0.0009 |
| MRecExp_HighPoly_HighMig_Patchy | 0.0368 | 0.0011 | 0.0291 | 0.0008 |
| MRecExp_HighPoly_LowMig_Patchy | 0.1165 | 0.0028 | 0.0332 | 0.0007 |
| MRecExp_LowPoly_HighMig_Patchy | 0.0384 | 0.0013 | 0.0307 | 0.0011 |
| MRecExp_LowPoly_LowMig_Patchy | 0.1205 | 0.0036 | 0.0346 | 0.0009 |

**Table S3** Linear regression results for $K_{80}$ versus time (in units of 1,000 simulated years) and the type of $K_{80}$ (net/positive) 1,000-10,000 simulated years after the last environmental change in simulation models with high polygenicity and population expansions. $\beta_1$ is the slope of $K_{80}(net)$ , while $(\beta_1 + \beta_3)$ is the slope of $K_{80}(positive)$.

| **Linear model:** $\boldsymbol{K_{80}} = \beta_0 + \beta_1 Time + \beta_2 Type_{[positive]} + \beta_3 Time{:}Type_{[positive]}$ | | | | | | | | | | |
|---|---|---|---|---|---|---|---|---|---|---|
| SLiM models | | | *Time* | | | | $Time{:}Type_{[positive]}$ interaction | | | |
| Population history | Migration level | Map | $\boldsymbol{\beta_1}$ | **std err** | $\boldsymbol{t_{356}}$ | **p** | $\boldsymbol{\beta_3}$ | **std err** | $\boldsymbol{t_{356}}$ | **p** |
| Single expansion (MSelExp) | high | cline | 4.323 | 0.202 | 21.350 | <0.001 | 3.997 | 0.286 | 13.960 | <0.001 |
| | | patchy | 3.640 | 0.244 | 14.907 | <0.001 | 6.438 | 0.345 | 18.643 | <0.001 |
| | low | cline | 3.230 | 0.205 | 15.746 | <0.001 | 5.604 | 0.290 | 19.318 | <0.001 |
| | | patchy | 3.215 | 0.227 | 14.151 | <0.001 | 8.317 | 0.321 | 25.885 | <0.001 |
| Recurrent expansions and contractions (MRecExp) | high | cline | 3.287 | 0.207 | 15.878 | <0.001 | 3.986 | 0.293 | 13.615 | <0.001 |
| | | patchy | 2.863 | 0.234 | 12.231 | <0.001 | 5.410 | 0.331 | 16.346 | <0.001 |
| | low | cline | 2.421 | 0.205 | 11.811 | <0.001 | 5.486 | 0.290 | 18.925 | <0.001 |
| | | patchy | 2.678 | 0.223 | 12.032 | <0.001 | 6.790 | 0.315 | 21.570 | <0.001 |
